# Extensive mutual influences of SMC complexes shape 3D genome folding

**DOI:** 10.1101/2024.07.31.606012

**Authors:** Han Zhao, Lirong Shu, Fuhai Liu, En Lin, Sijian Xia, Baiyue Wang, Manzhu Wang, Fengnian Shan, Yinzhi Lin, Lin Zhang, Yufei Gu, Gerd A. Blobel, Haoyue Zhang

## Abstract

Mammalian genomes are folded by the distinct actions of SMC complexes which include the chromatin loop-extruding cohesin, the sister-chromatid cohesive cohesin, and the mitotic chromosome-associated condensins. While these complexes function at different stages of the cell cycle, they co-exist on chromatin during the G2/M-phase transition, when genome structure undergoes a dramatic reorganization. Yet, how distinct SMC complexes affect each other and how their mutual interplay orchestrates the dynamic folding of 3D genome remains elusive. Here, we engineered all possible cohesin/condensin configurations on mitotic chromosomes to delineate the concerted, mutual influential action of SMC complexes. We find that: (i) The mitotic SMC complex condensin disrupts the focal accumulation of extrusive-cohesin at CTCF binding sites, thereby promoting the disassembly of interphase TADs and chromatin loops during mitotic progression. Conversely, extrusive-cohesin can impair condensin activity and alter mitotic chromosome helicity. (ii) Condensin diminishes cohesive-cohesin focal enrichment and, conversely, cohesive-cohesin can counteract condensin function and impede mitotic chromosome longitudinal shortening. (iii) The co-presence of extrusive- and cohesive-cohesin synergistically antagonizes condensin function and dramatically delays mitotic chromosome condensation. (iv) Extrusive-cohesin positions cohesive-cohesin at CTCF binding sites. However, cohesive-cohesin by itself is insufficient to mediate the formation of TADs or chromatin loop, implying non-overlapping function with extrusive-cohesin. Instead, cohesive-cohesin restricts chromatin loop expansion, potentially by limiting extrusive-cohesin movement. Collectively, our data describe a comprehensive three-way interplay among major SMC complexes that dynamically sculpts chromatin architecture during cell cycle progression.

## Introduction

The structural maintenance of chromosomes (SMC) complexes including cohesin and condensin play vital roles orchestrating the dynamic folding patterns of eukaryotic genomes throughout the cell cycle ^1,2^. In G1-phase, cohesin organizes genome structure via loop extrusion, a process wherein cohesin molecules translocate the DNA fiber, progressively extruding chromatin loops until stalled at convergent CTCF binding sites ^3,4^. The extrusive form of cohesin packages the genome into topologically associating domains (TADs) and chromatin loops ^5–9^. By maintaining these structures, extrusive-cohesin can facilitate or insulate contacts between gene promoters and distal enhancers, thereby influencing transcription ^10^. During S-phase, a distinct set of cohesin complexes emerges to topologically encircle replicated DNA strands, thereby mediating cohesion between sister-chromatids ^11–14^. This cohesive form of cohesin may appear by *de-novo* loading during or after DNA replication or through conversion of pre-existing chromatin-associated cohesin molecules ^15–17^. In G2-phase, extrusive- and cohesive-cohesin complexes co-exist on the chromatin, raising the possibility that they encounter each other. It was recently reported that sister-chromatid cohesion may hinder chromatin loop expansion, indicating a potential interplay between the two forms of cohesin complexes ^18^. However, how cohesive-cohesin is distributed on the genome is largely unexplored. Furthermore, the contribution of cohesive-cohesin to the folding of chromatin fiber remains poorly understood. Finally, it is unclear whether and how cohesive-cohesin influences extrusive-cohesin-mediated structures such as TADs and chromatin loops in mammalian cells.

As cells enter mitosis, both extrusive- and cohesive-cohesin are progressively displaced from chromosome arms ^19,20^. Meanwhile, condensin is gradually loaded and begins to extrude DNA loops in a sequence-independent manner ^21–23^. This in turn compacts the interphase chromatin into condensed mitotic chromosomes ^23,24^. At the time of the condensin/cohesin exchange, there is a window during which these complexes co-exist on chromatin and influence each other. Indeed, condensin has been implicated in breaking up sister-chromatid cohesion during mitotic progression^25^. Additionally, condensin facilitates the removal of cohesin from chromosome arms in meiotic yeast cells ^26^. These findings suggest that condensin influences cohesive-cohesin. Yet, it is unknown whether condensin also modulates the distribution and functionality of extrusive-cohesin. Furthermore, it remains to be determined whether extrusive- or cohesive-cohesin exert a reciprocal influence on condensin’s loop extrusion activity to modulate mitotic chromosome condensation.

When different SMC complexes encounter each other, various outcomes may occur: stalling at the site of collision, mutual bypass, co-translocation along the DNA fiber, or eviction from the chromatin. Here, we aimed to determine the precise rules of engagement among extrusive-cohesin, cohesive-cohesin and condensin and interrogate their independent as well as combinatorial contributions to 3D genome folding. Experimental distinction of cohesive-from extrusive-cohesin during interphase is challenging. Therefore, we leveraged mitotic chromosomes as a unified experimental system that has the capacity to accommodate all three types of SMC complexes either simultaneously or independently under defined conditions. We describe a series of doubly and triply-inducible degron cell lines, which yielded mitotic genomes with eight distinct SMC complex configurations encompassing all combinations of extrusive-cohesin, cohesive-cohesin and condensin. Our study uncovered (1) a reciprocal influence between condensin and extrusive-cohesin and revealed how such interplay contributes to the transformation of interphase chromatin into mitotic chromosomes; (2) an inhibitory role of cohesive-cohesin on condensin function and mitotic chromosome condensation; (3) a positional dependence of cohesive-cohesin on extrusive-cohesin and (4) the impact of cohesive-cohesin on extrusive-cohesin mediated structures such as TADs and chromatin loops. Our data highlight a triad of mutual functional interplay among these SMC complexes that shape chromatin organization during G2-/M-phase transition.

## Results

### Condensin disrupts cohesin focal binding

Depletion of the cohesin releasing factor Wapl leads to the co-presence of cohesin and condensin on mitotic chromosomes ^27^. This perturbation enables the dissection of the “engagement rules” between condensin and cohesin. We targeted Wapl and the condensin core subunit SMC2. Using CRISPR/Cas9 directed genome editing, we generated a compound homozygous G1E-ER4 erythroblast cell line carrying a proteolysis targeting chimera (PROTAC) FKBP12^F36V^ degron (dTag)^28^ at the *Wapl* locus and a minimal auxin-inducible degron (mAID)^29^ at the *Smc2* locus respectively (Wapl^dTag^/SMC2^mAID^) (Extended Data Fig. 1a-d). The orthogonality of the dTag and AID system enabled independent degradation of Wapl and SMC2 (Extended Data Fig. 1e, f) ^30^.

Wapl depletion resulted in (1) extensive presence of Rad21 (cohesin subunit) on mitotic chromosomes (Extended Data Fig. 1g, h), (2) a high degree (95.5±1.5%) of “closed” chromatid arms in metaphase chromosome spreads (Extended Data Fig. 1i), (3) defective chromosome segregation in ana/telophase (Extended Data Fig. 1j, k) and (4) significantly delayed cell growth (Extended Data Fig. 1l). Taken together, these data demonstrate that Wapl depletion led to mitotic retention of cohesin. Importantly, cohesin retention did not affect the occupancy of condensin on mitotic chromosomes (Extended Data Fig. 1g), thereby allowing us to study their interplay.

To illustrate the influence of condensin on cohesin, we began by characterizing the baseline binding profile of mitotically bound cohesin, independent of any potential influence from condensin. We exploited the fact that CTCF was partially retained on mitotic chromosomes in G1E-ER4 cells ^20^ and posit that if cohesin preserved loop extrusion activity during mitosis, it would be halted by these mitotically bound CTCF molecules. We treated the Wapl^dTag^/SMC2^mAID^ cells with dTag-13 and 5-Ph-IAA for 4 hours followed by nocodazole mediated prometaphase arrest (Extended Data Fig. 2a Upper panel). Mitotic cells co-depleted of Wapl and SMC2 were then purified through fluorescence activated cell sorting (FACS) (Extended Data Fig. 2b, c Upper panel). We then performed chromatin immunoprecipitation and sequencing (ChIP-seq) against Rad21. We identified 20,508 Rad21 peaks, majority (16,866, ∼82.2%) of which also appeared in interphase, suggesting a common mechanism that governs cohesin positioning (Extended Data Fig. 3a, b). Mitotically preserved Rad21 peaks coincided with stronger CTCF retention compared to those that were lost, implying a role for residual CTCF for cohesin anchoring (Extended Data Fig. 3c). The overall Rad21 ChIP-seq signal intensity was weaker in mitosis compared to interphase, reflecting mitotic eviction of CTCF (Extended Data Fig. 3b, d). Interestingly, we also identified 3,642 Rad21 peaks that were exclusively present in mitosis but absent in interphase cells (Extended Data Fig. 3a, b, d). These peaks lacked CTCF binding motif, were significantly broader in size and frequently co-localized with *cis*-regulatory elements (CREs), which were marked by high levels of active histone marks (Extended Data Fig. 3e-h). This finding aligns with a prior report that cohesin relocates to CREs upon CTCF depletion ^31^. Together, our data indicate that cohesin can be positioned at both residual CTCF binding sites and CREs, suggesting a capacity of loop extrusion in the condensin-deficient mitotic chromosomes.

To investigate whether condensin affects cohesin positioning, we FACS purified Wapl-deficient mitotic cells without 5-Ph-IAA treatment such that condensin co-existed with cohesin on mitotic chromosomes (Extended Data Fig. 2a-c lower panel). ChIP-seq experiments revealed that condensin extensively diminished cohesin focal binding, showing a ∼47.7% (9,783) reduction in detectable Rad21 peaks, with the remaining peaks (10,725) exhibiting a drastic decrease in signal intensity (Extended Data Fig. 4a, b). To further dissect the impact of condensin on cohesin, we subcategorized mitotic Rad21 peaks based on their genomic co-localization: (1) with CTCF (n=14,753), (2) with CREs (n=2,923) and (3) with both CTCF and CREs (n=1,133). We found that condensin significantly attenuated Rad21 peak strength at CTCF binding sites (Extended Data Fig. 4c, d). Importantly, CTCF focal enrichment was not significantly affected, indicating that condensin mediated disruption of cohesin peaks was not caused by direct displacement of CTCF (Extended Data Fig. 4c, d). Furthermore, the CRE and CTCF/CRE co-occupied Rad21 peaks were also dramatically reduced upon condensin loading (Extended Data Fig. 4c, e). Collectively, these findings suggest that condensin broadly impedes the focal accumulation of cohesin.

### Condensin disrupts extrusive-cohesin focal enrichment

We next wondered whether condensin exerts similar or distinct influences on extrusive- and cohesive-cohesin. To address this, we sought to selectively preserve extrusive-cohesin on mitotic chromosomes while ablating cohesive-cohesin. We targeted Sororin, a key factor that safeguards cohesive-cohesin from Wapl mediated release following DNA replication ^32,33^. We engineered an orthogonal dual inducible degron cell line Wapl^dTag^/Sororin^mAID^, which allowed rapid degradation of the two target proteins within 4 hours through dTag13 and 5-Ph-IAA co-treatment (Fig. 1a; Extended Data Fig. 5a-c). Consistent with previous reports, degradation of Sororin resulted in a near-complete dissociation of sister-chromatids ^33^, signifying the absence of cohesive-cohesin (Fig. 1b, c). However, concurrent Wapl depletion, a step necessary for retaining extrusive-cohesin, abrogated this effect of Sororin loss, thereby hindering the separation of the two cohesin forms (Fig. 1b, c). To resolve this issue, we implemented a multi-step G2/M arrest and release protocol. First, asynchronous Wapl^dTag^/Sororin^mAID^ cells were treated with 5-Ph-IAA to depleted Sororin and arrested at the G2/M transition using the CDK1 specific inhibitor RO-3306 (Fig. 1d upper panel). G2/M arrest in the absence of Sororin enabled Wapl to evict the unprotected cohesive-cohesin, while leaving extrusive-cohesin unaffected. The extent of cohesive-cohesin removal was positively correlated with the duration of G2/M arrest and a 16-hour arrest in G2 led to complete separation of sister-chromatids in ∼80% of cells (Extended Data Fig. 5d-f). Subsequently, the cells were incubated with dTag13 for 4 hours to promote Wapl degradation, thus allowing extrusive-cohesin to retain after mitotic entry (Fig. 1d upper panel). Finally, RO-3306 was washed out to permit mitotic progression and nocodazole was applied to enrich prometaphase cells predominantly harboring extrusive-but not cohesive-cohesin (Fig. 1d upper panel).

**Figure 1.**
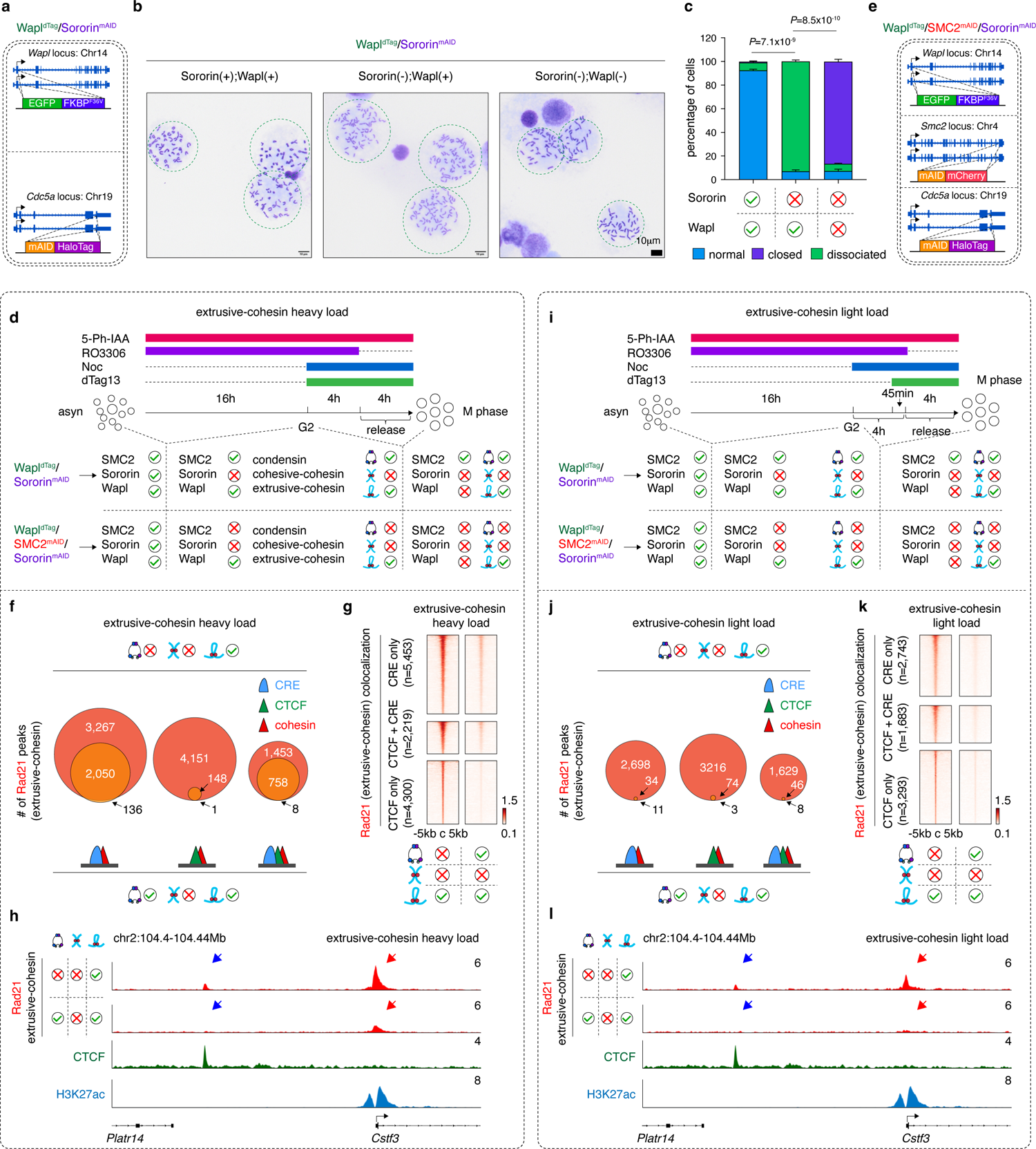
Condensin disrupts extrusive-cohesin focal binding. **a**, Schematic showing the genome editing strategy to generate the Wapl^dTag^/Sororin^mAID^ cell line. **b**, Metaphase spread analysis showing a complete loss of sister-chromatid cohesion in Sororin depleted mitotic Wapl^dTag^/ Sororin^mAID^ cells. Co-depletion of Wapl and Sororin, abrogated this phenotype. Scale bar: 10μm. **c**, Quantification of (**b**) showing the percentage of sister-chromatid dissociation in indicated samples. *P* values were calculated by two-sided student’s *t*-test. **d**, Upper panel: G2/M arrest/release strategy to obtain condensin-replete mitotic chromosomes with extrusive-cohesin (“heavy load”). For simplicity, we employed three illustrative symbols to represent condensin, extrusive-cohesin and cohesive-cohesin. These symbols are used hereafter throughout the study. Lower panel: G2/M arrest/release strategy to obtain condensin-deficient mitotic chromosomes with extrusive-cohesin (“heavy load”). **e**, Schematic showing the genome editing strategy to generate the Wapl^dTag^/SMC2^mAID^/Sororin^mAID^ cell line. **f**, Venn-diagrams showing the loss of extrusive-cohesin peaks (“heavy load”) in the presence of condensin. Peaks co-localized to CTCF, CREs or both were shown separately. **g**, Density heatmap plots showing reduced mitotic extrusive-cohesin peak strength (“heavy load”) in the presence of condensin. Peaks co-localized to CTCF, CREs or both were plotted separately. **h**, Browser tracks showing that condensin disrupts extrusive-cohesin (Rad21) peaks (“heavy load”). Peaks coincided with CTCF and CRE were indicated by blue and red arrows respectively. Tracks of CTCF and H3K27ac from parent cells were shown. **i**, Similar to (**d**) showing the strategies to obtain condensin-replete or deficient mitotic chromosomes with low levels of extrusive-cohesin (“light load”). (**j**-**l**) Similar to (**f**-**h**) with Venn-diagrams (**j**), Density heatmap plots (**k**) and browser tracks (**l**) showing the complete disruption of extrusive-cohesin (“light load”) by condensin.

To interrogate the influence of condensin on extrusive-cohesin, we further engineered a tri-inducible degron cell line with an additional mAID tag fused to the endogenous SMC2 protein (Wapl^dTag^*/*SMC2^mAID^*/*Sororin^mAID^) (Fig. 1e; Extended Data Fig. 5a, g, h). Subjecting this and the above generated Wapl^dTag^/Sororin^mAID^ cell line to the G2/M arrest/release protocol enabled us to obtain condensin-deficient or replete mitotic chromosomes specifically loaded with extrusive-cohesin (Fig. 1d). In the absence of condensin, extrusive-cohesin displayed 12,755 peaks as revealed by Rad21 ChIP-seq. We also identified 382 cohesin islands with a mean size of ∼73kb that were undetectable in interphase cells (Extended Data Fig. 6a, b). This observation is consistent with a previous finding that cohesin forms islands in CTCF and Wapl double knockout cells ^31^. Upon the introduction of condensin, we observed a dramatic loss of extrusive-cohesin peaks by ∼76.4%, (9,749 peaks lost). Of note, peaks co-occupied by CTCF exhibited a more pronounced disruption by condensin (∼96.5% lost) compared to those located at CREs (∼60% lost) (Fig. 1f-h; Extended Data Fig. 7a-c). In addition, cohesin islands also vanished due to the presence of condensin (Extended Data Fig. 6b-e). In summary, our data demonstrate that condensin is capable of interrupting the focal enrichment of extrusive-cohesin on mitotic chromosomes.

We noticed that the CRE co-localized extrusive-cohesin peaks displayed higher signal intensity than those coincided with CTCF (Fig. 1g, h), suggesting a potential reverse correlation between cohesin focal abundance and the susceptibility to condensin mediated disruption. To investigate this possibility, we sought to manipulate extrusive-cohesin occupancy and evaluate subsequent alterations in the sensitivity of extrusive-cohesin peaks to condensin perturbation. We devised a strategy to reduce extrusive-cohesin level on mitotic chromosomes by shortening dTag13 treatment from 4 hours to 45 minutes before G2-release (Fig. 1i). Immunofluorescence staining uncovered weaker retention of Rad21 in samples incubated with dTag13 for 45 minutes (“light load”) compared to those receiving the full course of 4-hour dTag13 treatment (“heavy load”) (Extended Data Fig. 7d). As expected, the “light load” group exhibited fewer extrusive-cohesin peaks (n=7,821) and reduced ChIP-seq signal intensity (Extended Data Fig. 7e, f). Remarkably, the “light load” group displayed a near-complete elimination of detectable Rad21 peaks (∼97.5% peak loss) in response to condensin, irrespective of their genomic context (co-localization with CTCF, CRE or both) (Fig. 1j-l; Extended Data Fig. 7g, h).

Taken together, our data indicate that condensin completely disrupts the focal accumulation of extrusive-cohesin when it is present at low level. This disruptive effect may exhibit an upper limit, leaving residual signals when extrusive-cohesin abundance exceeds certain threshold. Our observations imply a potential role for condensin in dismantling of extrusive-cohesin focal accumulation in prophase, thereby expediting the disassembly of interphase genome folding patterns such as TADs and chromatin loops.

### Condensin interrupts cohesive-cohesin focal accumulation analogously to its disruption of extrusive-cohesin

To decipher the potential influence of condensin on cohesive-cohesin, we sought to map the binding profile of Sororin, which selectively marks cohesive-cohesin ^14,32^, in the absence versus the presence of condensin. Since Sororin is typically evicted from mitotic chromosome arms by CDK1/Aurora B mediated phosphorylation, we employed a previously characterized mutant form of Sororin, harboring nine evolutionarily conserved phosphosites mutated to alanine ^34,35^ (Extended Data Fig. 8a). The flag-tagged non-phosphorylatable Sororin^9A^ mutant was stably introduced into Wapl^dTag^*/*SMC2^mAID^*/*Sororin^mAID^ and Wapl^dTag^/Sororin^mAID^ cells via retroviral transduction. Consistent with previous findings, Sororin^9A^ exhibited extensive mitotic retention and promoted widespread sister-chromatid cohesion (Extended Data Fig. 8b, c), thereby enabling us to track cohesive-cohesin during mitosis ^35^.

To establish the basal binding pattern of cohesive-cohesin, we employed the aforementioned G2/M arrest/release protocol on Wapl^dTag^*/*SMC2^mAID^*/*Sororin^mAID^ cells expressing Sororin^9A^ and obtained condensin-deficient mitotic chromosomes co-loaded with cohesive-(mediated by Sororin^9A^) and extrusive-cohesin (Fig. 2a, b upper panel). Cells with “heavy” or “light” load of extrusive-cohesin were both harvested (Fig. 2a, b upper panel). Rad21 ChIP-seq, which recognized both cohesin forms unveiled 10,426 and 8,339 peaks in the “heavy” and “light” load groups respectively. These peaks displayed a high degree of overlap with previously identified extrusive-cohesin binding sites (Extended Data Fig. 9a). This observation suggests that cohesive-cohesin may lack the capacity to establish *de novo* peaks independently of extrusive-cohesin. Next, we employed the Sororin^9A^ expressing Wapl^dTag^*/*Sororin^mAID^ cells to introduce condensin and interrogate its influence on cohesive-cohesin (Fig. 2a, b lower panel). Importantly, condensin diminished Rad21 peaks (representing both cohesive- and extrusive-cohesin) in a manner similar to its disruption of extrusive-cohesin alone: (1) the “heavy load” group displayed a marked reduction in Rad21 ChIP-seq signal (∼64% peak loss) (Extended Data Fig. 9b-e); (2) Rad21 peaks in the “light load” group became virtually undetectable (99% loss) (Extended Data Fig. 9e-h); and (3) cohesin islands were dramatically diminished by condensin (Extended Data Fig. 6f-i). We thus conclude that the co-presence of cohesive- and extrusive-cohesin does not noticeably differ from extrusive-cohesin alone in genomic binding patterns and response to condensin.

**Figure 2.**
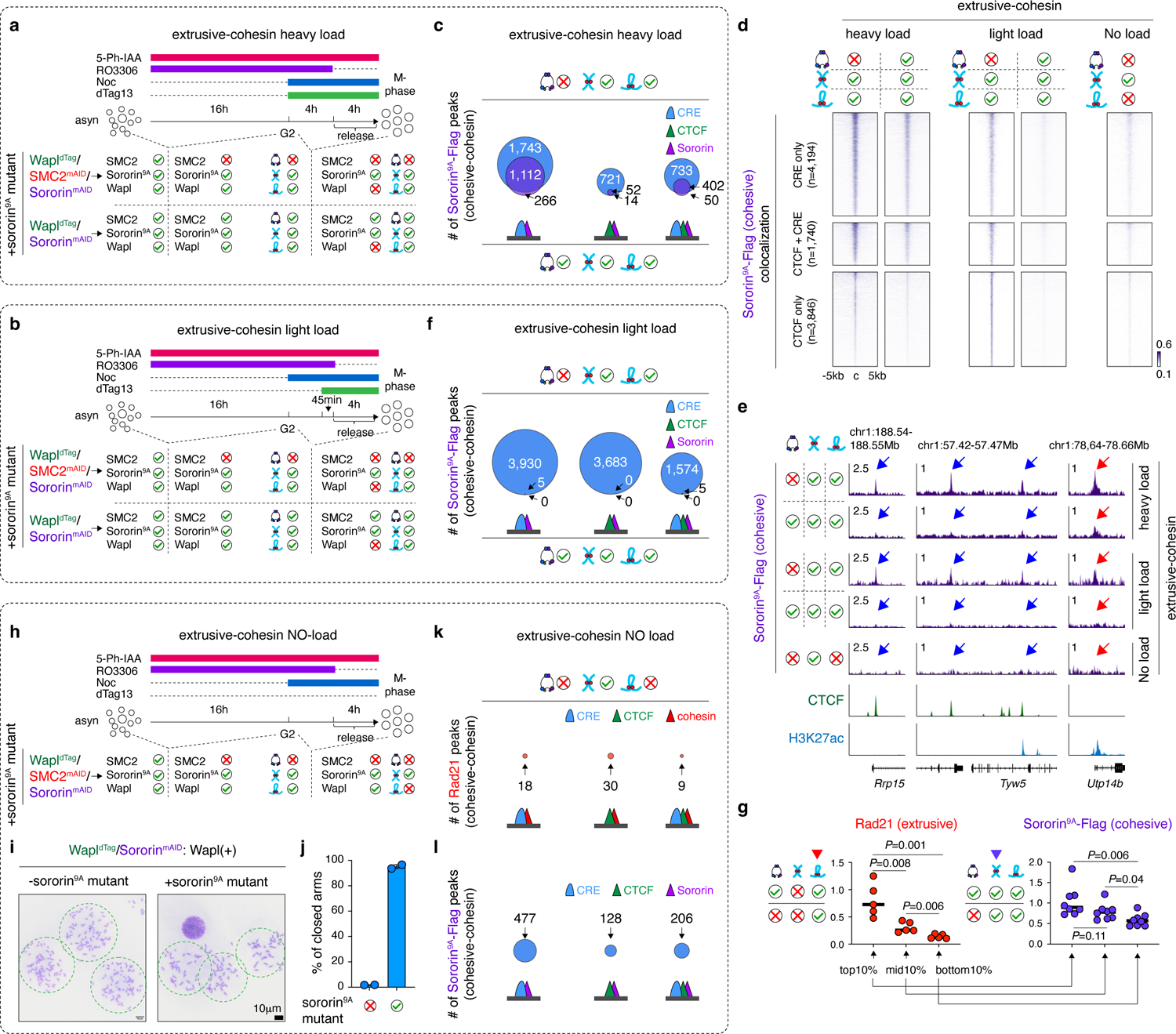
Condensin disrupts cohesive-cohesin focal binding. **a**, Upper panel: G2/M arrest/release strategy to obtain condensin-deficient mitotic chromosomes with extrusive-cohesin (“heavy load”) and cohesive-cohesin in cells expressing Sororin^9A^ mutant. Lower panel: G2/M arrest/release strategy to obtain condensin-replete mitotic chromosomes with extrusive-cohesin (“heavy load”) and cohesive-cohesin in cells expressing Sororin^9A^ mutant. **b**, Similar to (**a**) showing the strategies to obtain condensin-deficient or replete mitotic chromosomes with low levels of extrusive-cohesin (“light load”) and cohesive-cohesin in cell expressing Sororin^9A^ mutant. **c**, Venn-diagrams showing the loss of flag-tagged Sororin^9A^ peaks in the context of heavily loaded extrusive-cohesin in response to condensin. Sororin peaks co-localized to CTCF, CREs or both were shown separately. **d**, Left panel: Density heatmap plots showing reduced peak strength of Sororin^9A^ in the context of heavily loaded extrusive-cohesin in response to condensin. Middle panel: Density heatmap plots showing complete loss of peak Sororin^9A^ peak signal in the context of light loaded extrusive-cohesin in the presence of condensin. Right panel: Density heatmap plots showing minimal Sororin^9A^ peak signal in the absence of extrusive-cohesin in condensin-deficient cells. Sororin^9A^ peaks co-localized to CTCF, CREs or both were plotted separately. **e**, Browser tracks showing representative Sororin^9A^ peaks in indicated conditions. Peaks colocalized to CTCF and CRE were indicated by blue and red arrows respectively. Tracks of CTCF and H3K27ac from parental cells were shown. **f**, Similar to (**c**) showing the complete loss of flag-tagged Sororin^9A^ peaks in the context of lightly loaded extrusive-cohesin in response to condensin. **g**, Left panel: Dot plots showing the fold change of extrusive-cohesin peak signals in response to condensin. Top, middle and bottom 10% peaks that were most sensitive to condensin were shown. *P* values were computed using two-sided student’s *t*-test (n=5). Right panel, Dot plots showing the fold change of cohesive-cohesin (Sororin^9A^-flag) peak signals in regions corresponding to those in left panel. *P* values were computed using two-sided student’s *t*-test (n=8). Note that the responses of cohesive-cohesin to condensin followed that of extrusive-cohesin. **h**, Similar to (**a**) showing the strategies to obtain condensin-deficient mitotic chromosomes with cohesive-cohesin but not extrusive-cohesin in cell expressing Sororin^9A^ mutant. **i**, Metaphase spread analysis showing that Sororin^9A^ maintains sister-chromatid cohesion in the presence of Wapl. **j**, Quantification of (**i**) showing the percentage of “closed-arms” in cells expressing Sororin^9A^. **k**, Venn-diagrams showing negligible Rad21 peak calls in condensin-deficient mitotic cells with cohesive-but not extrusive-cohesin. **l**, Venn-diagrams showing low number of Sororin^9A^ peaks in condensin-deficient mitotic cells with cohesive-but not extrusive-cohesin.

Next, we performed ChIP-seq targeting the flag-tagged Sororin^9A^ mutant to directly monitor cohesive-cohesin (Fig. 2a, b). In the context of heavily loaded extrusive-cohesin, Sororin^9A^ displayed 5,205 peaks. Vast majority (∼88.9%, 4,625) of them coincided with extrusive-cohesin peaks, consolidating the notion that cohesive-cohesin may not exhibit independent binding sites. Condensin triggered a substantial reduction of detectable Sororin^9A^ peaks by ∼69.6% (3,621 peaks lost). Of note, Sororin^9A^ peaks co-occupied by CTCF exhibited a greater susceptibility to the disruptive force of condensin compared to those located at CREs (Fig. 2c, d left panel, e upper panel). In cells lightly loaded of extrusive-cohesin, we observed a more severe disruption of Sororin^9A^ peaks (>99% peak lost) by condensin (Fig. 2d middle panel, e middle panel, f). Thus, the response of cohesive-cohesin to condensin recapitulated what we observed for extrusive-cohesin (Fig. 1). To further corroborate this finding, we stratified the 12,755 extrusive-cohesin binding sites and segmented them into three tiers on the basis of their resistance to condensin: top 10% (most resistant), middle 10% and bottom 10% (most sensitive) (Fig. 2g left panel). Notably, Sororin^9A^ displayed a similar pattern of signal reduction across all three groups upon condensin loading, indicating a positive correlation between the disruption of cohesive- and extrusive-cohesin by condensin (Fig. 2g right panel). Taken together, our data suggest that condensin dislocates cohesive-cohesin in a manner that mirrors its effect on extrusive-cohesin.

### Cohesive-cohesin focal enrichment depends on extrusive-cohesin

The paralleled disruption patterns for both cohesin types by condensin implied a potential dependence of cohesive-cohesin’s focal accumulation on extrusive-cohesin. To experimentally validate this hypothesis, we sought to profile cohesive-cohesin binding patterns in the absence of extrusive-cohesin. To achieve this, we modified our G2/M arrest/release protocol by omitting dTag13 treatment, thereby preserving Wapl (Fig. 2h). Wapl mediates the removal of extrusive-cohesin during mitotic progression whereas cohesive-cohesin was protected by mitotically retained Sororin^9A^ mutant (Fig. 2i, j). We applied this protocol to Wapl^dTag^*/*SMC2^mAID^*/*Sororin^mAID^ cells expressing Sororin^9A^ and obtained condensin-deficient mitotic chromosomes selectively loaded with cohesive-cohesin. Strikingly, cohesive-cohesin alone failed to achieve identifiable focal accumulation with only 141 Rad21 peaks detected (Fig. 2k). This finding was further corroborated by ChIP-seq targeting Sororin^9A^, which revealed a dramatic reduction in peak calls compared to that observed in the presence of extrusive-cohesin (Fig. 2l). Our data indicate that cohesive-cohesin may lack the capacity of peak formation in the absence of extrusive-cohesin.

We realized that while Sororin^9A^ protects cohesive-cohesin from being released, Wapl may still gradually unload cohesive-cohesin during G2 and prometaphase arrest. This could potentially lead to reduced number of detectable peaks in our ChIP-seq experiments. To address this confounding factor, we aimed to minimize the dissociation of cohesive-cohesin from chromatin by reducing the duration of Wapl exposure. Hence, we further modified our G2/M arrest/release protocol by shortening the RO-3306 treatment to 8 hours (“short arrest”) and the final RO-3306 release to 30min (“short release”) (Extended Data Fig. 10a-c). We applied this G2/M short-arrest and release protocol to Sororin^9A^-expressing Wapl^dTag^*/*SMC2^mAID^*/*Sororin^mAID^ cells to yield condensin-deficient mitotic chromosomes with enhanced occupancy of cohesive-cohesin. Notably, Rad21 ChIP-seq only identified 604 peaks (Extended Data Fig. 10d). Finally, we performed ChIP-seq targeting flag-tagged Sororin^9A^ to directly monitor cohesive-cohesin in the “short-arrest and release” samples. We observed only 228 cohesive-cohesin peaks, a substantial reduction compared to thousands of peaks detected in the presence of extrusive-cohesin (Extended Data Fig. 10e-h). Our data thus demonstrate that loss of cohesive-mcohesin peaks is not attributable to diminished chromatin occupancy.

To this end, we conclude that cohesive-cohesin does not independently form peaks. Instead, they rely on extrusive-cohesin to achieve measurable focal accumulation. This positional dependence implies a potential mechanical interplay between cohesive- and extrusive-cohesin. It is therefore conceivable that cohesive-cohesin may influence the formation and/or maintenance of extrusive-cohesin mediated structures such TADs and chromatin loops.

### Extrusive- and cohesive-cohesin exhibit distinct, yet integrative impacts on mitotic chromosome organization

The observed disruption of cohesin by condensin prompted us to investigate the potential for a reciprocal interaction, where cohesin complexes influence condensin’s loop extrusion activity and consequently affect mitotic chromosome condensation. To address this possibility, we implemented the G2/M arrest and release protocol on the Wapl^dTag^*/*Sororin^mAID^ line in both the original form and a variant expressing the Sororin^9A^ mutant. We obtained condensin-replete mitotic chromosomes with four distinct configurations of cohesin (Fig. 3a):

1. condensin(+); extrusive-cohesin (-); cohesive-cohesin (-) (control)
2. condensin(+); extrusive-cohesin (+); cohesive-cohesin (-)
3. condensin(+); extrusive-cohesin (-); cohesive-cohesin (+, protected by Sororin^9A^)
4. condensin(+); extrusive-cohesin (+); cohesive-cohesin (+, protected by Sororin^9A^)

**Figure 3.**
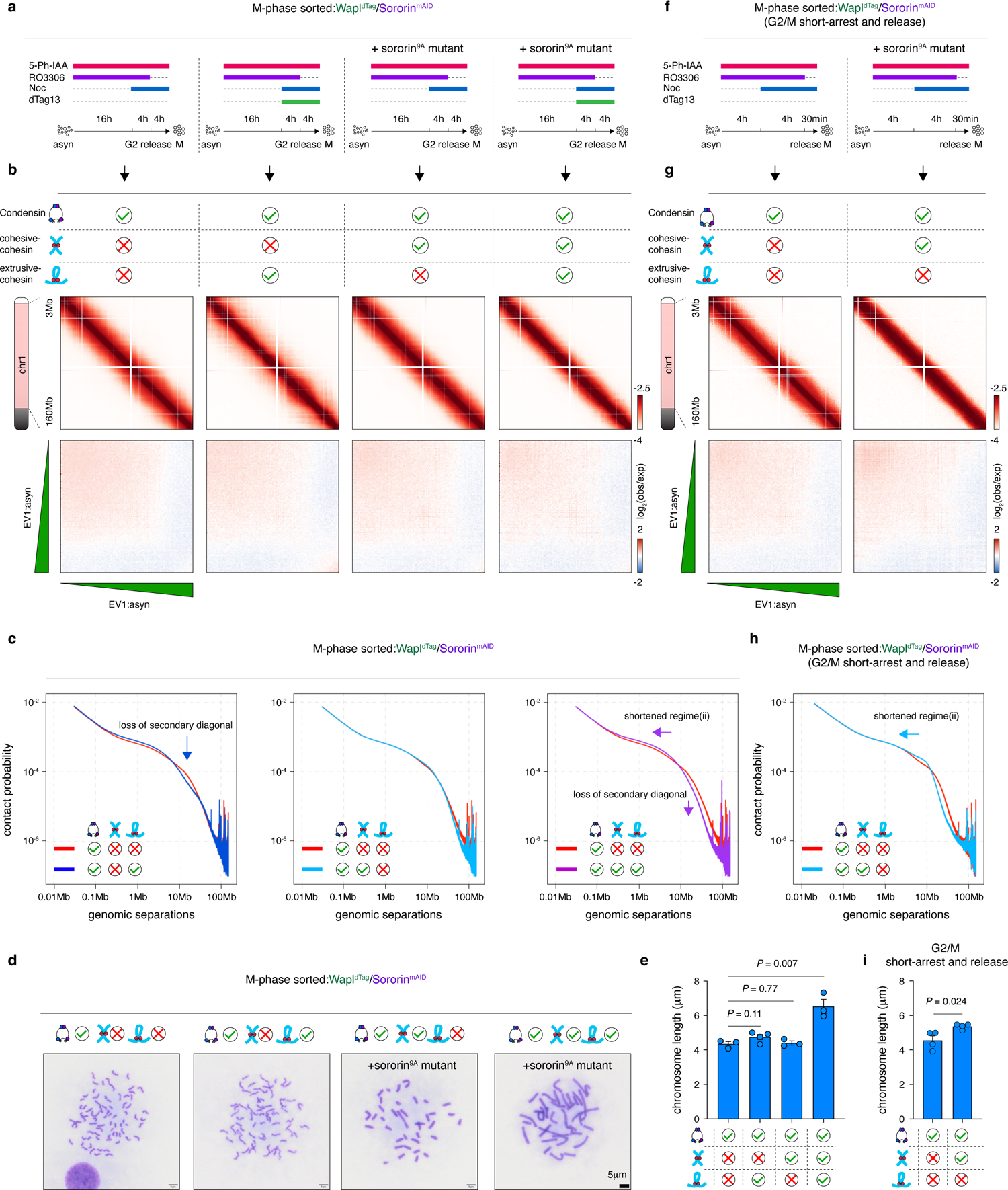
Extrusive- and cohesive-cohesin influences distinct but integrative aspects of mitotic chromosome structure. **a**, Schematic showing the G2/M arrest and release strategies on the Wapl^dTag^/Sororin^mAID^ cells to obtain condensin-replete mitotic cells with four different cohesin states (configurations:1-4). **b**, Upper panel: KR-balanced Hi-C contact maps of the mitotic Wapl^dTag^/Sororin^mAID^ cells with indicated cohesin configurations. Bin size: 100kb. Lower panel: Saddle plots showing that compartments were missing in all mitotic Wapl^dTag^/Sororin^mAID^ samples. **c**, *P(s)* curves showing the influences (indicated by arrows) of extrusive-cohesin or/and cohesive-cohesin on mitotic chromosome architecture. **d**, Metaphase spread analysis showing representative morphology of mitotic chromosomes with indicated cohesin configurations. **e**, Bar graphs showing the average length of mitotic chromosomes with indicated cohesin configurations. *P* values were calculated using two-sided student’s *t*-test. **f**, Schematic showing the G2/M short-arrest and release protocols to obtain mitotic condensin-replete mitotic chromosomes with or without cohesive-cohesin. **g**, Upper panel: KR-balanced Hi-C contact maps of the mitotic Wapl^dTag^/Sororin^mAID^ cells with indicated cohesin configurations (short-arrest and release protocol). Bin size: 100kb. Lower panel: Saddle plots showing that compartments were missing in mitotic Wapl^dTag^/Sororin^mAID^ cells obtained through the G2/M short-arrest/release protocol. **h**, *P(s)* curves showing the influences (indicated by arrows) of enhanced occupancy of cohesive-cohesin (short-arrest and release protocol) on mitotic chromosome architecture. **i**, Bar graphs showing the average length of mitotic chromosomes with indicated cohesin configurations (short-arrest and release protocol). *P* values were calculated using two-sided student’s *t*-test.

*In-situ* Hi-C experiments uncovered the anticipated “featureless” contact maps in the control mitotic cells, showing no sign of compartments ^20,23,36^ (Fig. 3b). In the control samples (configuration 1), the distance dependent contact probability decay curve (*P(s)* curve) displayed three regimes: (i) an initial decay corresponding to contacts within mitotic loops; (ii) a following shallow decay reflecting inter-loop interactions within the same axial layer of the cylindrical mitotic chromosome and (iii) a steep decay at larger genomic separations, indicative of diminished inter-layer contacts along the chromosome axis ^23^ (Extended Data Fig. 11a). A vague secondary diagonal was located between regime (ii) and (iii), likely representing the helical winding of mitotic chromosomes (Extended Data Fig. 11a) ^23^. In the presence of extrusive-cohesin (configuration 2), compartments remained undetectable (Fig. 3b). Notably, the secondary diagonal in the *P(s)* curve was dramatically obscured (Fig. 3c left panel; Extended Data Fig. 11b). This finding was still evident, albeit to a lesser extent in cells with lower level of extrusive-cohesin (“light load”) (Extended Data Fig. 11b lower panel). These data imply that extrusive-cohesin may antagonize mitotic chromosome helical winding and promote a non-helical organization. Intriguingly, the average length of mitotic chromosomes was unaffected by extrusive-cohesin, indicating that extrusive-cohesin does not perturb mitotic chromosome longitudinal shortening (Fig. 3d, e).

Mitotic chromosomes loaded with cohesive-cohesin (configuration 3) exhibited no discernable changes in the Hi-C contact matrix and only marginal alterations on the *P(s)* curve (Fig. 3b, c middle panel; Extended Data Fig. 11c). Consistently, we failed to detect significant changes in mitotic chromosome length (Fig. 3d, e). We next wondered whether enhancement of the cohesive-cohesin level may induce more pronounced impacts. To approach this, we implemented the G2/M short-arrest and release protocol (Fig. 3f). Increased occupancy of cohesive-cohesin led to a markedly shortened regime (ii) on the *P(s)* curve, indicative of insufficient expansion of chromosome helix (Fig. 3g, h; Extended Data Fig. 11d, e) ^23^. In line with this observation, the average length of mitotic chromosomes displayed a slight but statistically significant increase by ∼17.8% (Fig. 3i). These findings collectively support a dosage-dependent suppressive effect of cohesive-cohesin on the expansion and longitudinal shortening of mitotic chromosome helical scaffold.

Finally, we examined the combined influence of extrusive- and cohesive-cohesin (configuration 4) on mitotic chromosome structure. We found that simultaneous loading of the two cohesin forms integrated their individual impacts and generated a *P(s)* with markedly shortened regime (ii) (effect of cohesive-cohesin) and dramatically attenuated secondary diagonal (effect of extrusive-cohesin) (Fig. 3c right panel; Extended Data Fig. 11f). These features jointly indicate a defect in mitotic chromosomes condensation. Remarkably, we noticed a significant gain of mitotic chromosome length by ∼50%, surpassing the sum of individual effects observed with extrusive-cohesin (∼0% increase) and cohesive-cohesin (∼17.8% increase) (Fig. 3e). This finding implied a synergistic influence of the two cohesin forms in delaying mitotic chromosome condensation. Furthermore, to exclude potential artifacts arising from the Sororin^9A^ mutant, we endeavored to recapitulate our observations in the Wapl^dTag^/SMC2^mAID^ cells, which expressed endogenous wildtype Sororin. Treating these cells with dTag13 followed by nocodazole-induced prometaphase arrest enabled simultaneous retention of extrusive- and cohesive-cohesin on mitotic chromosomes (Extended Data Fig. 12a). Notably, the resulting chromosomes displayed similar changes in the Hi-C contact matrix, *P(s)* curve, and chromosome length corroborating the collaborative role of extrusive- and cohesive-cohesin in hindering mitotic chromosome condensation (Extended Data Fig. 12b-e).

We conclude that extrusive- and cohesive-cohesin exert distinct influences on mitotic chromosome organization with the former prohibiting helical winding and the latter restricting helix expansion. When loaded together, these effects synergize. The coordinated interplay between cohesin and condensin may regulate the dynamic condensation of chromosomes during mitotic entry.

### Extrusive-but not cohesive-cohesin mediates TAD formation

It is well-established that arrested loop extrusion by cohesin at CTCF sites is crucial for the establishment of TADs in interphase cells ^5,7,8^. The distinct mechanisms that govern the positioning of extrusive- and cohesive-cohesin drove us to investigate their potential similar or divergent roles in TAD formation. To address this, we utilized the Wapl^dTag^*/*SMC2^mAID^*/*Sororin^mAID^ cell line either with or without Sororin^9A^ expression and obtained condensin-deficient mitotic chromosomes with four distinct cohesin configurations:

(5) condensin(-); extrusive-cohesin (-); cohesive-cohesin (-) (control)
(6) condensin(-); extrusive-cohesin (+); cohesive-cohesin (-)
(7) condensin(-); extrusive-cohesin (-); cohesive-cohesin (+, protected by Sororin^9A^)
(8) condensin(-); extrusive-cohesin (+); cohesive-cohesin (+, protected by Sororin^9A^).

These samples were then subject to *in-situ* Hi-C. G1-phase cells with or without Wapl were processed in parallel as interphase controls. We found that deleting condensin during mitosis caused a significant gain of inter-chromosomal *trans*-interactions from ∼5% to ∼26.5% (configuration 5), indicating increased intermingling of mitotic chromosomes (Extended Data Fig. 13a, b; Table S1). This finding supports the role of condensin in maintaining chromosome separation ^24^. Introduction of cohesive-cohesin (configuration 7) reduced *trans*-interactions to ∼20.7% (Extended Data Fig. 13a, b). Remarkably, extrusive-cohesin (configuration 6) displayed a more pronounced effect, decreasing the percentage of *trans*-contacts down to ∼10.2% (Extended Data Fig. 13a, b). Combining cohesive- and extrusive-cohesin (configuration 8) did not further reduce this percentage, suggesting that extrusive-cohesin plays a more prominent role than cohesive-cohesin to resolve entanglements of chromatids (Extended Data Fig. 13a, b). Our data are reminiscent with a recent report that loop extruding cohesin facilitates sister-chromatid resolution in G2/M-phase transition ^37^.

Analysis of the *P(s)* curves for condensin-deficient mitotic samples unveiled that extrusive-cohesin induced a striking increase in contacts for genomic separations below 10Mb, indicative of domain formation (Extended Data Fig. 14a). To further quantify this, we utilized the “Arrowhead” domain calling algorithm in combination with local minima of insulation score profiles (methods). Collectively, we identified 8,795 domains across all mitotic samples and interphase controls (Fig. 4a; Table S2). These domains were categorized into 2,073 (∼23.6%) CTCF dependent TADs and 1,415 (∼16.1%) CTCF independent compartmental domains (methods). Compared to TADs, compartmental domains were significantly smaller in size and more enriched with active histone marks (Fig. 4b, c). Consistent with our previous finding ^24^, condensin-deficient control mitotic samples without cohesin (configuration 5) were dominated by compartmental domains (∼40%) (Fig. 4d, e). Introduction of extrusive-cohesin (configuration 6) resulted in 2,479 domains, with ∼24.3% classified as TADs, a percentage comparable to interphase controls (∼28.9%) (Fig. 4a, d). Formation of TADs by extrusive-cohesin were further corroborated by the “corner-dot” signals in the composite domain plots (Fig. 4e upper left panel; Extended Data Fig. 15a). Co-loading of cohesive-cohesin (configuration 8) failed to induce any increase in the proportion of TADs (∼24.3%) (Fig. 4d). In addition, cohesive-cohesin by itself (configuration 7) failed to induce algorithmically detectable TADs (Fig. 4d, e upper left panel). Taken together, our data demonstrate that extrusive-but not cohesive-cohesin were capable to mediate TAD formation. It is noteworthy that condensin-replete mitotic cells displayed negligible TAD signatures (<60 domain calls), irrespective of their cohesin states (configurations: 1-4) (Fig. 4a, e upper right panel). This finding supports the notion that condensin contributes to TAD disassembly during mitotic entry, presumably through dislocating extrusive-cohesin from CTCF binding sites.

**Figure 4.**
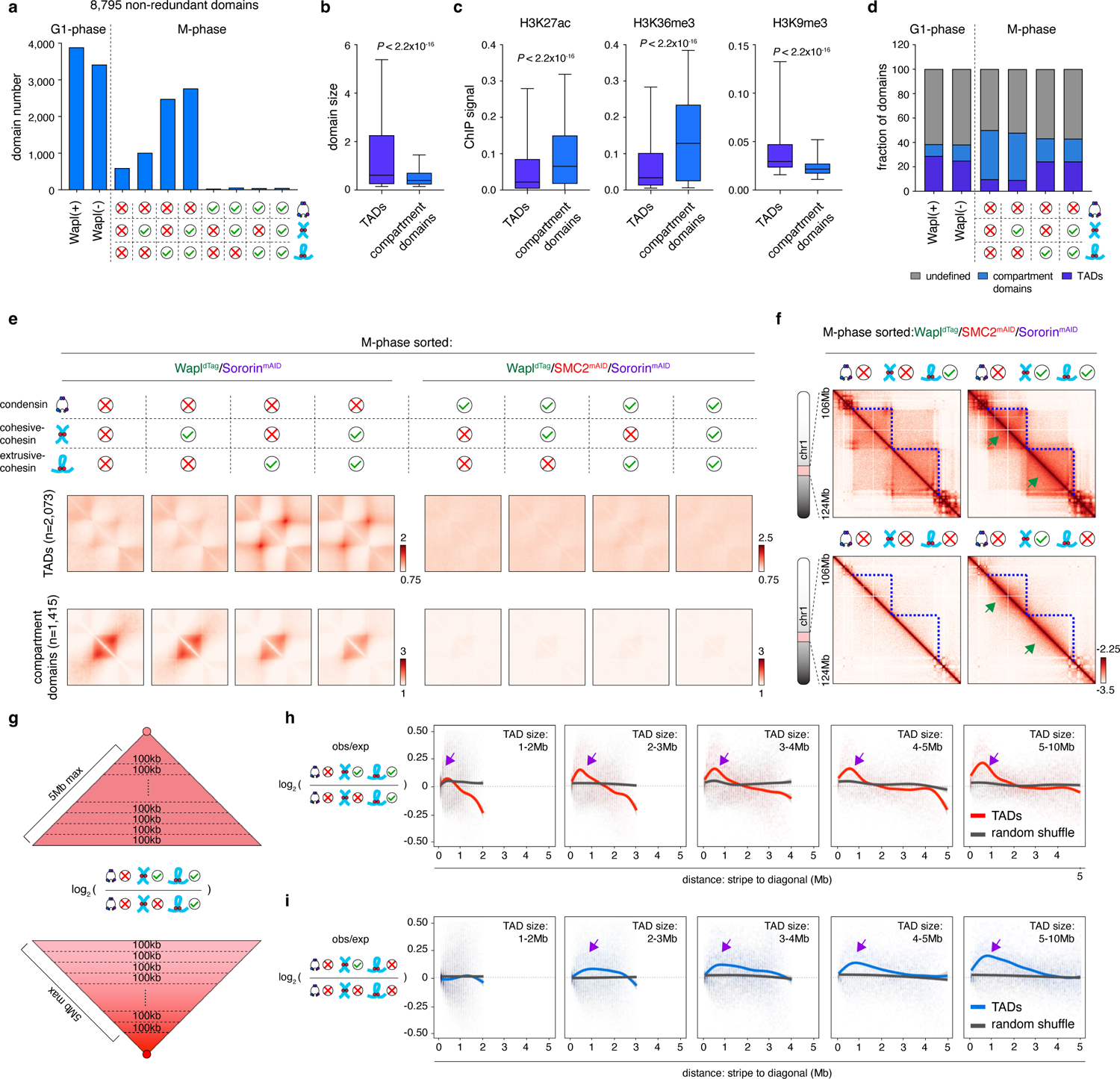
Cohesive-cohesin intensifies extrusive-cohesin mediated TADs. **a**, Bar graph showing the number of domains identified in interphase and mitotic samples. **b**, Box plots showing that TADs are significantly larger than compartment domains. For both box plots, central lines denote medians; box limits denote 25th–75th percentile; whiskers denote 5th–95th percentile. *P* values were calculated using a two-sided Wilcoxon signed-rank test. **c**, Box plots showing that compartmental domains displayed higher levels of active histone marks and lower levels of repressive mark than TADs. For both box plots, central lines denote medians; box limits denote 25th–75th percentile; whiskers denote 5th–95th percentile. *P* values were calculated using a two-sided Wilcoxon signed-rank test. **d**, Bar graph showing the fraction of TADs and compartmental domains in interphase and mitotic samples. **e**, Re-scaled composite domain plots of TADs and compartmental domains mitotic samples with eight types of SMC protein configurations. **f**, KR-balanced Hi-C contact maps showing representative TADs in condensin-deficient mitotic samples with indicated cohesin states. TADs were marked by blue dotted lines. Intensification of TADs were marked indicated by green arrows. Bin size: 25kb. **g**, Schematic showing the segmentation of domains into 100kb-wide stripes. **h**, Line plots showing the gain of short-range interactions (purple arrows) within TADs in the presence of cohesive-cohesin. TADs of different size-range were separately shown. Note that random position control did not show such gain of interactions. **i**, Line plots showing the gain of short-range interactions within pre-defined TAD regions (purple arrows) induced by cohesive-cohesin in the absence of extrusive-cohesin.

### Cohesive-cohesin intensifies TADs

We observed that co-appearance of cohesive-cohesin with extrusive-cohesin further enhanced the contact probability for genomic separations below 5Mb on the *P(s)* curve (Extended Data Fig. 14b). This finding suggested a potential role of cohesive-cohesin in strengthening pre-existing TADs. Visual inspection of the Hi-C contact maps for two independent loci revealed more robust intra-TAD interactions upon the introduction of cohesive-cohesin (Fig. 4f upper panel; Extended Data Fig. 15b). To quantify this observation, we segmented each TAD into 100kb-wide stripes (Fig. 4g). The log_2_ fold change in contact frequency between cohesive-cohesin replete and depleted conditions (configuration 8 vs. 6) was calculated for each stripe (Fig. 4g). We observed a progressive increase in contact frequency in response to cohesive-cohesin as the stripe diverged from the diagonal, reaching a peak at ∼500kb of genomic separations (Fig. 4h; Extended Data Fig. 15c). This pattern held true for TADs with varying sizes (Fig. 4h; Extended Data Fig. 15c). Similar observations were made when TADs were established by low levels of extrusive-cohesin (light load) (Extended Data Fig. 16a, b). Notably, randomly selected control regions failed to display any increase in contact frequency upon cohesive-cohesin loading (Fig. 4h; Extended Data Fig. 15c, 16b). Taken together, our data highlight a previously unnoted role of cohesive-cohesin to strengthen TADs.

We speculate that cohesive-cohesin induced TAD intensification may arise from either enhanced inter-sister-chromatid *trans*-interactions or from reinforced intra-chromatid *cis*-interactions within TADs. To dissect the contributions of *trans*-contacts to TAD reinforcement, we sought to ablate the intra-TAD *cis*-interactions by removing extrusive-cohesin and compare residual TAD strength for samples with or without cohesive-cohesin (configuration 7 vs. 5). Consistent with our previous report ^24^, the condensin-deficient control mitotic chromosomes without any cohesin (configuration 5) displayed negligible TAD signature (Fig. 4f lower panel; Extended Data Fig. 15b). Remarkably, we observed visually discernable intra-TAD contacts when cohesive-cohesin was present (configuration 7) (Fig. 4f lower panel, 4i; Extended Data Fig. 15b, d). Similar findings were made in samples obtained through the G2/M short-arrest and release protocol (Extended Data Fig. 17a-d). Given the absence of extrusive-cohesin, these residual TAD signatures were likely to be attributable to cohesive-cohesin mediated *trans*-contacts. It is noteworthy that these interactions were relatively weak and did not manifest as clear triangular domain-like structures on the Hi-C contact map, potentially explaining the failure of domain calling algorithms (Fig. 4f lower panel; Extended Data Fig. 15b). Taken together, our results indicate that cohesive-cohesin may intensify TADs by directly mediating inter-chromatid *trans*-contacts. However, we do not rule out the possibility that cohesive-cohesin may modulate extrusive-cohesin function and promote intra-TAD *cis*-interactions.

### Cohesive-cohesin restricts extrusive-cohesin mediated chromatin loop expansion

Comparative analysis of the composite TAD plots unveiled a reduction of the corner-dot signature when cohesive-cohesin was co-present with extrusive-cohesin, implying that cohesive-cohesin might interrupt extrusive-cohesin’s looping function (Fig. 4e upper left panel). Utilizing a modified HICCUPS approach, we detected 45,242 loops across all mitotic samples and interphase controls (Table S3). Among them, 26,086 (∼57.7%) were classified as structural loops based on the co-occupancy of CTCF and cohesin at both anchors. In the condensin-deficient mitotic cells, extrusive-cohesin (configuration 6) generated 14,437 loops, of which 9,027 (∼62.5%) were structural loops (Fig. 5a, b). Therefore, extrusive-cohesin maintains a capacity to establish structural loop during mitosis. Conversely, cohesive-cohesin alone (configuration 7) failed to induce structural loops, as evidenced by the negligible aggregated peak analysis (APA) signals (Fig. 5b). Interestingly, co-presence of cohesive- and extrusive-cohesin (configuration 8) resulted in a slight decrease of structural loop calls (7,428) compared to extrusive-cohesin alone. Visual inspection of individual genomic locus and genome-wide aggregated plots both confirmed a reduction of structural loop signal intensity upon the introduction of cohesive-cohesin (Fig. 5b, c; Extended Data Fig. 18a, b). Notably, structural loops were barely detectable in the presence of condensin (configurations: 1-4) (Fig. 5a, b; Extended Data Fig. 18a), consolidating that condensin participates to deconstruct chromatin loops during mitotic entry.

**Figure 5.**
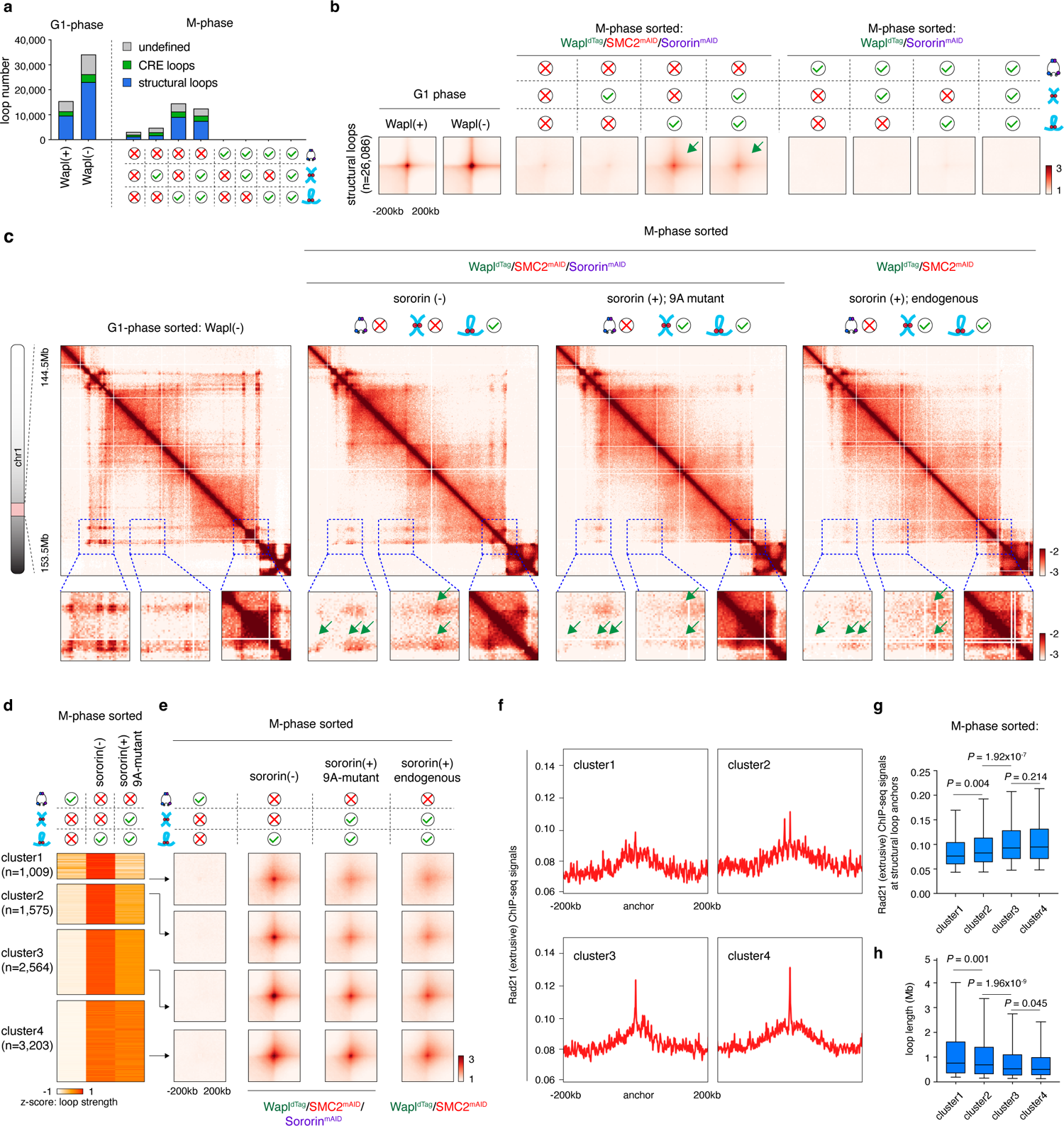
Cohesive-cohesin restricts extrusive-cohesin mediated structural loops. **a**, Bar graph showing the number of loops identified by HICCUPS in interphase and mitotic samples. The fraction of structural and CRE loops in each sample were also shown. **b**, APA plots showing the composite structural loop signals in interphase and mitotic samples with eight distinct SMC protein complex configurations. Note that structural loops were appreciable in condensin-deficient mitotic cells loaded with extrusive-cohesin (green arrows). **c**, KR-balanced Hi-C contact maps showing the weakening of structural loops (green arrow) upon the introduction of cohesive-cohesin during mitosis. Maps for Wapl^dTag^/SMC2^mAID^ cells which expresses endogenous wildtype Sororin were shown on the right. Bin size: 25kb. **d**, Heatmap showing the clustering result structural loops based on their responses to cohesive-cohesin. **e**, APA plots for each cluster in (**d**) showing loss of structural loop signal intensity in the presence of cohesive-cohesin. Plot for Wapl^dTag^/SMC2^mAID^ cells which expresses endogenous wildtype Sororin were shown on the right. **f,** Meta-region plots showing Rad21 ChIP-seq signal strength at structural loop anchors from each cluster. **g**, Box plots showing the quantification of (**f**). For both box plots, central lines denote medians; box limits denote 25th–75th percentile; whiskers denote 5th–95th percentile. *P* values were calculated using a two-sided Wilcoxon signed-rank test. **h**, Box plots showing the sizes of structural loops from cluster1, cluster2, cluster3 and cluster4. For both box plots, central lines denote medians; box limits denote 25th–75th percentile; whiskers denote 5th–95th percentile. *P* values were calculated using a two-sided Wilcoxon signed-rank test.

To further elucidate the influence of cohesive-cohesin on structural loops, we performed unsupervised *k-means* clustering and subcategorized the 9,027 mitotic structural loops into four clusters (Fig. 5d). Cluster1 exhibited a dramatic reduction of structural loop signal intensity in response to cohesive-cohesin, while cluster2, 3 and 4 were progressively more resistant (Fig. 5d, e; Extended Data Fig. 18c). These patterns were maintained when extrusive-cohesin was present at lower level (“light load”) (Extended Data Fig. 16c, d). Importantly, an inverse correlation emerged between extrusive-cohesin occupancy at loop anchors and loop sensitivity to cohesive-cohesin: i.e., the most sensitive cluster1 loops displayed the lowest abundance of extrusive-cohesin at their anchors, while the most resistant cluster 4 loops exhibited the highest enrichment of anchor bound extrusive-cohesin (Fig. 5f, g). This observation indicates a dose-dependent effect, where high levels of extrusive-cohesin confer greater resistance to cohesive-cohesin mediated interruption. Furthermore, we found that cluster1 loops were significantly larger in size compared to the other clusters, suggesting that larger loops are more sensitive to cohesive-cohesin mediated disruption (Fig. 5h). These results are consistent with a recent report that sister-chromatid cohesion restricts loop expansion in yeast ^18^.

To rule out the possibility that cohesive-cohesin mediated structural loop interruption was an artifact induced by the Sororin^9A^ mutant, we employed Wapl^dTag^*/*SMC2^mAID^ cell line, which expressed wildtype Sororin. We applied the G2/M arrest and release protocol and obtained condensin-deficient mitotic chromosomes co-occupied by extrusive- and cohesive-cohesin: (9). condensin(-); extrusive-cohesin (+); cohesive-cohesin (+, protected by endogenous Sororin).

Comparative analysis of this sample with the condensin-deficient mitotic chromosomes harboring only extrusive-cohesin (configuration 6) revealed a reduction of structural loop strength across all four clusters (Fig. 5c, e; Extended Data Fig. 18b, c). This finding confirmed a weakening effect of cohesive-cohesin on structural loops when coupled with endogenous Sororin.

Taken together, we provide compelling evidence that cohesive-cohesin antagonizes the formation of structural loops, with a more pronounced effect on larger loops. Our observations from mitotic cells imply a potential regulatory role for cohesive-cohesin in TAD and loop formation during S- and G2-phases.

### Cohesive-cohesin does not affect the insulation function of structural loop anchors

We previously demonstrated that structural loop anchors were capable of insulating contacts between CREs during interphase ^10^. Given that cohesive-cohesin impairs structural loop integrity, we wondered whether it also influences the insulation function of structural loops. We identified 11,399 CRE contacts across all interphase and mitotic samples (methods) (Extended Data Fig. 19a; Table S4). APA plots revealed prominent CRE loop signals in the condensin-deficient mitotic chromosomes (configuration 5), corroborating our prior finding that CRE contacts emerge during mitosis upon condensin depletion (Extended Data Fig. 19b) ^24^. Notably, we observed a progressive decline in CRE contact strength as the number of internal structural loop anchors increased, confirming the capacity of structural loops to interrupt CRE contacts on mitotic chromosomes (Extended Data Fig. 19c, d).

To examine the impact of cohesive-cohesin on structural loop mediated CRE insulation, we stratified four distinct groups of anchors, each belonging to a specific structural loop cluster (Table S5). For each anchor set, we identified all CREs located within a 500kb radius. We then paired up these CREs, parsed out those that flank the target anchor and computed their aggregated interacting strength (Extended Data Fig. 19e). Cohesive-cohesin by itself (configuration 7) failed to insulate these CRE contacts (Extended Data Fig. 19g, f), consistent with our finding that cohesive-cohesin does not establish structural loops. In the presence of extrusive-cohesin (configuration 6), all anchor sets displayed an insulating effect with cluster4 anchors showing the most prominent insulation and cluster1 anchors exhibiting a weaker effect (Extended Data Fig. 19g, f). Surprisingly, co-presence of cohesive-cohesin (configuration 8) failed to restore the insulated CRE contacts, even for those separated by cluster1 loop anchors (Extended Data Fig. 19g, f). This result was unexpected given that cluster1 loop intensity was substantially diminished by cohesive-cohesin. In summary, our analyses indicate that cohesive-cohesin, while counteracting structural loop formation, does not compromise the insulation function of structural loop anchors. These findings imply a potential decoupling of the looping function and insulation capacity of extrusive-cohesin.

### Extrusive-but not cohesive-cohesin modulates mitotic chromosome compartments

It was well established that cohesin suppresses chromatin compartmentalization during interphase^7,8^. We next examined whether extrusive- and cohesive-cohesin exert similar or distinct effects on chromatin compartments. As expected, condensin-deficient mitotic chromosomes without cohesin (configuration 5) displayed a clear checkerboard pattern of compartments (Fig. 6a, b). In line with our prior report, these compartments could be partitioned into four different types: mA1, mA2, mB1 and mB4 (Extended Data Fig. 20a) ^24^. The presence of cohesive-cohesin (configuration 7) alone did not alter the strength or patterns of mitotic compartments as evidenced by comparable Hi-C contact matrices, saddle plots, aggregation-repulsion plots and well-correlated eigenvector 1 (EV1) values (Fig. 6a-e; Extended Data Fig. 20a, b). Cohesive-cohesin loaded through the G2/M short arrest and release protocol yielded similar results (Extended Data Fig. 17e, f). Therefore, cohesive-cohesin does not appear to affect mitotic chromosome compartmentalization.

**Figure 6.**
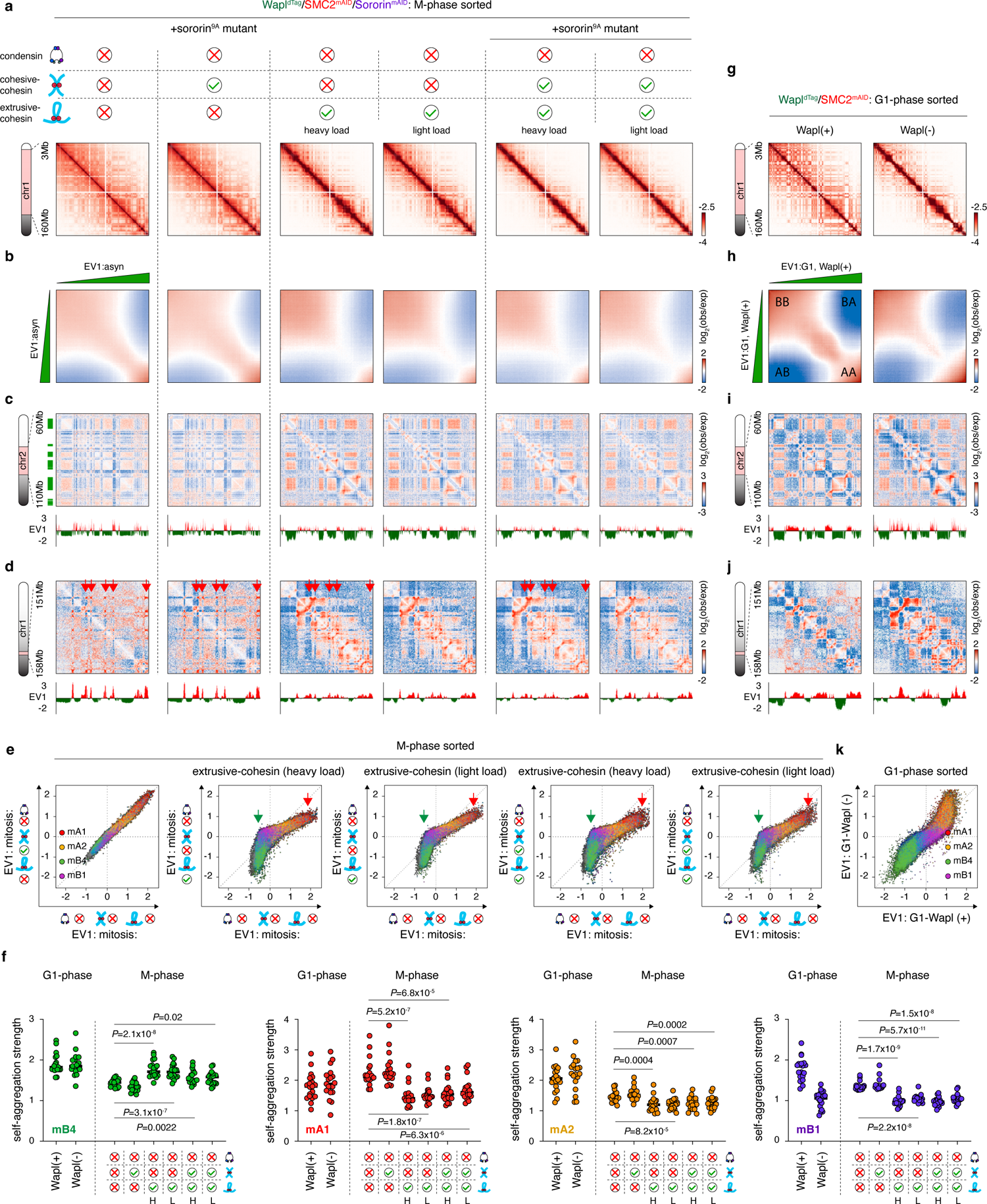
Extrusive-but not cohesive-cohesin influences mitotic specific compartmentalization patterns. **a**, KR-balanced Hi-C contact matrices of condensin-deficient mitotic chromosomes, showing the checkerboard compartmentalization patterns in condensin-deficient mitotic chromosomes with indicated configurations of cohesin. Bin size: 100kb. **b**, Saddle plots of condensin-deficient mitotic chromosomes. **c**, KR-balanced Hi-C contact matrices showing the increased homotypic interactions among mB4 mitotic compartments in the presence of extrusive-cohesin (“heavy” or “light load”) but not cohesive-cohesin. Bin size: 100kb. EV1 tracks were shown in parallel mB4 compartments were marked by green bars. **d**, KR-balanced Hi-C contact matrices showing loss of mA1 homotypic interactions (red arrows) in the presence of extrusive-cohesin (“heavy” or “light load”) but not cohesive-cohesin. Bin size: 24kb. EV1 tracks were shown in parallel. **e**, Scatter plots showing the EV1 values of 25kb genomic bins in the condensin-deficient mitotic chromosomes without any cohesin (*x*-axis) against condensin-deficient mitotic chromosomes with cohesive-cohesin, extrusive-cohesin (“heavy” or “light load”) or both (*y*-axes). Bins were color coded based on their compartment assignment. Green and red arrows indicate the reduction of EV1 values of bins within mB4 and mA1 compartments respectively. **f**, Dot plots showing the homotypic self-interaction strength of indicated mitotic compartments (mB4, mA1, mA2 and mB1) in the condensin-deficient mitotic samples. G1-phase samples were shown as control. *P* values were calculated using a two-sided Wilcoxon signed-rank test. **g**, KR-balanced Hi-C contact matrices of G1-phase samples with or without Wapl. Bin size: 100kb. **h**, Saddle plots showing the compartment changes induced by Wapl depletion in G1-phase. **i**, KR-balanced Hi-C contact maps showing mB4 self-interaction in G1-phase samples with or without Wapl. EV1 tracks were shown in parallel. Bin size: 100kb. **j**, KR-balanced Hi-C contact maps showing mA1 self-interaction in G1-phase samples with or without Wapl. EV1 tracks were shown in parallel. Bin size: 25kb. **k**, Scatter plots showing EV1 values of 25kb genomic bins in the control G1-phase sample (*x*-axis) and Wapl-deficient G1-phase sample (*y*-axis). Bins were color coded based on their compartment assignment.

Extrusive-cohesin (configuration 6) led to reduced separation of mA1 or mA2 (euchromatin) from mB4 (constitutive heterochromatin) compartments (Extended Data Fig. 20a), consistent with the notion that cohesin counteracts genome compartmentalization by juxtaposing chromatin of opposing states ^8^. Strikingly, we observed a massive gain of mB4 homotypic interactions and a corresponding decrease in EV1 values (Fig. 6c, e, f; Extended Data Fig. 20a, c). Conversely, self-association within mA1, mA2 or mB1 compartments (facultative heterochromatin) were substantially attenuated (Fig. 6d, f; Extended Data Fig. 20a). EV1 values corresponding of mA1 or mA2 compartments were also decreased (Fig. 6e; Extended Data Fig. 20c). These findings were recapitulated in samples lightly loaded with extrusive-cohesin (Fig. 6a-f; Extended Data Fig. 20a, c). Co-loading of cohesive-cohesin (configuration 8) did not impose additional alterations to compartments (Fig. 6a-f; Extended Data Fig. 20). Notably, Wapl degradation mediated excessive loading of extrusive-cohesin in interphase cells failed to reproduce these changes (Fig. 6g-k). In conclusion, these findings demonstrate that extrusive-but not cohesive-cohesin possesses the capacity to modulate mitotic chromosome compartmental interactions. Using condensin-deficient mitotic chromosomes, our data illuminate a context-dependent influence of extrusive-cohesin on genome compartmentalization, that was underappreciated in interphase cells. Specifically, extrusive-cohesin promotes constitutive heterochromatin self-association and meanwhile disrupts the self-aggregation of euchromatin or facultative heterochromatin.

## Discussion

In this study, we adopted mitotic chromosomes as a unique platform to accommodate all possible configurations of extrusive-cohesin, cohesive-cohesin and condensin. We describe a tripartite pattern of interplay among these three types of SMC protein complexes.

### Condensin vs. extrusive-cohesin

We provide compelling evidence that condensin effectively eliminates the focal peaks of extrusive-cohesin. Two mechanisms may potentially explain this disruption: (1) condensin destabilizes the interaction between extrusive-cohesin and CTCF, pushing it away while leaving the genome-wide overall occupancy unaffected (relocation model, Fig. 7a); (2) condensin directly expels extrusive-cohesin from the chromatin (eviction model, Fig. 7a). Immunofluorescence staining revealed that ∼90.7%±5.6% of Wapl and SMC2 co-deficient mitotic cells displayed Rad21 retention. In comparison, this percentage dropped to ∼66.8%±7.7% in the presence of SMC2. Furthermore, live cell imaging demonstrated a marked increase in the mitotic enrichment of fluorescently labeled Rad21 when SMC2 was lost (data not shown). Finally, a recent study using chromatin enrichment for proteomics (ChEP) technology reported more rapid cohesin dissociation in the presence of condensin during mitotic progression ^38^. These findings collectively support condensin’s capacity to unload extrusive-cohesin independently of Wapl. Furthermore, our observations also suggest a potential role for condensin in regulating chromosome landscape by relocating or displacing additional chromatin-associating factors during mitosis.

**Figure 7.**
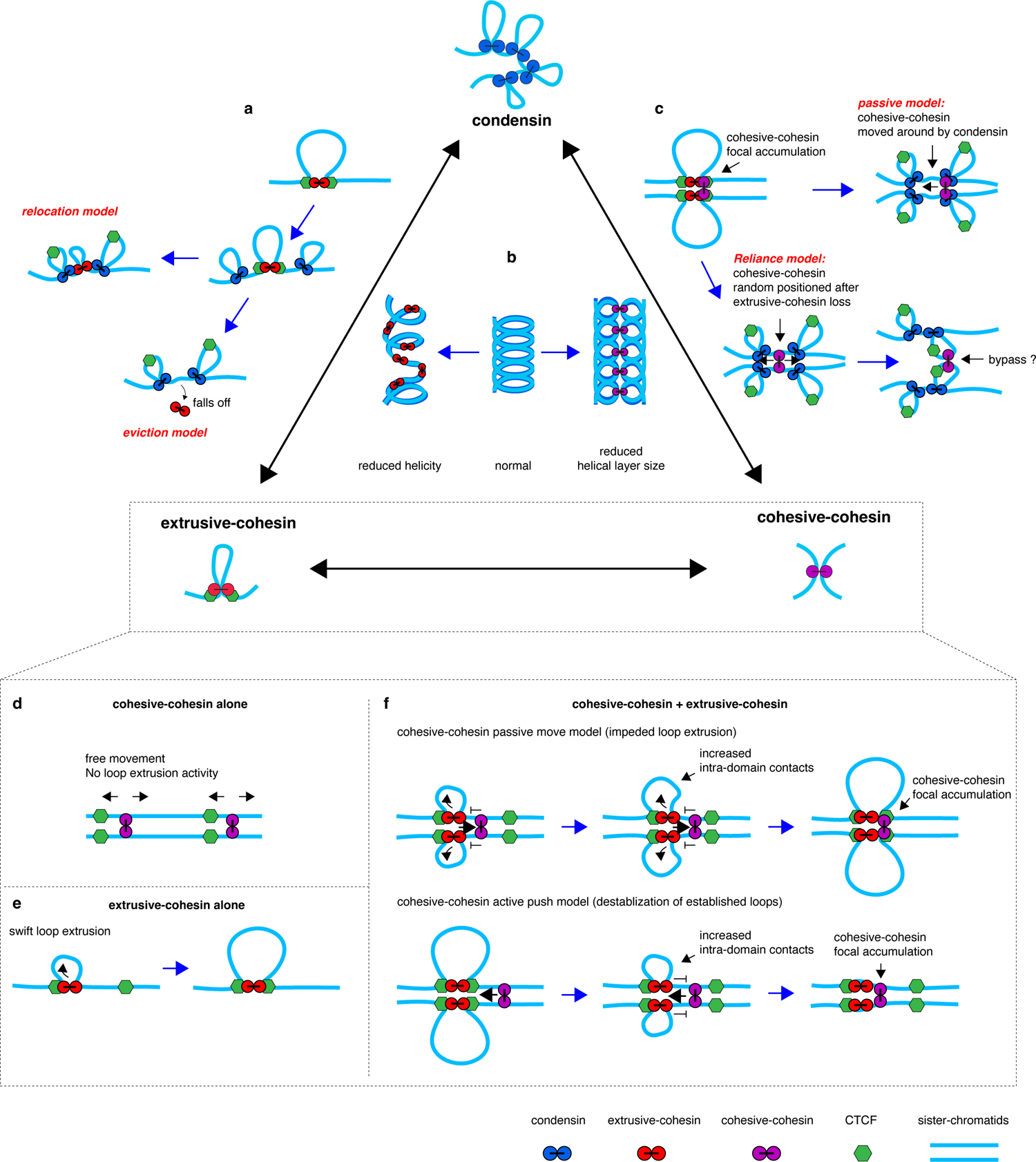
The tripartite pattern of interactions among extrusive-cohesin, cohesive-cohesin and condensin. **a**, Schematic illustration showing two potential mechanisms (relocation model and eviction model) through which condensin disrupts extrusive-cohesin positioning. **b**, Schematic illustration showing the influence of extrusive-cohesin or cohesive-cohesin on mitotic chromosome structure. **c**, Schematic illustration showing the two potential models (passive model and reliance model) through which condensin disrupts cohesive-cohesin focal peaks. **d**, Schematic showing that cohesive-cohesin moves along cohered sister-chromatids without extruding loops. **e**, Schematic showing the classic loop extrusion process. **f**, Upper panel, schematic showing that extrusive-cohesin pushes cohesive-cohesin along the DNA fiber and cohesive-cohesin serves as an obstacle impeding loop expanding by extrusive-cohesin. Cohesive-cohesin is positioned by extrusive-cohesin at CTCF binding sites. Lower panel, schematic showing that cohesive-cohesin actively destabilizes extrusive-cohesin from CTCF binding sites, thereby interrupting chromatin loop formation.

To evaluate the impact of extrusive-cohesin on condensin’s loop extrusion activity, we performed *in-situ* Hi-C. We observed a dose-dependent attenuation in the secondary diagonal of the *P(s)* curve by extrusive-cohesin. Obscuration of the secondary diagonal was previously described by our group and others in mitotic chromosomes devoid of condensin II (Extended Data Fig. 11g) ^23,24^. Therefore, extrusive-cohesin may antagonize condensin II activity and prohibit mitotic chromosomes coiling (Fig. 7b). We speculate that extrusive-cohesin may function as roadblocks on the chromatin fiber and impede extruding chromatin loops motorized by condensin II.

### Condensin vs. cohesive-cohesin

Our data indicate that condensin diminishes cohesive-cohesin peaks. Of note, unlike extrusive-cohesin, which potentially undergoes condensin-mediated eviction, cohesive-cohesin is more likely to be redistributed rather than displaced. This is evidenced by extensive sister-chromatid cohesion in the condensin-replete mitotic cells. Two potential models may explain condensin-mediated cohesive-cohesin relocation: (1) a passive model wherein cohesive-cohesin is passively pushed away from its focal binding sites by condensin (Fig. 7c). (2) a reliance model in which cohesive-cohesin depends on extrusive-cohesin to form focal peaks. Once extrusive-cohesin peaks are disrupted by condensin, cohesive-cohesin moves away from the original accumulating sites, leading to disappearance of focal peaks (Fig. 7c). Three lines of evidences support the reliance model. First, cohesive-cohesin exhibits similar responses to condensin as extrusive-cohesin. Second, cohesive-cohesin fails to form detectable peaks in the absence of extrusive-cohesin. Finally, a recent study using polymer simulation and immunofluorescence staining predicted that condensin is able to bypass cohesive-cohesin on mitotic chromosomes, suggesting that condensin does not impose direct mechanical influence on cohesive-cohesin (Fig. 7c) ^38^. These findings together indicate that disruption of cohesive-cohesin peaks by condensin is a secondary consequence of diminished extrusive-cohesin focal enrichment.

*In-situ* Hi-C analysis using condensin-replete mitotic chromosomes solely loaded with cohesive-cohesin uncovered a shortened regime (ii) in the *P(s)* curves. This observation inversely mirrored our prior finding that condensin I depletion extended regime (ii), leading to dramatically shorter mitotic chromosomes (Extended Data Fig. 11h) ^24^. Therefore, loading of cohesive-cohesin appeared to promote condensin I activity, ultimately achieving longer and narrower mitotic chromosomes (Fig. 7b). Notably, cohesive-cohesin preserved a sharp secondary diagonal in the *P(s)* curve, indicating a minimal influence on chromosome coiling (Fig. 3h). Our findings are consistent with a recent study showing that cohesive-cohesin delays chromosome condensation during mitotic progression ^38^.

### Extrusive-cohesin vs. cohesive-cohesin

A previous study using sister-chromatid-sensitive Hi-C (scsHi-C) uncovered that cohesive-cohesin mediated *trans*-chromatid interactions generated structures resembling TADs ^39^. This led to a hypothesis that cohesive-cohesin may mediate TAD formation. Here, using condensin-deficient mitotic chromosomes loaded exclusively with cohesive-cohesin, we demonstrate that cohesive-cohesin by itself, is incapable of establishing algorithmically identifiable TADs or chromatin loops (Fig. 7d, e). Furthermore, cohesive-cohesin does not influence mitotic chromosome compartmentalization or facilitate chromosome individualization. Together, these findings highlight the non-overlapping functionalities of the two cohesin forms. Our results are consistent with studies showing distinct mechanisms for cohesin mediated loop extrusion and chromatid cohesion ^14,40^.

It was previously observed that cohesive-cohesin accumulates at CTCF binding sites following DNA replication ^32^. However, our data suggest that cohesive-cohesin is not directly stalled by CTCF. Instead, its focal accumulation appears to be contingent upon the presence of extrusive-cohesin. Mechanistically, two potential scenarios could explain this observation: (1) Upon engagement, extrusive-cohesin pushes cohesive-cohesin along the DNA fiber, until arrested by convergently oriented CTCF molecules (Fig. 7f, upper panel). (2) Alternatively, cohesive-cohesin may actively scan along the chromatin and relocate extrusive-cohesin until the latter is halted by CTCF (Fig. 7f, lower panel). Both scenarios predict an increase in the number of intermediate transient loops within TADs and a decrease in the signal strength of domain summit loops (Fig. 7d). Consistent with these predictions, we observed that cohesive-cohesin intensifies intra-TAD interactions while reducing genome-wide loop signals. Importantly, we found that larger loops are more susceptible to the influence of cohesive-cohesin than smaller ones. This could be explained by a higher likelihood of extrusive-cohesin to encounter cohesive-cohesin barriers during translocation over larger genomic distances. This finding therefore argues in favor of the first scenario. Although loops are attenuated by cohesive-cohesin, their insulation capability remains intact. This finding suggests a potential decoupling of looping and insulation functions by extrusive-cohesin.

### Implications throughout the cell cycle

The rules of engagement between SMC complexes as described above may also apply to interphase genome. Two recent studies have shed light on the interplay between extrusive-cohesin and the DNA replication machinery ^41,42^. Specifically, one study suggested that the MCM helicase complex involved in DNA replication may act as a physical barrier, hindering extrusive-cohesin mediated loop expansion. Here, we propose that cohesive-cohesin established after DNA replication, could serve as the roadblock for extrusive-cohesin. Another study uncovered that structural loop anchors might determine the initiation site of DNA replication ^42^. This finding offered a potential explanation for the positional dependence of cohesive-cohesin on extrusive-cohesin. These reports along with our findings highlight the functional interplay between extrusive- and cohesive-cohesin during S- and G2-phase. Future investigations are required to determine whether the influence of cohesive-cohesin on chromatin loops translates into altered transcriptional outcome.

It is well documented that Wapl facilitates the removal of extrusive-cohesin during mitotic progression. However, the potential contribution of condensin to this process remained unclear. Our findings present a novel perspective that condensin loading during prophase facilitates the eviction of extrusive-cohesin, thereby contributing to the disassembly of interphase chromatin structures such as TADs and loops. Furthermore, our live-cell imaging experiments demonstrate that condensin remains associated with the genome throughout anaphase and telophase during mitotic exit (Extended Data Fig. 1j) ^24^. This association may explain the previously observed slow recruitment of extrusive-cohesin in ana/telophase ^20^. Collectively, our findings imply a potential role of condensin regulating chromatin landscape as cells transit from G2-phase into mitosis and then into the next cell cycle.

In summary, by leveraging a series of orthogonal acute inducible degradation cell lines, we comprehensively dissected the interplay among extrusive-cohesin, cohesive-cohesin and condensin. Our work reveals a novel and intricate regulatory network governing these three factors during the dynamic remodeling of chromosome architecture from interphase to mitosis.

## Acknowledgements

We thank Dr. Marit Vermunt, Dr. Haizhen Long, Dr. Kai Huang, Ms. Jessica Lam and members of the Zhang lab for helpful discussions. We are indebted to Mengyuan Li, Zhiqing Huang and Dechun cheng from the Flow-cytometry Core of Shenzhen Bay Laboratory for technical support on cell sorting. We thank Shixian Huang and Mei Yu from the Bioimaging Core of Shenzhen Bay Laboratory for imaging support.

## Author contributions

H.Z. (Haoyue Zhang) conceived the study and designed experiments. H.Z. (Han Zhao) and L.S. created the inducible acute degradation cell line used in this study. H.Z. (Han Zhao) and L.S. performed sample preparation, FACS sorting, *in-situ* Hi-C, ChIP-seq and microscopic experiment with help from E.L., B.W., M.W., F.S., Y.L., L.Z. and Y.G. Data analysis were performed by H.Z. (Haoyue Zhang) with help from F.L. and S.X. H.Z. (Haoyue Zhang) wrote the manuscript with inputs from all authors.

## Declaration of interests

The authors declare no competing interests.

## Methods

### Cell culture

The G1E-ER4 parental ^43^ and all engineered cell lines were maintained in IMDM medium containing 15% FBS (v/v), 2% penicillin-streptomycin (v/v), 50ng/ml murine stem cell factor, 7.5ng/ml Epogen and 1:100000 1-Thioglycerol (v/v). Cells were maintained below a density of 1 million/ml. All G1E-ER4 derived cell lines were periodically tested to be negative for mycoplasma. All cell lines were cultured in a humidified incubator at 37°C, with 5% CO2.

### CRISPR/Cas9 mediated genome editing

The Wapl^dTag^/SMC2^mAID^ cell line was generated on the basis of a previously generated G1E-ER4:SMC2^mAID^ cell line. To enable orthogonal degradation of Wapl, we took advantage of the PROTAC strategy. The repair template donor containing the GFP-FKBP^F36V^ tag was designed as below: (1) a 540bp fragment covering the 5’ UTR of the *Wapl* gene before start codon, was PCR amplified from mouse genomic DNA as the left homology arm; (2) a 513bp fragment after the start codon of the *Wapl* gene was amplified as the right homology arm; (3) GFP and FKBP^F36V^ fragments were separately amplified, assembled together with the left and right homology arms through Gibson assembly and cloned into pcDNA3.1 plasmids. A sgRNA targeting the 5’ of the *Wapl* gene was designed by Benchling online tool and cloned into the pLenti-Crispr-TagBFP plasmid. 18μg of Donor template and 18μg of the sgRNA containing pLenti-Crispr-TagBFP plasmid were co-transfected into 5 million G1E-ER4:SMC2^mAID^ cells with an Amaxa 2b nucleofector using the program G16. 24h after transfection, single cells with high GFP signals were sorted into 96 well plates. 7 days later, homozygous knock-in was confirmed by genotyping using PCR primers outside the homology arms.

The Wapl^dTag^/Sororin^mAID^ cells were generated using similar strategies. To begin with, a G1E-ER4:Wapl^dTag^ cell line was created by applying the above described protocol on the parental G1E-ER4 cells. On the basis of this line, a minimal auxin inducible degron (mAID) tag was then inserted into the C-terminus of endogenous Sororin. In detail, the repair template containing the mAID-HaloTag was constructed as below: (1) a 856bp fragment before the stop codon of the *cdc5a* gene was amplified from genomic DNA as the left homology arm; (2) a 825bp fragment after the stop codon of *cdc5a* gene was amplified as the right homology arm; (3) mAID and HaloTag was amplified separately and assembled together with the left and right homology arms using Gibson assembly and cloned into pcDNA3.1 plasmids. sgRNA targeting the 3’ of the *cdc5a* gene was designed by Benchling online tool and cloned into pLenti-Crispr-TagBFP plasmid. Transfection was performed as described above. 24h after transfection, the cells were stained with 50nM Janelia Fluor® 646 (JF646) HaloTag ligand for 30min, washed three times with PBS and then subject to FACS. Single cells with high JF646 signals were sorted into 96 well plates. 7 days later, homozygous knock-in was confirmed by genotyping using PCR primers outside the homology arms.

The Wapl^dTag^/SMC2^mAID^/Sororin^mAID^ cells were generated on the basis of Wapl^dTag^/Sororin^mAID^ cells. mAID was fused to the C-terminus of SMC2 as described previously. Homozygous knock-in was confirmed by genotyping using PCR primers outside the homology arms. Oligos used in genome editing and genotyping are listed in Table S6

### Retroviral infection of murine cells

To enable rapid degradation of SMC2 or/and Sororin, OsTIR2 were delivered into Wapl^dTag^/SMC2^mAID^, Wapl^dTag^/Sororin^mAID^ and Wapl^dTag^/SMC2^mAID^/Sororin^mAID^ cells through retroviral infection. In detail, the coding sequence of OsTIR2 was fused to that of TagBFP fluorescent protein and inserted into a MigR1 retroviral vector. 15μg of the MigR1-OsTIR2-TagBFP construct and 15μg of pCL-Eco packaging plasmid was co-transfected into HEK293T cells using PEI (Cat#23966, Polysciences). 48 and 72 hours later, virus-containing medium was collected and filtered through a 0.45μm filter to remove cell debris. To infect target cells, 3 million cells were plated per well into a 6-well plate with 1ml of culture medium. Then, 1ml of virus containing medium was added together with 8mg/ml polybrene and 10mM HEPES buffer (Gibco, Cat#15630-106). The plate was spun at 3000 rpm for 1.5 hours at room temperature. Cells were then washed and placed in fresh medium. 48 hours later, cells expressing TagBFP were sorted using a BD FACS AriaIII cell sorter.

### Generation of Sororin^9A^ mutant

The Sororin coding sequence was amplified from the cDNA library of parent G1E cells. Site-directed mutagenesis was performed by PCR with mutation containing primers. The mutated Sororin^9A^ coding sequence was then assembled and fused to GFP through Gibson assembly. A repetitive Flag sequence containing 5xFlag tags were then placed at the C terminal of the fusion protein. Sororin^9A^-GFP-5xFlag was then cloned into MigR1 retroviral vector. Virus packaging and transduction was performed as above described. To purify positively infected cells, Wapl^dTag^/Sororin^mAID^ and Wapl^dTag^/SMC2^mAID^/Sororin^mAID^ cells were transiently treated with dTag13 for 4 hours to eliminate the GFP signals from endogenous Wapl. Then, cells with positive GFP (Sororin^9A^) signals were purified using a BD FACS AriaIII cell sorter. Note, the GFP signals were maintained to be consistent among different clones.

### Cell growth curve

To examine the influence of Wapl or/and SMC2 depletion on cell growth, ∼50,000 Wapl^dTag^/SMC2^mAID^ cells were seeded in untreated non-adhesive 6 well plates in 2ml of fresh culture medium. Cells were then incubated with either 200nM of dTag13 (MCE Cat#HY-114421) or/and 1μM of 5-Ph-IAA or both to degrade Wapl or/and SMC2. Cells without any treatment were kept as control. Cell number of each condition was counted every 24 hours for a total of 72-hour duration. Three independent biological replicates were performed for both independent clones (clone1 and clone9).

### Quantification of lagging chromosomes in Wapl-deficient cells

To assess the impact of Wapl depletion on chromosome segregation during mitosis, Wapl^dTag^/SMC2^mAID^ cells (both clone1 and clone9) were treated either with or without dTag13 (200nM) for 4 hours. Meanwhile, cells were arrested at prometaphase by nocodazole treatment (200ng/ml, Sigma Cat#M1404). To trigger cell division, nocodazole arrested cells were washed once with fresh culture medium and released into nocodazole free medium with or without dTag13. One hour after nocodazole release, cells were fixed with 1% formaldehyde and stained with DAPI for 5min. Cells were then seeded into a glass-bottom 384-well plate and imaged with a Dragonfly CR-DFLY-202-2540 spinning disc confocal microscope equipped with 100 × objective lense (NA1.45). Lagging chromosome were manually identified in Wapl(+) and Wapl(-) dividing cells based on DAPI signals from “bridge-shaped” unresolved chromosomes. Over 100 dividing cells were examined for both conditions and three independent biological replicates were performed for both clones.

### Metaphase chromosome spread assay

Cells with indicated treatment were pelleted and re-suspended in 0.075M KCl and incubated at 37℃ for 15min. Cells were then pelleted at room temperature at 200g for 5min and re-suspended in 5 ml of fresh Carnoy’s Fixative (3:1 ratio of methanol: acetic acid). Fixation was performed twice. Cells were pelleted and re-suspended in 200μl of fresh Carnoy’s Fixative. Drop the cells from 0.5 meter above onto a slide tilted with a 45° angle. The slides chromosome spreads were immediately dried over the flames of an alcohol burner and stained Giemsa solution (Solarbio, Cat#G1015) for 8min. Slides were then washed by running water and air dried. The slides were then mounted for long term storage and imaging. Chromosome spreads were imaged through an Olympus SLIDEVIEW VS200 slide scanner. Quantification of the sister-chromatid dissociation status and the chromosome length were manually performed.

### Quantification of mitotic chromosome length

Images were visualized in OLYMPUS OlyVIA software (4.1). Chromosome lengths were measured along the outline of the center of the chromosomes using a point-by-point tracing tool at 160x magnification. Thirty cells were randomly selected for condition and all chromosomes in the cells were measured. Data were exported by OLYMPUS OlyVIA software.

### Immunofluorescent staining

Cells with indicated treatment were fixed with 4% formaldehyde in 1×PBS for 5min at room temperature. Fixation were quenched by 10mM Tris-HCl (ph. 7.5, Sangon Biotech Cat#B548139) at room temperature for 5min. Cells were then permeabilized with 0.2% Triton X-100 (Diamond Cat#A110694) for 5min at room temperature. Cells were then washed with 1×PBS twice and re-suspended in 250µl of blocking buffer (2% BSA, 0.1% Tween in 1×PBS) for 30min at room temperature. After blocking, cells were pelleted and re-suspended in 100µl of blocking buffer containing diluted primary antibody and incubated at room temperature for 3 hours. Cells were then washed twice with 1×PBS and re-suspended in 100µl of blocking buffer containing diluted secondary antibody. Secondary antibody incubation was performed for 1 hour at room temperature. Cells were then pelleted and re-suspended in blocking buffer containing DAPI (4μg/ml) and seeded into glass bottom 384-well plates. Primary antibody involved in this experiment is: anti-Rad21 (Millipore, 1:50).

### Western blotting

Western blot was performed as previously described. 5×10^6^ cells were lysed with RIPA buffer (Beyotime, p0013b) supplemented with 1:500 protease inhibitor (Sigma-Aldrich, P8340) on ice for 20 min. Cell lysates were then subjected to sonication for 5 min (40% amplitude, 15 s ON and 15 s OFF, 13 min sonication) using a QSonica Q800R3 sonicator. Blots were probed with antibodies against SMC2 (Abclonal, A17867, 1:1000), Wapl (CST, 77428, 1:1000) and β-actin (ABclonal, AC004; 1:5,000) and imaged using a BioRad ChemiDoc MP Imaging System.

### Nocodazole mediated prometaphase arrest

To obtain mitotic chromosomes loaded with cohesin (extrusive-plus cohesive-cohesin). Asynchronous Wapl^dTag^/SMC2^mAID^ cells were treated with dTag13 either with or without 5-Ph-IAA for 4 hours. The cells were then incubated with Nocodazole for 4 hours for prometaphase enrichment before crosslinking and FACS sorting. As a control, Wapl^dTag^/SMC2^mAID^ cells without dTag13 treatment were processed in parallel.

### G2/M arrest/release protocols (original)

To obtain mitotic chromosomes with all eight possible configurations of extrusive-cohesin, cohesive-cohesin and condensin, we devised a series of G2/M arrest/release strategies.

*Configuration 1: condensin(+); extrusive-cohesin (-); cohesive-cohesin (-).* Asynchronously growing Wapl^dTag^/Sororin^mAID^ cells were treated with the CDK1 inhibitor RO-3306 (10μM) for G2-arrest. Meanwhile, 5-Ph-IAA (1μM) was added to the culture to degrade Sororin. 16 hours later, Nocodazole (200ng/ml) was then added to the culture for another 4 hours. Then, cells were washed with 1xPBS for 3 times and released into fresh medium containing 5-Ph-IAA and Nocodazole but not RO-3306 to allow mitotic progression. Prometaphase cells were further enriched for 4 hours before crosslinking and FACS sorting.

*Configuration 2: condensin(+); extrusive-cohesin (+); cohesive-cohesin (-).* Asynchronously growing Wapl^dTag^/Sororin^mAID^ cells were treated with the CDK1 inhibitor RO-3306 (10μM) for G2-arrest. Meanwhile, 5-Ph-IAA (1μM) was added to the culture to degrade Sororin. A 16-hour G2-arrest was sufficient for Wapl to unload the majority of cohesive-cohesin. Then, dTag13 (200nM) was added into the culture to degrade Wapl and allow extrusive-cohesin to accumulate. Nocodazole (200ng/ml) was added into the culture at the same time. 4 hours later, cells were washed with 1xPBS for 3 times and released into fresh medium containing 5-Ph-IAA, dTag13 and Nocodazole but not RO-3306 to allow mitotic progression. Prometaphase cells were further enriched for 4 hours before crosslinking and FACS sorting.

*Configuration 3: condensin(+); extrusive-cohesin (-); cohesive-cohesin (+).* The exact same protocol for configuration 1 was applied to Wapl^dTag^/Sororin^mAID^ cells ectopically express the Sororin^9A^ mutant. In this setup, Wapl was maintained to unload extrusive-cohesin during mitotic progression. Meanwhile, cohesive-cohesin was safeguarded by the Sororin^9A^ mutant against Wapl and therefore retained during mitosis.

*Configuration 4: condensin(+); extrusive-cohesin (+); cohesive-cohesin (+).* The exact same protocol for configuration 2 was applied to Wapl^dTag^/Sororin^mAID^ cells ectopically expressing the Sororin^9A^ mutant.

*Configuration 5: condensin(-); extrusive-cohesin (-); cohesive-cohesin (-).* The exact same protocol for configuration 1 was applied to Wapl^dTag^/SMC2^mAID^/Sororin^mAID^ cells. Condensin was simultaneously degraded with Sororin through 5-Ph-IAA treatment.

*Configuration 6: condensin(-); extrusive-cohesin (+); cohesive-cohesin (-).* The exact same protocol for configuration 2 was applied to Wapl^dTag^/SMC2^mAID^/Sororin^mAID^ cells.

*Configuration 7: condensin(-); extrusive-cohesin (-); cohesive-cohesin (+).* The exact same protocol for configuration 1 was applied to Wapl^dTag^/SMC2^mAID^/Sororin^mAID^ cells ectopically expressing the Sororin^9A^ mutant.

*Configuration 8: condensin(-); extrusive-cohesin (+); cohesive-cohesin (+).* The exact same protocol for configuration 2 was applied to Wapl^dTag^/SMC2^mAID^/Sororin^mAID^ cells ectopically expressing the Sororin^9A^ mutant.

Configuration 9: *condensin(-); extrusive-cohesin (+); cohesive-cohesin (+, mediated by endogenous Sororin).* The exact same protocol for configuration 2 was applied to Wapl^dTag^/SMC2^mAID^ cells. In these cells, cohesive-cohesin was safeguarded by endogenous Sororin.

### G2/M arrest/release protocol (extrusive-cohesin light load)

To reduce the amount of extrusive-cohesin loaded on mitotic chromosomes, we modified the protocol for configuration 2 by reducing the duration of dTag13 treatment from 4 hours to 45 minutes. This protocol can be applied to Wapl^dTag^/Sororin^mAID^ or Wapl^dTag^/SMC2^mAID^/Sororin^mAID^ cells either in their original form or ectopically expressing the Sororin^9A^ mutant, allowing us to obtain the “extrusive-cohesin light load” version of configurations 2, 4, 6 and 8.

### G2/M arrest/release protocol (short-arrest and release)

To account for the possibility that Sororin^9A^ mutant protected cohesive-cohesin may still be unloaded by Wapl during the 4-hour course of prometaphase enrichment, thereby mitigating the potential effect of cohesive-cohesin, we designed a modified G2/M and short arrest and release protocol. In detail, asynchronous cells (Wapl^dTag^/Sororin^mAID^ or Wapl^dTag^/SMC2^mAID^/Sororin^mAID^ cells ectopically expressing Sororin^9A^ mutant) were treated with RO-3306 for 4 hours for G2-arrest. Meanwhile, 5-Ph-IAA was added to degrade endogenous Sororin. Nocodazole was then added for another 4 hours. Cells were then washed with 1xPBS for 3 times and released into fresh medium containing 5-Ph-IAA and Nocodazole but not RO-3306 to allow mitotic progression. Prometaphase cells were further enriched for 30 minutes before crosslinking and FACS sorting. In this modified protocol, the total duration in RO-3306 was shortened from 20 hours in the original protocol to 8 hours. Meanwhile, the duration of mitotic enrichment was shortened from 4 hours to 30min. These modifications aimed to reduce the exposure of cohesive-cohesin to Wapl. As controls, Wapl^dTag^/Sororin^mAID^ or Wapl^dTag^/SMC2^mAID^/Sororin^mAID^ cells without Sororin^9A^ mutant were subject to the exact same protocol to obtain mitotic chromosomes without cohesive-cohesin.

### Fluorescence activated cell sorting (FACS)

To obtain mitotic populations, samples that underwent either the nocodazole mediated prometaphase arrest or the original or modified G2/M arrest/release protocols were crosslinked with formaldehyde (1% for ChIP-seq experiments and 2% for Hi-C experiments) for 10min at room temperature. Crosslinking was stopped by glycine (Sangon Biotech Cat#A100167) with a final concentration of 1M. Cells were then permeabilized by 0.1% Triton X-100 for 5min. All cells were then stained with anti-pMPM2 antibody (Millipore Cat#05-368, 0.2µl/10million cells) for 50min at room temperature, followed by APC-conjugated F(ab’)2-goat anti-mouse secondary antibody (Thermo Fisher Scientific, Cat#17-4010-82, 2µl/10million cells) for 45min at room temperature. After staining, cells were re-suspended in 1× FACS buffer (1× PBS containing 2mM EDTA and 2% FBS) at a density of 100 million cells per ml. To enrich mitotic population, cells were sorted based on pMPM2 signal (+) and DAPI signal (4N) via a BD FACS AriaIII cell sorter. 2 million cells were collected per ChIP experiment per condition.

To enrich G1-phase control cells, asynchronous Wapl^dTag^/SMC2^mAID^ cells were treated either with or without dTag13 for 4 hours. The cells were then sorted based on DAPI (2N) and GFP signals to enrich G1-phase populations either with or without Wapl. Roughly 7-8 million interphase cells were collected per ChIP experiment per condition. Protease inhibitor (1:500 Sigma, P8340) and PMSF (Sangon Biotech Cat#A100754, 1:100) were added to all buffers during the entire sample preparation procedure. Sorted cells were pelleted, snap-frozen through liquid nitrogen and stored at −80℃ until further use.

### ChIP-seq

Chromatin immunoprecipitation (ChIP) was performed as previously described, using antibodies against CTCF (Millipore Cat#07-729, 7.5μl/IP), Rad21 (Abcam Cat#ab992, 5μl/IP) and Flag (Sigma, Cat#M2) respectively. Briefly, FACS purified cells were re-suspended in 1ml cold Cell Lysis Buffer (10mM Tris pH 8, 10mM NaCl and 0.2% NP-40) supplemented with 1:500 protease inhibitor and 1:100 PMSF for 20min on ice. Meanwhile, roughly 30% of crosslinked 293T cells were added into the cell mixture. Nuclei were then pelleted and re-suspended in 1ml Nuclear Lysis Buffer (50mM Tris pH 8, 10mM EDTA, 1% SDS, fresh supplemented with PI and PMSF) for 20min on ice. Chromatin was then fragmented by sonication (Qsonica Q800R3, 80% amplitude, 20s ON, 40s OFF) for 17min. Cell debris were removed by centrifugation at 15000g for 10min at 4℃. The supernatant was collected and diluted with 4 volumes of IP dilution buffer (20mM Tris pH 8.0, 2mM EDTA, 150mM NaCl, 1% Triton X-100, 0.01% SDS) supplemented with protease inhibitor, PMSF and 50μl of A/G agarose beads (Santa Cruz Cat#sc-2003). Preclear of chromatin was performed at 4℃ for 4-8h with rotation. Precleared chromatin was then incubated with 35µl A/G agarose beads (A:G = 1:1) pre-bound with antibody and incubated on a rotator at 4℃ overnight. Beads were washed once with IP wash buffer I (20mM Tris pH 8, 2mM EDTA, 50mMNaCl, 1% Triton X-100, 0.1% SDS), twice with high salt buffer (20mMTris pH 8, 2mM EDTA, 500mMNaCl, 1% Triton X-100, 0.01% SDS), once with IP wash buffer II (10mMTris pH 8, 1mM EDTA, 0.25 M LiCl, 1% NP-40/Igepal, 1% sodium deoxycholate) and twice with TE buffer (10mM Tris pH 8, 1mM EDTA pH 8). All washing steps were performed on ice. Beads were moved to room temperature and DNA was eluted in 200µl fresh Elution Buffer (100mM NaHCO3, 1%SDS). ChIP and input chromatin were reverse crosslinked at 65℃ overnight in the presence of proteinase K. IP and input samples were supplemented with 10ul of 3M sodium acetate (pH 5.2), and DNA was purified with PCR purification kit (QIAGEN Cat#28106). ChIP-seq libraries were constructed using the VAHTS Universal DNA Library Prep Kit for MGI (Vazyme, Cat#NDM607-02) according to the manufacture’s protocol. ChIP libraries were sequenced on the MGI DNBSEQ-T7 sequencing platform.

### In-situ Hi-C

In-situ Hi-C was performed as previously described. In brief, FACS purified cells (0.1 million) were lysed in 1ml cold Cell Lysis Buffer (10mM Tris pH 8, 10mM NaCl, 0.2% NP-40) for 10min on ice. Nuclei were pelleted at 7200rpm for 2min at 4℃ and washed once with cold 1.4× NEB buffer 3.1 (NEB Cat#B7203S). Nuclei were then re-suspended in 25μl of 1.4× NEB buffer 3.1 containing 0.1% SDS and incubated at 65℃ for 10min. Triton X-100 was then added into the system to a final concentration of 1% to quench SDS. Restriction digestion was performed by adding 25U DpnII restriction enzyme (NEB, Cat#R0543M) and incubating the mixture at 37℃ overnight with shaking (950rpm on a thermal mixer). After overnight digestion, an additional stroke of 25U DpnII was added into the system to further digest the genome for 4 hours at 37℃. Nuclei were incubated at 65℃ for 20min to inactivate DpnII. Digested chromatin fragments were then cooled down to room temperature and repaired with dCTP, dGTP, dTTP (Diamond Cat#B110049, B110050, B110051) and Biotin-14-dATP (active motif, Catalog No: 14138) using 6.6U DNA Polymerase I, Large (Klenow) fragment (NEB, Cat#M0210). Chromatin was then ligated with 2000U T4 DNA ligase (NEB, Cat#M0202M) for 4 hours at 16℃ followed by further incubation for 4 hours at room temperature. Reverse-crosslinking was then performed overnight at 65℃ in the presence of 1% SDS and proteinase K (G-clone Cat#EZ0970). After reverse-crosslinking, DNase-free RNase A (Vazyme Cat#DE111-01) was added to the system for 30min at 37℃ to remove RNA. DNA was then extracted by phenol-chloroform extraction, precipitated, and dissolved in 130μl of nuclease free water. DNA was fragmented with a QSonica Q800R3 sonicator (40% amplitude, 15s ON and 15s OFF, 13min sonication) and purified with 1.8x VAHTS DNA Clean Beads (Vazyme Cat#N411). Biotin-labeled DNA fragments was enriched by incubation with 50µl Dynabeads MyOne Streptavidin C1 beads (Thermo Fisher Scientific, Cat#65002) at room temperature for 15min. Streptavidin C1 beads were then washed twice with 1 × B&W buffer (5mM Tris-HCl, pH 7.5; 0.5mM EDTA; 1M NaCl). Beads were then washed twice with 1 × B&W buffer and subject to library construction. DNA end repair, dA-tailing and adaptor ligation were constructed on beads, using the VAHTS Universal DNA Library Prep Kit for Illumina or MGI (Vazyme, Cat# ND627-01, NDM607-02) based on the manufacture’s protocol. To eluted DNA from streptavidin beads, the library mixture was treated with 0.1% SDS and incubated at 98℃ for 10min. Released DNA was collected from the supernatant and purified with 1.8x VAHTS DNA Clean Beads. Finally, the library was amplified for 9 cycles on a thermal cycler, using the VAHTS® HiFi Amplification Mix. PCR products were then purified with 1.8x VAHTS DNA Clean Beads and sequenced on MGI DNBSEQ-T7 sequencing platform.

### Quantification and data analysis

#### ChIP-seq data processing

Conventional ChIP-seq processing pipeline were adopted. Briefly, paired-end reads were aligned to mouse reference genome mm9 through the Bowtie2 (v2.3.5.1) software with default parameters and soft clipping allowed (“--local”). All unmapped reads and low quality alignments with MAPQ score lower than 30 was removed using SAMtools (v1.9). PCR duplicates were removed using Picard (v2.23.3) with default parameters. Reads aligned to mitochondria, random contigs and ENCODE blacklisted regions were also removed using BEDtools (v2.27.1)^44^. Counts per million mapped reads (CPM) normalization was performed using the bamCoverage function of Deeptools (v3.1.3).

### Peak calling

#### CTCF ChIP-seq

CTCF ChIP-seq were performed on Wapl^dTag^/SMC2^mAID^ cells. Peak calling was performed on individual biological replicates using MACS2 (v2.2.7.1) with default parameters and a 0.05 q-value cutoff for narrow peaks. Corresponding input sample was adopted as background controls for MACS2 peak calling. To ensure the validity of peaks, we employed stringent peak identification criterion:

- interphase Wapl(+): peaks present in 2 out of 2 biological replicates.
- interphase Wapl(-): peaks present in 2 out of 2 biological replicates.
- mitosis condensin(-),Wapl(-): peaks present in ≥2 out of 4 biological replicate.
- mitosis condensin(+),Wapl(-): peaks present in ≥2 out of 4 biological replicates.

#### Rad21 ChIP-seq

Rad21 ChIP-seq were performed on Wapl^dTag^/SMC2^mAID^, Wapl^dTag^/Sororin^mAID^ and Wapl^dTag^/SMC2^mAID^/Sororin^mAID^ cells. Peak calling was performed on individual biological replicates using MACS2 (v2.2.7.1) with default parameters and a 0.05 q-value cutoff for the first cell line and 1×10^-6^ q-value cutoff for the last two cell lines for narrow peaks. Input samples corresponding to SMC2(+) or SMC2(-) conditions from *Wapl^dTag^/SMC2^mAID^*cells were used as background. We implemented stringent peak identification criterion as below:

*For Wapl^dTag^/SMC2^mAID^ cells:*

- mitosis condensin(-),Wapl(-): peaks present in ≥3 out of 5 biological replicates.
- mitosis condensin(+),Wapl(-): peaks present in ≥3 out of 5 biological replicates.

*For Wapl^dTag^/SMC2^mAID^/Sororin^mAID^ cells:*

- mitosis condensin(-),extrusive-cohesin(+),cohesive-cohesin(-)(extrusive-cohesin heavy load): peaks present in ≥2 out of 5 biological replicates.
- mitosis condensin(-),extrusive-cohesin(+),cohesive-cohesin(-)(extrusive-cohesin light load): peaks present in ≥1 out of 2 biological replicates.
- mitosis condensin(-),extrusive-cohesin(+),cohesive-cohesin(+)(extrusive-cohesin heavy load): peaks present in ≥2 out of 4 biological replicates.
- mitosis condensin(-),extrusive-cohesin(+),cohesive-cohesin(+)(extrusive-cohesin light load): peaks present in ≥2 out of 3 biological replicates.
- mitosis condensin(-),extrusive-cohesin(-),cohesive-cohesin(+): peaks present in ≥2 out of 4 biological replicates.
- mitosis condensin(-),extrusive-cohesin(-),cohesive-cohesin(+)(short-arrest and release protocol): peaks present in ≥2 out of 3 biological replicates.

*For Wapl^dTag^/Sororin^mAID^ cells:*

- mitosis condensin(+),extrusive-cohesin(+),cohesive-cohesin(-)(extrusive-cohesin heavy load): peaks present in ≥2 out of 5 biological replicates.
- mitosis condensin(+),extrusive-cohesin(+),cohesive-cohesin(-)(extrusive-cohesin light load): peaks present in ≥1 out of 2 biological replicates.
- mitosis condensin(+),extrusive-cohesin(+),cohesive-cohesin(+)(extrusive-cohesin heavy load): peaks present in ≥2 out of 5 biological replicates.
- mitosis condensin(+),extrusive-cohesin(+),cohesive-cohesin(+)(extrusive-cohesin light load): peaks present in ≥2 out of 4 biological replicates.

#### Sororin^9A^-Flag ChIP-seq

*Sororin^9A^-Flag* ChIP-seq were performed on Wapl^dTag^/Sororin^mAID^ and Wapl^dTag^/SMC2^mAID^/Sororin^mAID^ cells. Peak calling was performed on individual biological replicates using MACS2 (v2.2.7.1) with default parameters and a 0.05 q-value cutoff for narrow peaks. Input samples corresponding to SMC2(+) or SMC2(-) conditions from *Wapl^dTag^/SMC2^mAID^*cells were used as background. We implemented stringent peak identification criterion as below:

*For Wapl^dTag^/SMC2^mAID^/Sororin^mAID^ cells:*

- mitosis condensin(-), extrusive-cohesin(+),cohesive-cohesin(+)(extrusive-cohesin heavy load): peaks present in ≥2 out of 5 biological replicates.
- mitosis condensin(-), extrusive-cohesin(+),cohesive-cohesin(+)(extrusive-cohesin light load): peaks present in 1 out of 1 biological replicates.
- mitosis condensin(-), extrusive-cohesin(-),cohesive-cohesin(+): peaks present in ≥1 out of 2 biological replicates.
- mitosis condensin(-), extrusive-cohesin(-),cohesive-cohesin(+)(short-arrest and release): peaks present in ≥2 out of 3 biological replicates.

*For Wapl^dTag^/Sororin^mAID^ cells:*

- mitosis condensin(+),extrusive-cohesin(+),cohesive-cohesin(+)(extrusive-cohesin heavy load): peaks present in ≥4 out of 8 biological replicates.
- mitosis condensin(+),extrusive-cohesin(+),cohesive-cohesin(+)(extrusive-cohesin light load): peaks present in ≥2 out of 4 biological replicates.

For peak annotation, Rad21 or Sororin^9A^-Flag peaks were intersected with the overall CTCF peak from this study and the cis-regulatory element from previous studies ^20^. For all ChIP-seq experiments, signal intensity of CTCF, Rad21 or Sororin^9A^-Flag peaks were obtained through the UCSC utility: BigWigAverageOverBed. Heatmap profiles of CTCF, Rad21 and Sororin^9A^-Flag were generated through Deeptools. Genome browser track of ChIP-seq results were generated using Pygenometracks (v3.7).

#### Identification of cohesin islands

We observed large patches of regions with elevated Rad21 ChIP-seq signals in the condensin-deficient mitotic cells with extrusive-cohesin (configuration 6). We noticed that these patches may contain multiple MACS2 identified Rad21 peaks. We thus merged the Rad21 ChIP-seq peaks with a minimal distance of 20kb using the *merge* function of bed tools: We defined cohesin islands as below, a cohesin island must contain > 10 MACS2 identified Rad21 peaks. In this way, we identified 382 cohesin islands that we absent in interphase cells.

#### Hi-C data preprocessing

*In-situ* Hi-C from all biological replicates were independently aligned to mouse reference genome mm9 using bowite2 (global parameters: --very-sensitive -L 30 --score-min L, −0.6,-0.2 --end-to-end –reorder; local parameters: --very-sensitive -L 20 --score-min L,-0.6,-0.2 --end-to-end --reorder). Hi-C pro software (v3.0.0) were employed to generate valid interaction pair files. The output pair files were then converted to “.hic” files using the hicpro2juicebox utility. HiC files were also converted to “.cool” files using the Hicexplorer (v3.7). For replicate or clone-merged samples, similar steps were taken on reads merged from each individual biological replicate or clones.

### *Trans*-interactions

The percentage of trans-chromosomal interactions was adapted from the HiC Pro statistic summary.

#### Contact probability decay curves (*P(s)* curves)

The *P(s)* curve of each individual chromosome for each sample was generated using the “expected-cis” function of Cooltools (v0.4.0). The genome-wide *P(s)* curve was computed by averaging individual chromosomes together.

#### Insulation score profiling and boundary calls

Insulation scores were computed on 10kb binned, KR balanced contact matrices, using the “insulation” function of Cooltools (v0.4.0). The sliding window was set to be 12bin x 12bin as we previously practiced ^20^. Bins with a threshold of over 0.2 were selected as the tentative boundaries for each sample. This procedure was applied to interphase cells with or without Wapl as well as mitotic cells of configurations 1 to 8. Note that the “extrusive-cohesin light load” version for configurations 2, 4, 6 and 8 were also included.

#### Domain identification

To identify domains, we combined the domain calling algorithm “Arrowhead” and insulation score profiles. To begin with, we generated an overall non-redundant list of boundaries using boundary files across all samples. Boundaries from each sample were merged together with a minimal gap distance of 50kb using the *merge* function of bedtools. To this end, we obtained 13,227 non-redundant boundaries across all processed samples. To identify domains, we applied the domain calling algorithm “Arrowhead” to all of the above mentioned samples. Domain calls were independently performed on 10kb, 25kb and 50kb binned Hi-C contact matrices for all samples. For a given sample, all identified domains were then collected together to generate an initial domain list. To eliminate spurious domains, for each sample, we intersected the initial domain list with the above acquired non-redundant boundary list. In detail, boundaries from each chromosome were randomly paired up so that the size of each boundary pair was greater than 50kb and smaller than 10Mb. Each pair of boundary represents a potential domain. For a given sample, the initial domain list was intersected with the overall boundary pairs. For a domain to be considered as a real domain, it must overlap with a boundary pair with a wiggling within of 50kb for each boundary. This procedure was applied to all interphase and mitotic samples.

#### Domain partition

We partitioned all domains into TADs, compartmental domains and others. For a domain to be classified as a TAD, it must fulfill two criteria: (1) the intra-domain contact frequency (observed/expected) must be at least 1.2 fold higher than that of the condensin-deficient mitotic samples without any retained cohesin (configuration 5). This criterion was chosen because we previously demonstrated that condensin-deficient mitotic chromosomes did not form CTCF/cohesin mediated TADs. (2) The domain must contain CTCF peaks at both boundaries. The intersection rules are listed as below: <1> For domains < 1Mb, a wiggle room of 15kb from the center of each boundary was used. <2> For domains between 1 to 2Mb, a wiggle room of 25kb from the center of each boundary was used. <3> For domains between 2 to 3Mb, a wiggle room of 50kb from the center of each boundary was used. <4> For domains between 3 to 4Mb, a wiggle room of 100kb from the center of each boundary was used. <5> For domains > 4Mb, a wiggle room of 150kb from the center of each boundary was used.

For a domain to be classified as a compartmental domain, it must fulfill two criteria: (1) the intra-domain contact frequency (observed/expected) must be < 0.8 fold of that of the condensin-deficient mitotic samples without any retained cohesin (configuration 5). (2) it may bear CTCF peaks in at most one boundary.

#### Domain segmentation

To determine the influence of cohesive-cohesin on TADs, we segmented all identified TADs into 100kb-wide stripes that are parallel to the diagonal. The maximal distance between the stripe to the diagonal was set to be 5Mb. For a given sample, the average observed/expected value within each stripe was computed and denoted as the stripe intensity *SI_j_* (j denotes the distance between the stripe and the diagonal). The log_2_ fold change of *SI_j_* between cohesive-cohesin (+) and cohesive-cohesin (-) samples were then computed for each TAD. TAD with different size range were grouped together and plotted to generate the distance dependent stripe intensity fold change line graph.

#### Loop calling and post-processing

Chromatin loops were identified on interphase and mitotic samples using the previously described HICCUPS method with modifications. (1). Loop identification: For a given sample preliminary loops were called at 10kb resolution (p: 4bins, i: 16bins, FDR: 0.2) or 25kb resolution (p: 5bins, i: 15bins, FDR: 0.1) using the HICCUPS function of juicer_tools (v1.22.01). This procedure was applied to interphase cells with or without Wapl as well as mitotic cells of configurations 1 to 8. Note that the “extrusive-cohesin light load” version for configurations 2, 4, 6 and 8 were also included. (2) Loop merging (10kb and 25kb loops were merged separately): Two identical or nearby (next to each other) pixels from different samples may be redundantly recorded as two separate loops instead of one. Therefore, a merging step based on the greedy algorithm to unify closely spaced clustering loops was implemented. Briefly, for a given loop in the overall full list from step (1), a value *q*_min_ which represents the minimal *q* value across all interphase and mitotic samples was recorded. All loops were then reordered in an ascending order based on their *q*_min_, so that loops at the top of the list were the most confidant calls. We then scanned through the entire list to identify pixels that fell within a 20kb radius of the top pixel. If no additional nearby pixels were identified, the top pixel was then considered to be a single pixel by itself without any clustering. If nearby pixels were identified, then the top pixel and nearby pixels were together considered as a pixel cluster. Pixels in the cluster was removed from the full loop list and the centroid of the pixel cluster was re-identified. A distance s was calculated to be the distance from the centroid to the far-border of the pixel cluster. We then, started a second round of scanning in the rest of the full loop list to identify pixels that fell within a radius of 20kb + *s* from the pixel cluster centroid. Pixels within the loop cluster after the second merging step were removed from the full loop list. We then, focused again on the top pixel of the remaining full loop list and repeated the above merging steps until no further pixels were left in the full loop list. After iterating through all loops, we generated a list of loop clusters which may contain one or more pixels. For each loop cluster, a cluster summit was identified that represented the pixel with the lowest *q*_min_. The resulted list of loop summits represents the final non-redundant loops for a specific bin resolution (10kb or 25kb). (3) Merging 10kb and 25kb loops. Non-redundant loop lists from 10kb and 25kb binned matrices were merged together and if a 25kb loop cluster overlapped with a 10kb loop cluster, then the 25kb loop was removed from the list. (4) Removal of spurious loops. We made an assumption that no loop was present in the condensin-replete mitotic samples without any cohesin (configuration 1). Therefore, we implemented a loop strength filter. For a given loop summit, we computed the observed/donut-expected values of the summit itself and eight surrounding pixels. The average value of the nine pixels were denoted as the loop strength. For a loop to be considered real, its strength must be at least 1.5 fold higher than that of the condensin-replete control in at least one of the remaining samples.

#### Structural loop partition and clustering

For a loop to be considered as a structural loop in a broad sense, both of its anchors must contain both CTCF and Rad21 peaks. For a ChIP-seq peak to intersect with loop anchor, it must have at least 1bp overlap with a 30kb region centered around the midpoint of the loop anchor summit. In this way, we identified 26,086 structural loops. To interrogate the influence of cohesive-cohesin on extrusive-cohesin mediated structural loops, we performed unsupervised *k-means* clustering on the 9,027 structural loops identified in the condensin-deficient mitotic samples with heavily loaded extrusive-cohesin (configuration 6). We identified 4 major clusters of structural loops with distinct sensitivity to cohesive-cohesin.

#### Loop anchor partition

To assess the properties of structural loop anchors for a specific cluster, we combined all upstream and downstream anchors of the 9,027 structural loops identified in the condensin-deficient mitotic samples with heavily loaded extrusive-cohesin (configuration 6). The anchors were merged to a unified non-redundant anchor list using the *merge* function of bedtools. Each unique anchor was then intersected with the structural loop list to determine if it belongs to a specific cluster of loops or shared by multiple cluster of loops. Only cluster-specific anchors were processed for downstream analysis.

#### CRE contacts identification and partition

We employed a previously described strategy to identify CRE contacts in interphase and mitotic samples. Briefly, 10kb bins containing CREs on the same chromosome were randomly paired up with a selected range of (70 to 300kb). All CRE pairs were then subject to HICCUPS (-p: 4, -i:16) to calculate the observed/donut-expected values for all interphase and mitotic samples. Putative CRE pairs were then subject to the below filters. For a given sample, it must (1) has an observed/expected value of >2 and (2) has an observed/expected value that is over two fold of that of the condensin-replete mitotic control cells (configuration 1), to be considered as a real CRE contact. We then summarized the CRE contacts that were positive in at least one of the following samples: (1) Wapl-replete interphase, (2) Wapl-deficient interphase, (3) configuration 5 of mitosis, (4) configuration 6 (either heavy or light loaded) of mitosis, (5) configuration 7 of mitosis and (6) configuration 8 (either heavy or light load) of mitosis. Finally, CRE pairs with both anchors containing CTCF/cohesin co-occupied peaks were removed. To this end, we identified 11,399 CRE contacts.

To assess the influence of structural loop anchors on CRE contact strength, we stratified the CRE contacts based on the below criteria: (1) a CRE contact must not be encompassed by a structural loop, (2) a CRE contact may encompass either 0, 1, 2 or >2 structural loop anchors.

#### Compartmentalization related analysis

For each sample, eigenvector decomposition on the Pearson’s correlation matrix of the observed/expected values of 25kb binned Knight-Ruiz (KR) balanced cis-interaction matrices using the “eigs-cis” function of Cooltools (v0.4.0) ^45^. EV1 direction was assigned based on gene density. Saddle plots or aggregation-repulsion plots was generated as previously described by rearranging the rows and columns of the Hi-C contact matrices following certain rules. For saddle plots, the rows and columns were relocated based on the EV1 value of interphase samples. For aggregation-repulsion plots, rows and columns were relocated so that bins belonging to the same mitotic specific compartment were grouped together^24^.

#### Aggregated plots for loops and domains

Aggregated plots were generated using the python package Coolpup (v0.9.7) ^46^. For unscaled aggregated peak analysis (APA), loops smaller than 70kb were removed from the plots to avoid influence from pixels close to the diagonal.

## Data availability

The Hi-C and ChIP–seq data generated in this study are deposited at the GEO database with the accession number GSE269952. External ChIP–seq data from previous studies are listed as follows: H3K27ac (GSE61349) ^47^, H3K4me1 (GSM946535), H3K4me3 (GSM946533), H3K36me3 (GSM946529), H3K9me3 (GSM946542), H3K27me3 (GSM946531) ^48^ and CTCF (GSE98671) ^49^. External Hi-C data of Condensin I or condensin II deleted mitotic cells are available at GSE228402 ^24^. External ChIP-seq data of mitotically retained CTCF are available at GSE129997 ^20^.

## Code availability

Code will be available upon requests.

---

**Extended Data Figure 1.**
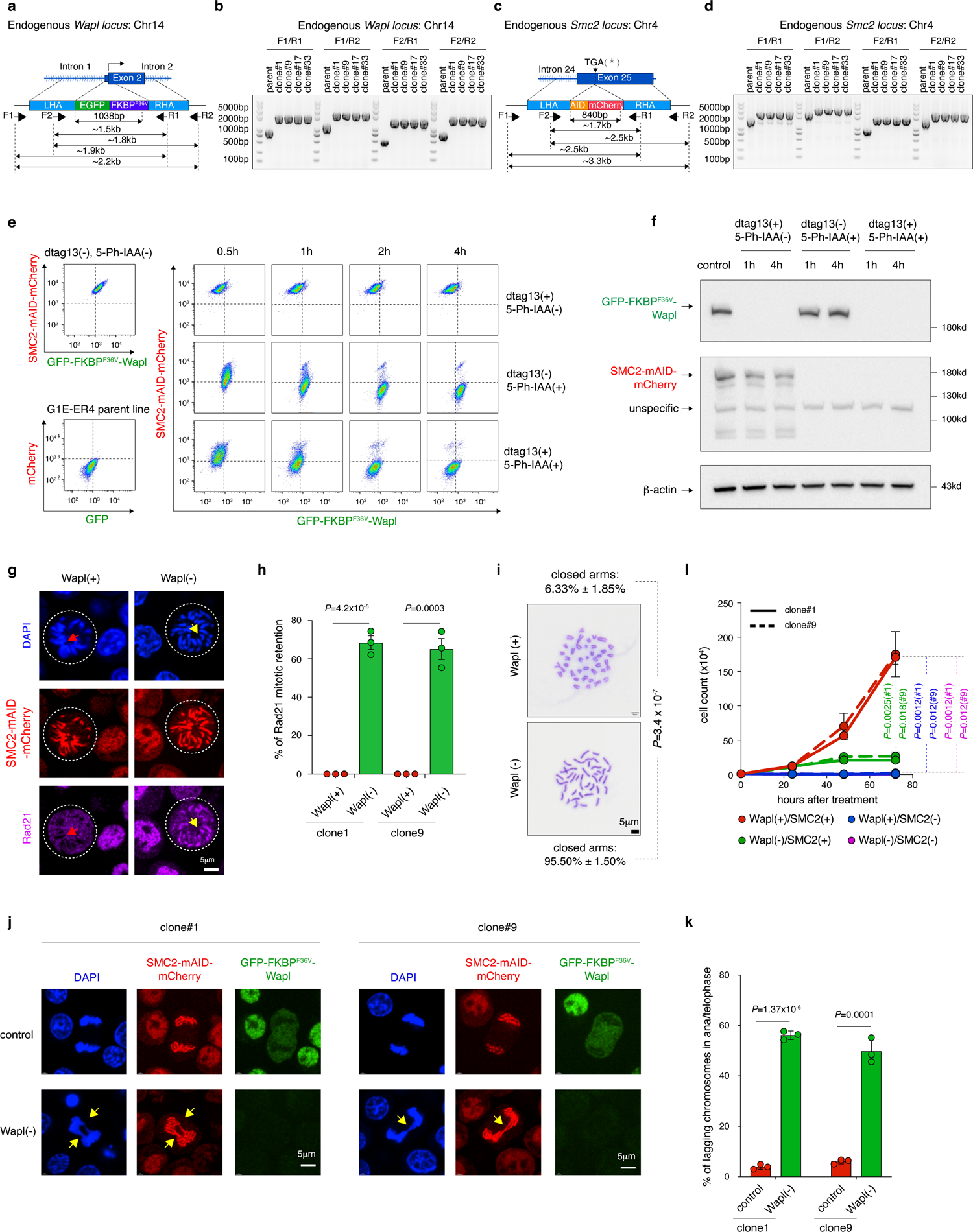
Characterization of the Wapl^dTag^/SMC2^mAID^ cell line. **a**, Schematic showing the edited *Wapl* locus and genotyping strategy. **b**, Genotyping results of positive clones targeting the *Wapl* locus. **c**, Schematic showing the edited *Smc2* locus and genotyping strategy. **d**, Genotyping results of positive clones targeting the *Smc2* locus. **e**, Flow cytometry plots showing progressive loss of Wapl or/and SMC2 fluorescence signals upon dTag13 or/and 5-Ph-IAA treatment. The parental cell line was shown as negative control. **f**, Western blots showing degradation of Wapl or/and SMC2 upon dTag13 or/and 5-Ph-IAA treatment. **g**, Immunofluorescence image showing mitotic retention of Rad21 upon Wapl depletion. Scale bar: 5μm. **h**, Quantification of the percentage of mitotic Rad21 retention upon Wapl loss (n=3 for each clone). Error bar denotes SEM. *P* values were calculated using two-sided student’s *t*-test. **i**, Metaphase spread analysis showing “closed-arm” mitotic cells in the absence of Wapl, indicating mitotic retention of cohesin. Scale bar: 5μm. *P* values were calculated using two-sided student’s *t*-test. **j**, Representative image showing the presence of lagging chromosomes in ana/telophase upon Wapl depletion. Scale bar: 5μm. Two independent clones were shown. **k**, Quantification of lagging chromosomes in Wapl-replete and depleted mitotic cells (n=3 for each clone). Error bar denotes SEM. *P* values were calculated using two-sided student’s *t*-test. **l**, Line graph showing cell growth defects in Wapl depleted, SMC2 depleted or Wapl and SMC2 co-depleted cells. Error bar denotes SEM. *P* values were calculated using two-sided student’s *t*-test.

**Extended Data Figure 2.**
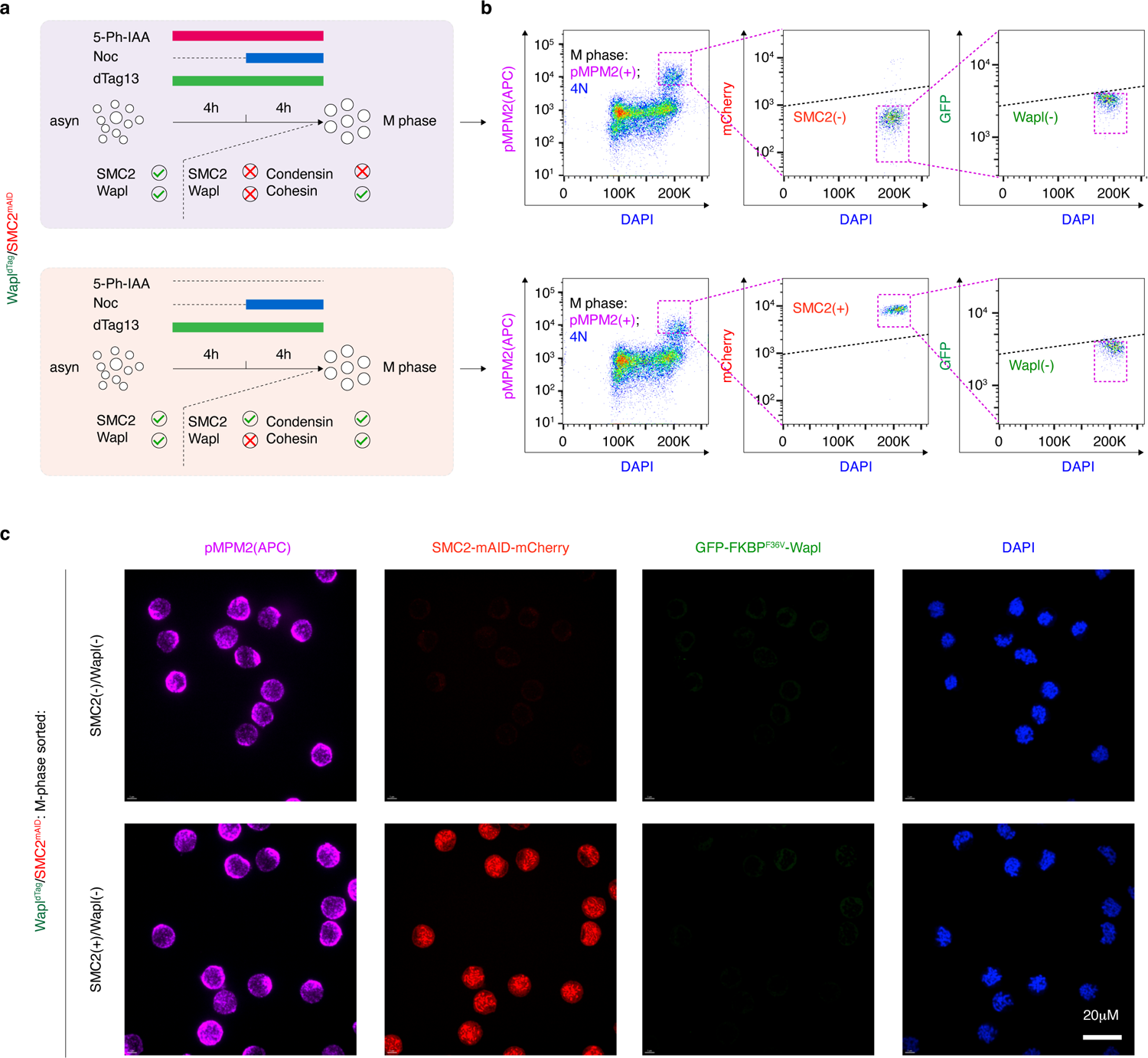
FACS purification of mitotic chromosomes. **a**, Experimental design, showing the strategy of prometaphase arrest in combination with 5-Ph-IAA or/and dTag13 treatment. **b**, Flow cytometry plot showing the gating strategy to purify Wapl-depleted and Wapl and SMC2 co-depleted mitotic cells. **c**, Representative images showing FACS purified Wapl-depleted or Wapl and SMC2 co-depleted mitotic cells. Two independent experiments were performed. Scale bar: 20μm.

**Extended Data Figure 3.**
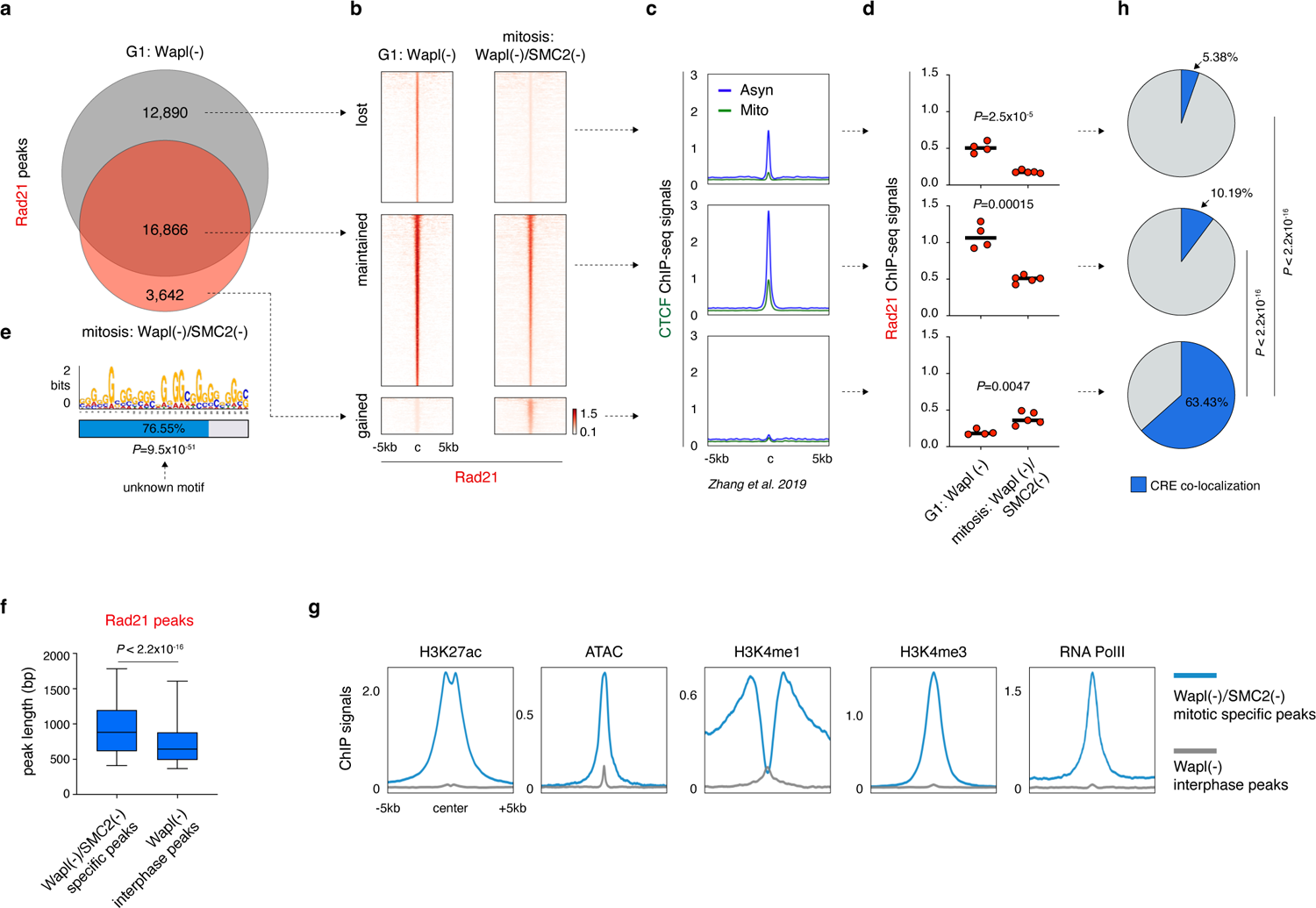
Cohesin forms focalized peaks in condensin-deficient mitotic chromosomes. **a**, Venn diagram showing the intersection results of Rad21 peaks in the Wapl deficient G1-phase samples and Wapl and SMC2 co-depleted mitotic samples. **b**, Density heatmap plots showing the ChIP-seq signals of Rad21 peaks that were lost, maintained or gained during mitosis. **c**, Meta region plots showing the corresponding mitotic CTCF ChIP-seq signals for Rad21 peaks that were lost, maintained or gained. Note that compared to the lost ones, the maintained Rad21 peaks were co-localized by stronger mitotic CTCF binding. CTCF ChIP-seq data was adapted from our previous study ^20^. **d**, Dot plots showing the average Rad21 ChIP-seq signals for peaks that were lost, maintained or gained during mitosis (n=4 and 5 for G1-phase and mitotic samples respectively). *P* values were calculated using two-sided student’s *t*-test. **e**, *De-novo* motif enrichment analysis for the mitotic specific Rad21 peaks. **f**, Box plot showing that mitotic specific Rad21 peaks were significantly larger compared to the interphase Rad21 peaks. For both box plots, central lines denote medians; box limits denote 25th–75th percentile; whiskers denote 5th–95th percentile. *P* values were calculated using a two-sided Wilcoxon signed-rank test. **g**, Meta region plots showing that mitotic specific Rad21 peaks were heavily decorated with active histone marks. **h**, Pie charts showing that the mitotic specific Rad21 peaks were significantly more likely to co-localize with CREs.

**Extended Data Figure 4.**
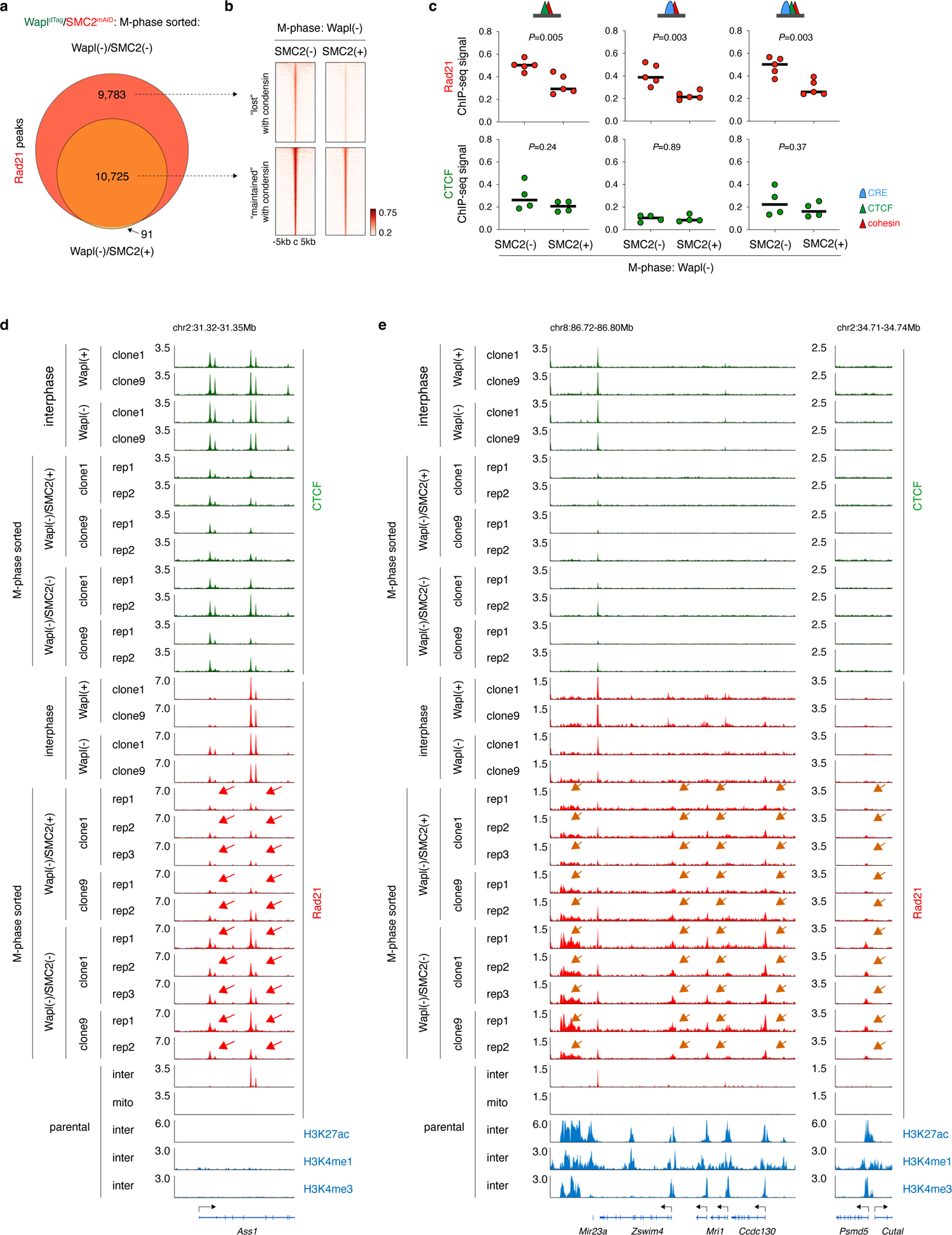
Condensin relocates mitotically retained Rad21 peaks. **a**, Venn diagram showing the intersection results of Rad21 peaks identified in Wapl-deficient mitotic cells with or without condensin. **b**, Density heatmap plots of Rad21 ChIP-seq signals in Wapl-deficient mitotic cells with or without condensin. Peaks that were lost or maintained upon the introduction of condensin were shown. **c**, Upper panel: Dot plots showing the significant reduction of mitotic Rad21 ChIP-seq signals in the presence of condensin. Rad21 peaks co-localized to CTCF, CRE or both were separately shown. *P* values were computed using two-sided student’s *t*-test (n=5 biological replicates). Lower panel: similar to the upper panel showing that the mitotic CTCF ChIP-seq signals were not significantly perturbed by condensin. **d**, Browser tracks of independent biological replicates showing the reduced Rad21 peaks (co-localized to CTCF, indicated by Red arrows) in response to condensin loading during mitosis. Tracks of interphase samples were shown as control. Tracks of mitotic and interphase CTCF ChIP-seq as well as interphase H3K27ac ChIP-seq results were also shown. **e**, Similar to (**d**), browser tracks showing of independent biological replicates showing the reduced Rad21 peaks (co-localized to CREs, indicated by orange arrows) in response to condensin loading during mitosis.

**Extended Data Figure 5.**
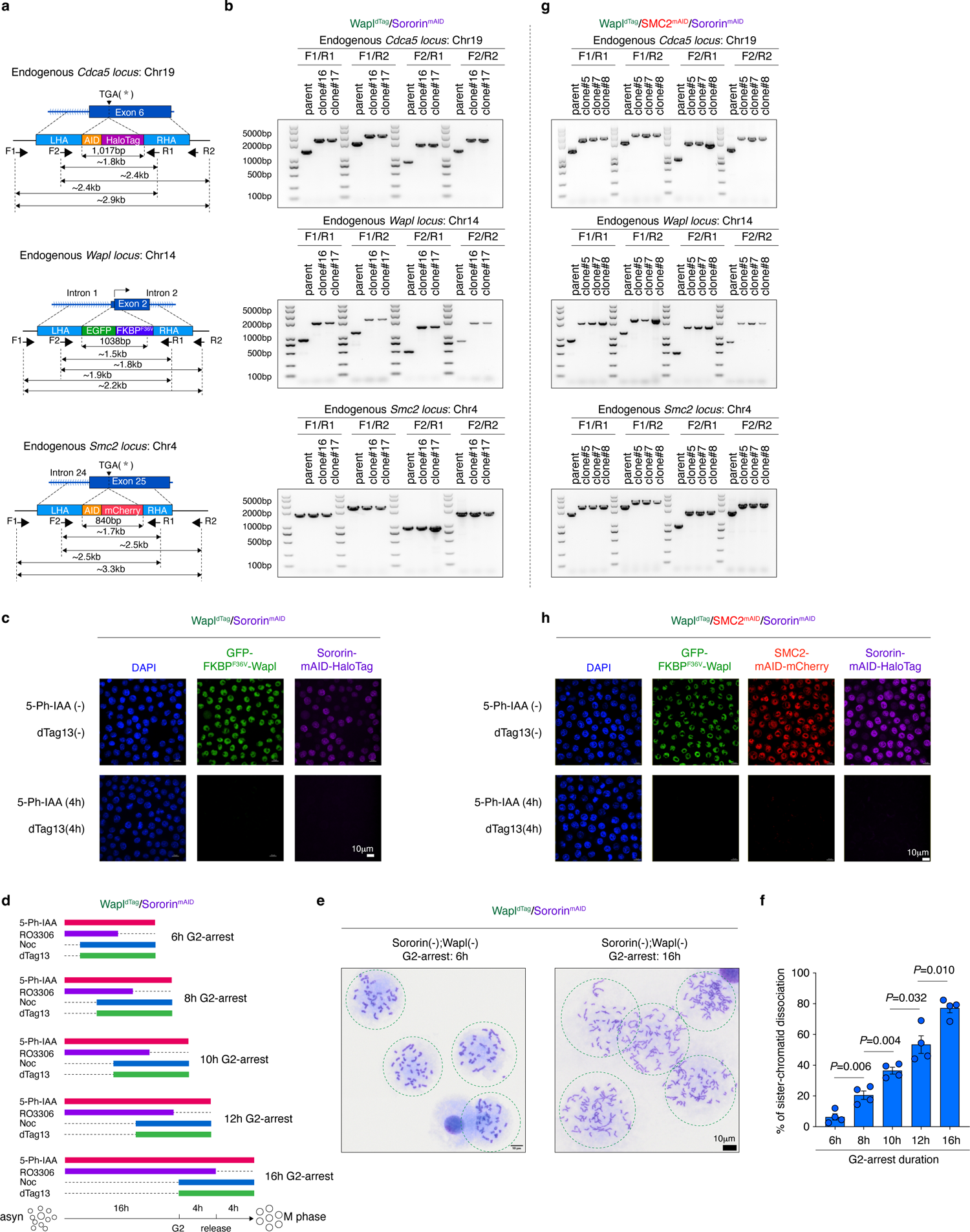
Characterization of the Wapl^dTag^/ Sororin^mAID^ and Wapl^dTag^/SMC2^mAID^/Sororin^mAID^ and cell lines. **a**, Schematic showing the edited *Cdca5*(encoding Sororin), *Wapl* and *Smc2* loci and the respective genotyping strategy. **b**, Genotyping results of the Wapl^dTag^/Sororin^mAID^ clones. **c**, Fluorescence images showing loss of Wapl and Sororin fluorescence signals upon 4 hours of dTag13 and 5-Ph-IAA treatment in the Wapl^dTag^/ Sororin^mAID^ cells. **d**, Schematic showing the strategy to determine G2/M arrest length. **e**, Metaphase spread analysis showing a more extensive dissociation of sister-chromatids in the Wapl^dTag^/ Sororin^mAID^ cells after 16 hours compared to 6 hours of RO-3306 mediated G2 arrest. **f**, Quantification of (**d** & **e**), showing progressive dissociation of sister-chromatids as G2-arrest duration were extended. Longer G2-arrest allowed more complete removal of cohesive-cohesin by Wapl. *P* values were calculated by two-sided student’s *t*-test. **g**, Genotyping results of Wapl^dTag^/SMC2^mAID^/Sororin^mAID^ clones. **h**, Fluorescence images showing loss of Wapl, Sororin and SMC2 fluorescence signals upon 4 hours of dTag13 and 5-Ph-IAA treatment in the Wapl^dTag^/SMC2^mAID^/Sororin^mAID^ cells.

**Extended Data Figure 6.**
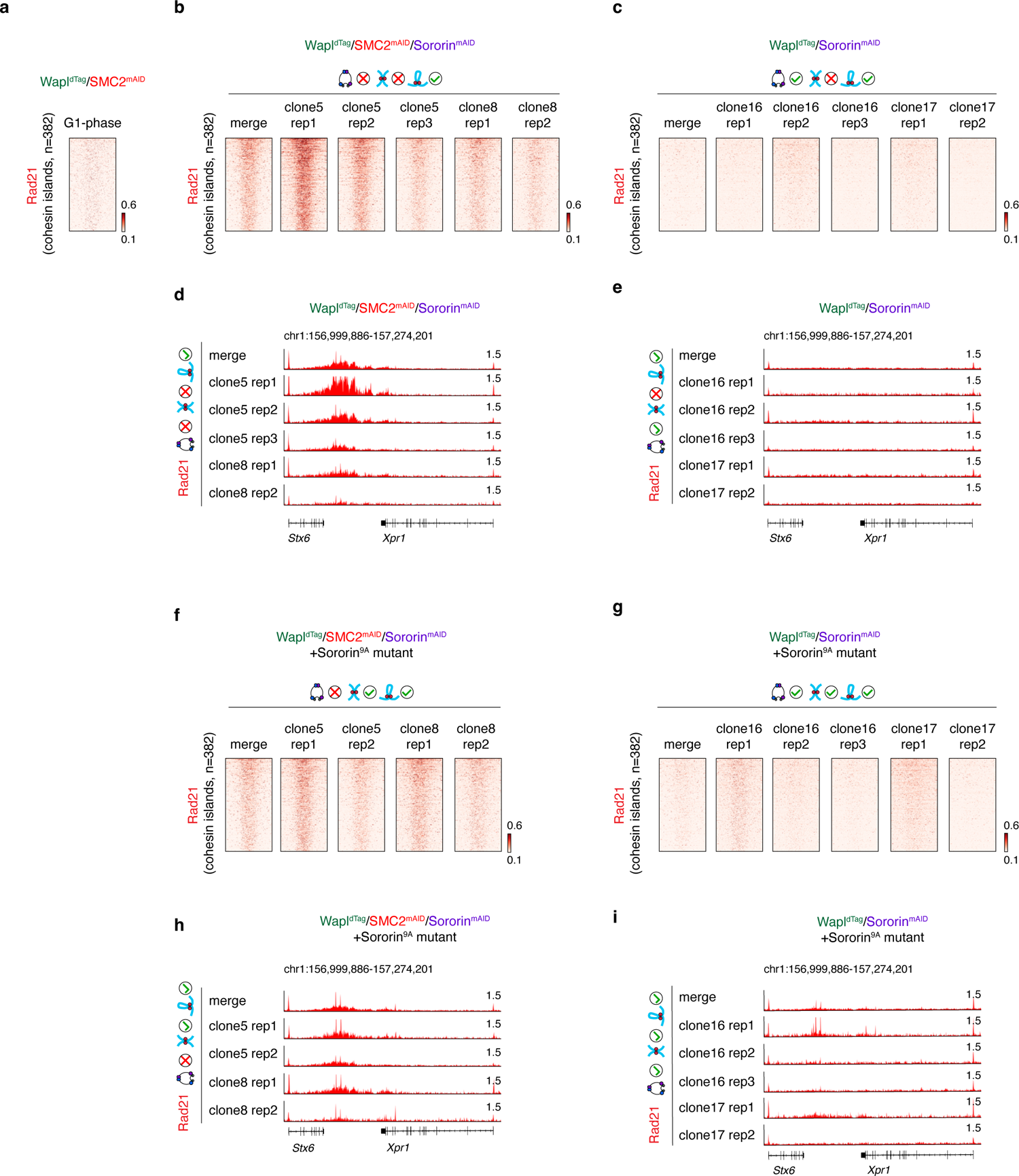
Cohesin islands are disrupted by condensin during mitosis. **a**, Density heatmap plots showing that cohesin islands were undetectable in G1 phase cells. **b**, Density heatmap plots of independent biological replicates showing the cohesin island signals by extrusive-cohesin during mitosis. **c**, Density heatmap plots showing that extrusive-cohesin mediated cohesin island signals were dramatically disrupted by condensin during mitosis. **d**, Browser tracks of independent biological replicates showing an example of cohesin island by extrusive-cohesin during mitosis. **e**, Browser tracks showing that the cohesin island in (**d**) was diminished by condensin during mitosis. **f** & **g**, Similar to (**b** & **c**) showing cohesin-islands in mitotic cells co-loaded with extrusive- and cohesive-cohesin. Cohesin islands were diminished by condensin during mitosis. **h** & **i**, Similar to (**d** & **e**), showing browser tracks of an exemplary cohesin island in independent mitotic biological replicates co-loaded with extrusive- and cohesive-cohesin. The cohesin island was disrupted by condensin during mitosis.

**Extended Data Figure 7.**
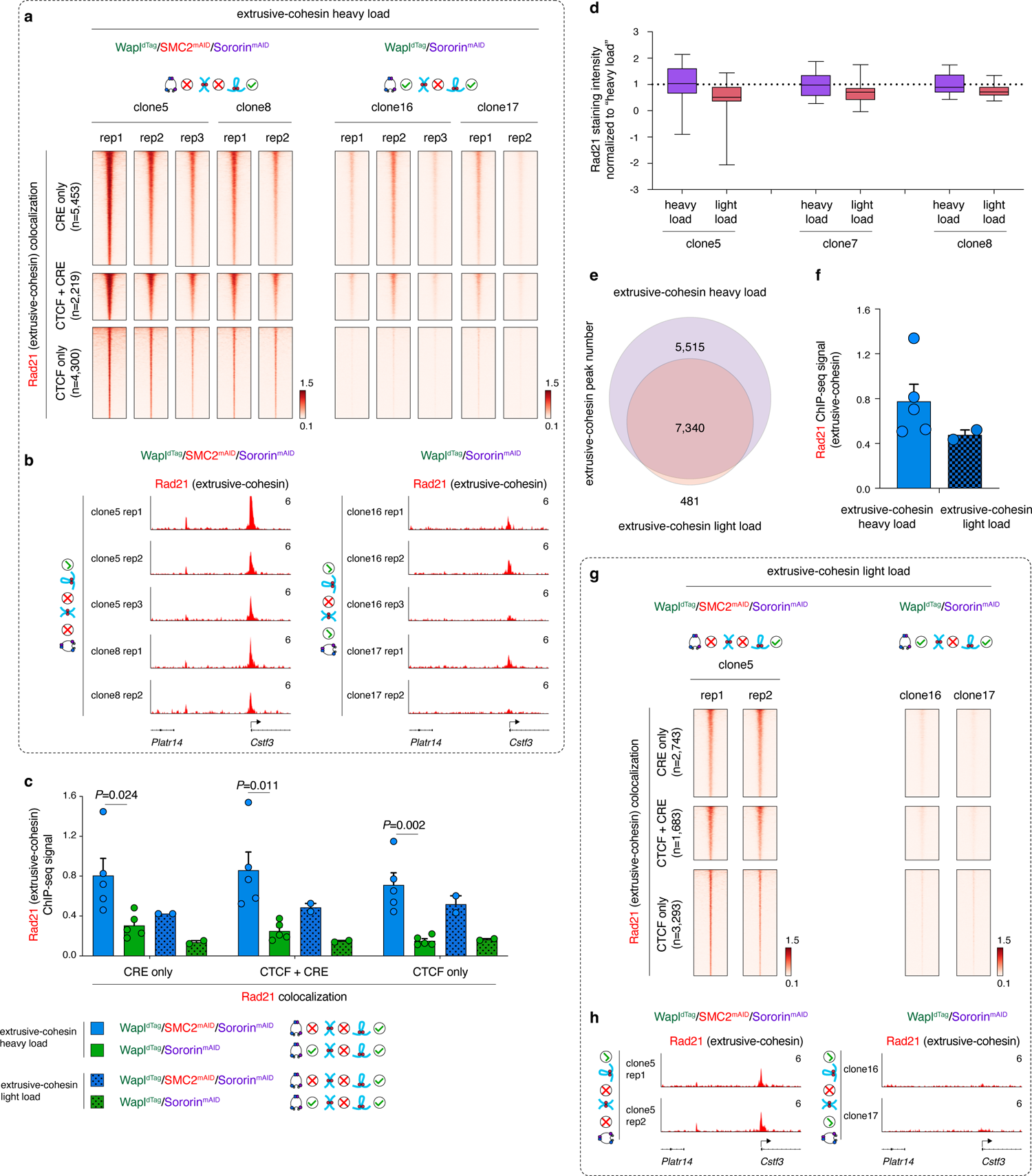
Extrusive-cohesin is relocated by condensin during mitosis. **a**, Density heatmap plots for independent biological replicates showing reduced extrusive-cohesin (“heavy load”) ChIP-seq signal intensity in response to condensin during mitosis. Extrusive-cohesin with different genomic contexts (co-localized to CTCF, CRE or both) were shown separately. **b**, Browser tracks of independent biological replicates showing weaker extrusive-cohesin (“heavy load”) peaks upon condensin loading. **c**, Bar graphs showing the quantification of (**a** and **g**), indicating reduced extrusive-cohesin (“heavy” or “light load”) ChIP-seq signals in the presence of condensin. *P* values were calculated by two-sided student’s *t*-test. **d**, Box plots showing reduced immunofluorescence staining signal of Rad21 in condensin-deficient mitotic samples lightly loaded with extrusive-cohesin, compared to those heavily loaded with extrusive-cohesin. Comparisons for independent clones were shown. For all box plots, central lines denote medians; box limits denote 25th–75th percentile; whiskers denote 5th–95th percentile. **e**, Venn-diagram showing the intersection results of Rad21 peaks identified in condensin-deficient mitotic samples heavily or lightly loaded with extrusive-cohesin. Note that cohesive-cohesin was not loaded in these samples. **f**, Dot plots showing the Rad21 ChIP-seq signal intensity in condensin-deficient mitotic cells heavily (n=4) or lightly (n=2) loaded with extrusive-cohesin. **g**, Density heatmap plots of independent biological replicates showing a complete loss of extrusive-cohesin (“light load”) ChIP-seq signals upon condensin loading. **h**, Browser tracks of independent biological replicates showing a complete loss of extrusive-cohesin (“light load”) ChIP-seq signals upon condensin loading.

**Extended Data Figure 8.**
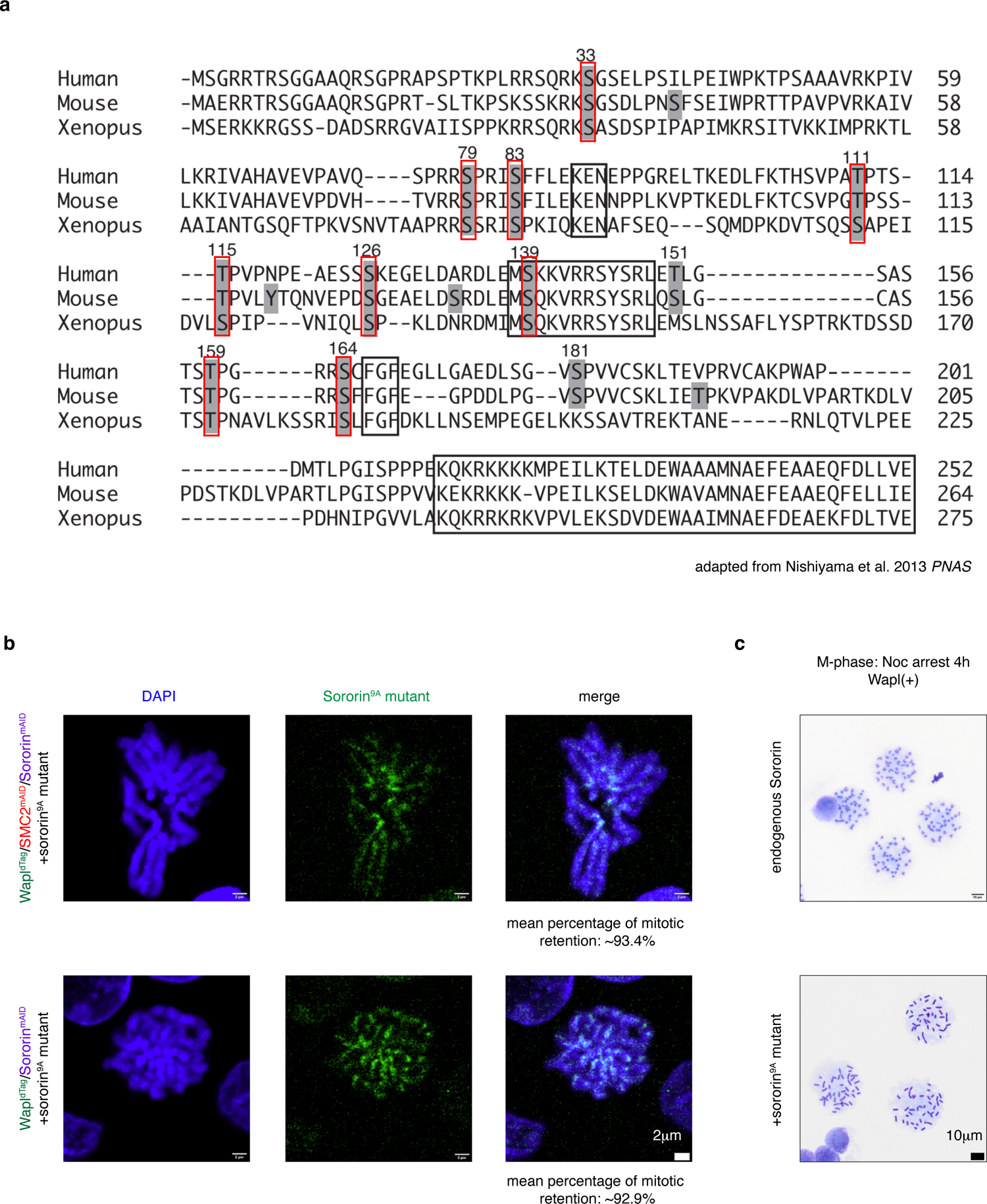
Characterization of Sororin^9A^ mutant. **a**, Schematic showing the evolutionarily conserved phosphosites in Sororin. Amino acids labeled by red boxes were mutated to alanine to generate Sororin^9A^ mutant. Image was adapted from the Nishiyama et al. 2013 PNAS paper ^35^. **b**, Representative images showing the mitotic retention of Sororin^9A^ mutant. **c**, Metaphase spread analysis of nocodazole arrested Wapl^dTag^/Sororin^mAID^ cells expressing Sororin^9A^ mutant, showing extensive cohesion between sister-chromatids during mitosis even in the presence of Wapl.

**Extended Data Figure 9.**
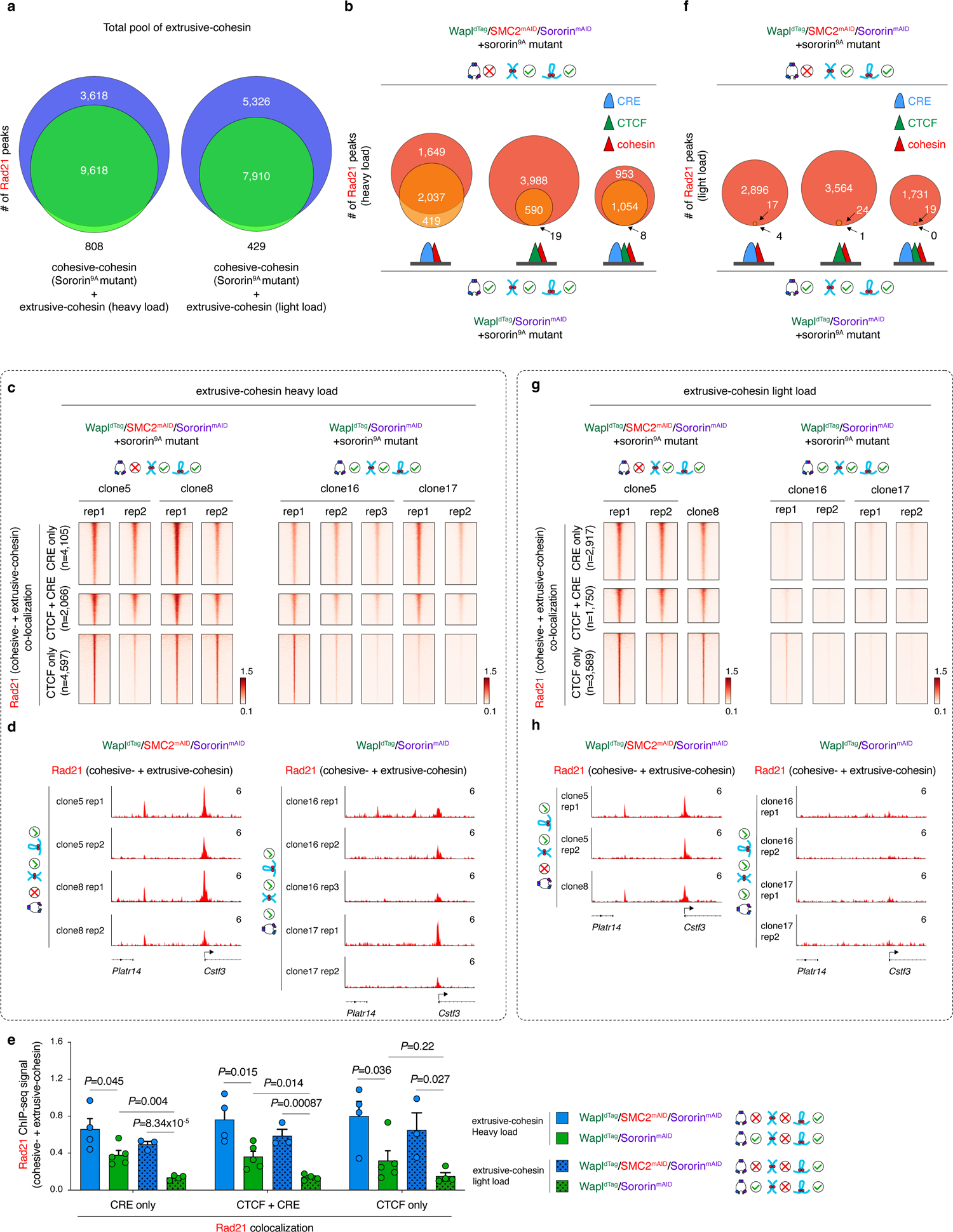
Condensin disrupts extrusive- and cohesive-cohesin peaks when they are co-loaded. **a**, Venn-diagrams showing the extensive overlap of Rad21 peak calls for extrusive-cohesin alone and cohesive-plus extrusive-cohesin in both “heavy” and “light load” conditions. **b**, Venn-diagrams showing loss of extrusive-(“heavy load”) + cohesive-cohesin peaks during mitosis upon condensin loading. Peaks co-localized to CTCF, CREs or both were shown separately. **c**, Density heatmap plots of independent biological replicates showing reduced Rad21 ChIP-seq signals (“heavy load” extrusive-cohesin + cohesive-cohesin) upon condensin loading during mitosis. Rad21 peaks with different genomic contexts (co-localized to CTCF, CRE or both) were shown separately. **d**, Browser tracks showing weakening of Rad21 peaks (“heavy load” extrusive-cohesin + cohesive-cohesin) in response to condensin. **e**, Bar graphs showing the quantification of (**c** and **g**), indicating reduced Rad21 (“heavy” or “light load” extrusive-cohesin + cohesive-cohesin) ChIP-seq signals in response to condensin during mitosis. *P* values were calculated by two-sided student’s *t*-test. **f**, Similar to (**b**), showing a complete loss of extrusive-(“light load”) + cohesive-cohesin peaks during mitosis upon condensin loading. **g**, Similar to (**c**). Density heatmap plots of independent biological replicates showing complete loss of Rad21 ChIP-seq signals (“light load” extrusive-cohesin + cohesive-cohesin) upon condensin loading during mitosis. **h**, Browser tracks of independent biological replicates showing a complete loss of Rad21 ChIP-seq signals (“light load” extrusive-cohesin + cohesive-cohesin) upon condensin loading.

**Extended Data Figure 10.**
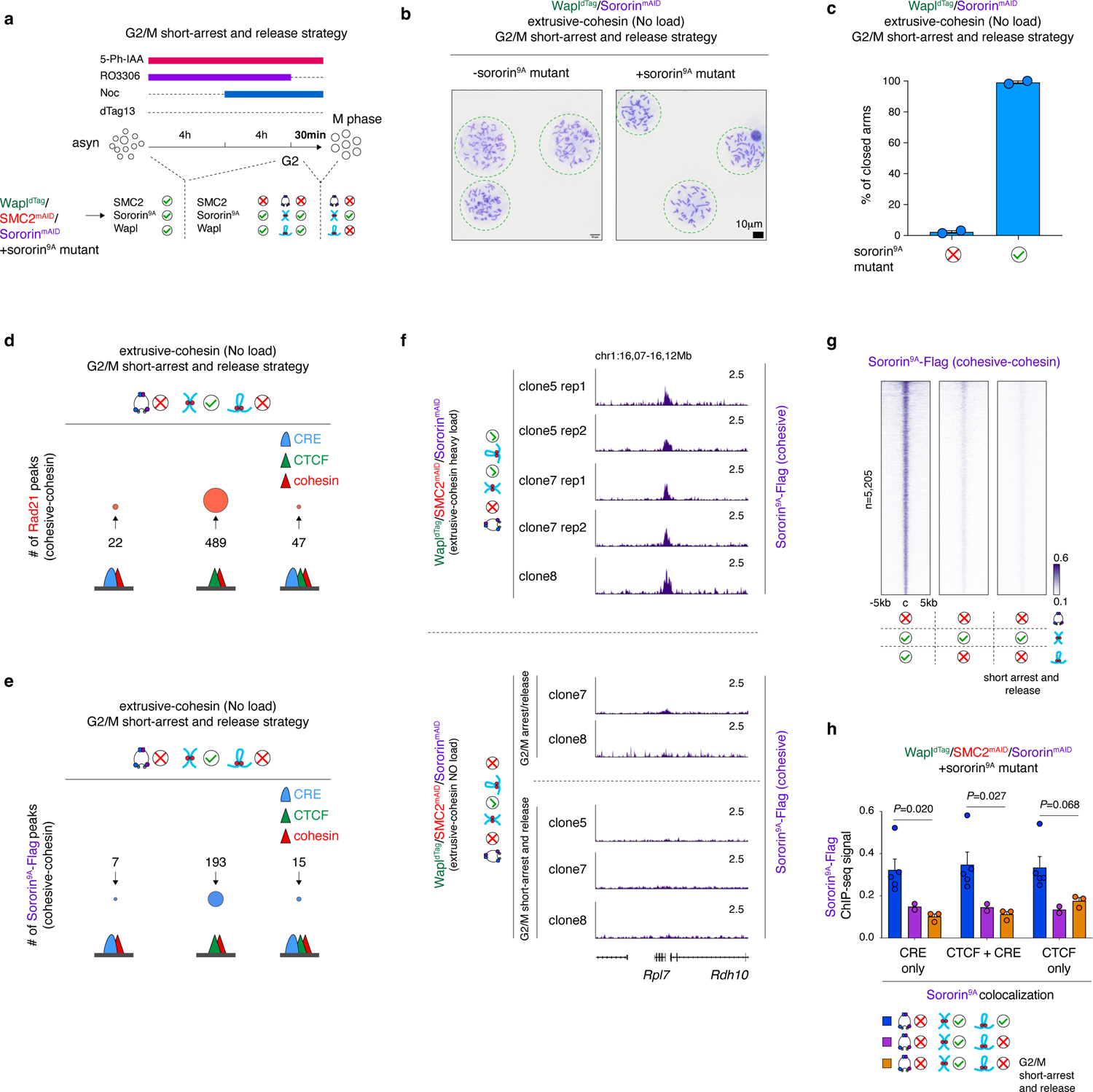
G2/M short-arrest and release protocol confirms that cohesive-cohesin alone does not form focalized peaks without extrusive-cohesin. **a**, Schematic showing the G2/M short-arrest and release protocol to obtain condensin-deficient mitotic chromosomes loaded with cohesive-cohesin alone. b, Metaphase spread analysis using the G2/M short-arrest and release protocol showing extensive sister-chromatid cohesion in the Wapl replete mitotic cells with Sororin^9A^ mutant. In comparison, cells without Sororin^9A^ mutant lost the cohesion between sister chromatids. c, Quantification of (b), showing the percentage of “closed arm” during mitosis for indicated conditions. d, Venn-diagrams showing very few Rad21 peak calls in condensin-deficient mitotic cells with cohesive-but not extrusive-cohesin (G2/M short-arrest and release protocol). e, Venn-diagrams showing very few Sororin^9A^ peak calls in condensin-deficient mitotic cells with cohesive-but not extrusive-cohesin (G2/M short-arrest and release protocol). f, Browser tracks showing loss of flag-tagged Sororin^9A^ (cohesive-cohesin) peaks in the condensin-deficient mitotic chromosomes without extrusive-cohesin. Samples obtained through original or short-arrest and released protocols were shown. Tracks for independent biological replicates were shown. g, Density heatmap plots of the flag-tagged Sororin^9A^, showing that cohesive-cohesin peaks were almost completely lost in the absence of extrusive-cohesin. Samples generated by both original or short-arrest and release protocols were shown. h, Bar graphs showing the dramatic loss of flag-tagged Sororin^9A^ ChIP-seq signals in the absence of extrusive-cohesin. *P* values were calculated using two-sided student’s *t*-test. Peaks colocalized to CTCF, CRE or both were shown separately.

**Extended Data Figure 11.**
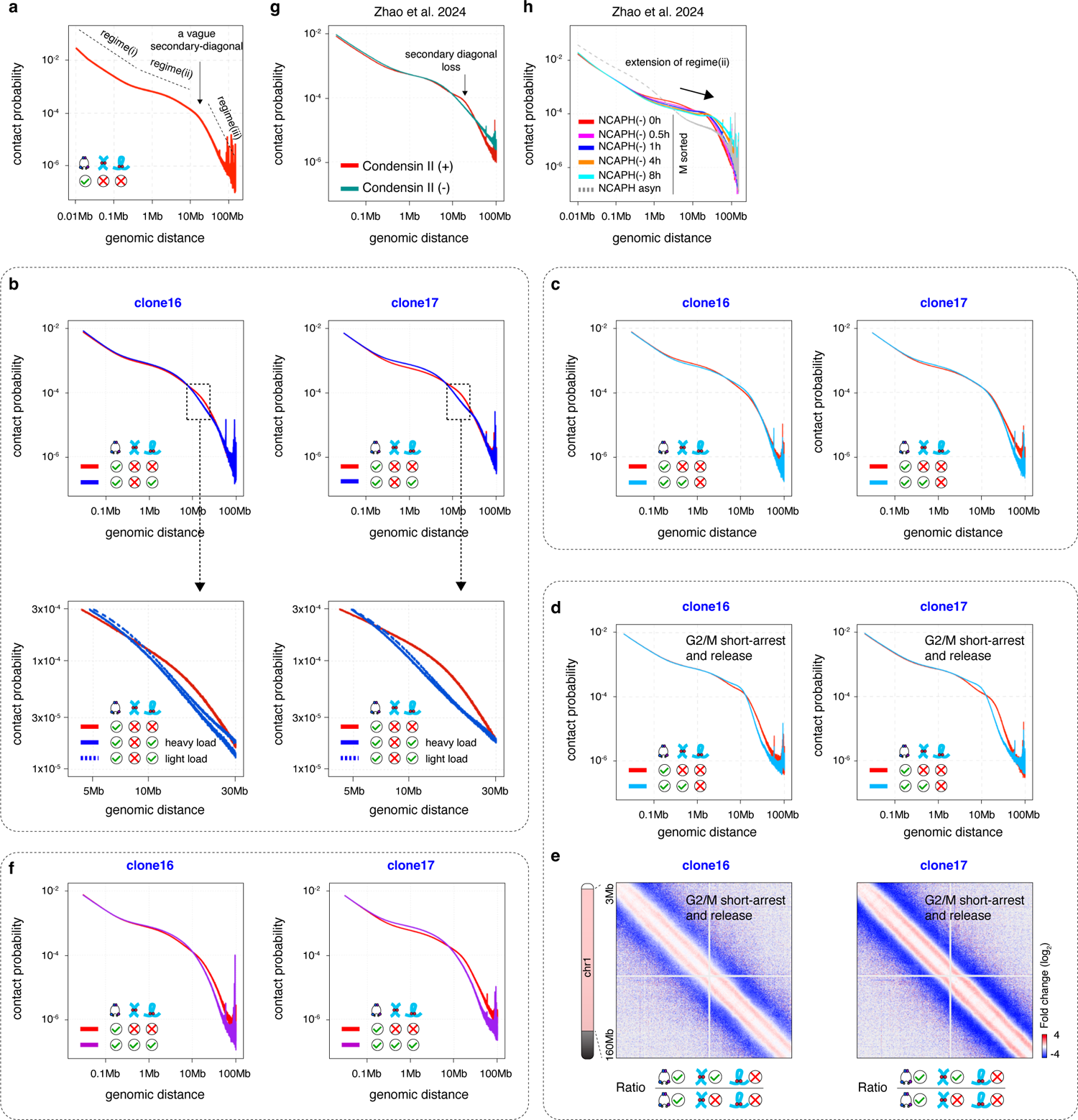
Influence of cohesive- or/and extrusive-cohesin on mitotic chromosome architecture. **a**, *P(s)* curve of the control mitotic cells. The three regimes were indicated by dotted lines. The secondary diagonal between regime (ii) and (iii) were indicated by arrow. **b**, Upper panel: *P(s)* curves of independent clones showing the influence of extrusive-cohesin (“heavy load”) on mitotic chromosome architecture. Lower panel: Zoom in *P(s)* curves showing the influence of heavily or lightly loaded extrusive-cohesin on mitotic chromosome architecture. **c**, *P(s)* curves of independent clones showing the influence of cohesive-cohesin on mitotic chromosome architecture. **d**, *P(s)* curves of independent clones showing the influence of cohesive-cohesin (G2/M short-arrest and release protocol) on mitotic chromosome architecture. **e**, KR-balanced Hi-C contact matrices for independent clones showing the log_2_ fold change of contact probabilities when cohesive-cohesin was loaded (G2/M short-arrest and release protocol). **f**, *P(s)* curves of independent clones showing the combined influence of cohesive-cohesin and extrusive-cohesin on mitotic chromosome architecture. **g**, *P(s)* curves showing the effect of condensin II depletion on mitotic chromosome architecture. Data adapted from our previous study ^24^. **h**, *P(s)* curves showing the effect of condensin I depletion on mitotic chromosome architecture. Data adapted from our previous study ^24^.

**Extended Data Figure 12.**
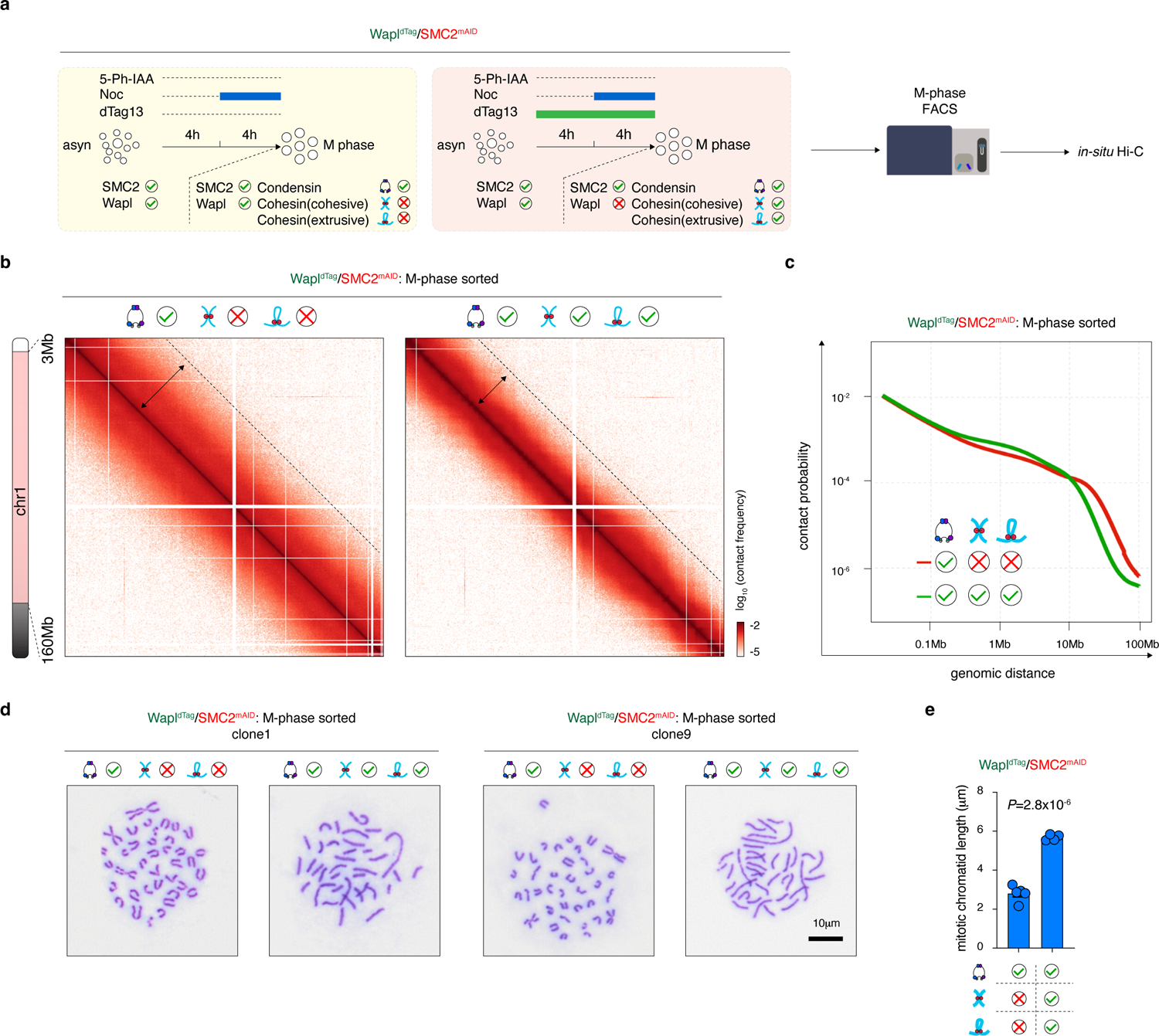
Co-loading of extrusive- and cohesive-cohesin inhibits mitotic chromosome condensation in the Wapl^dTag^/SMC2^mAID^ cells. **a**, Schematics showing the nocodazole based prometaphase arrest strategy in combination with dTag13 treatment to obtain condensin-replete mitotic chromosomes with or without cohesin. **b**, KR-balanced Hi-C contact matrices showing the condensin-replete mitotic cells with or without cohesin retention. Bin size: 100kb. **c**, *P(s)* curves showing the influence of extrusive-cohesin and cohesive-cohesin (mediated by wildtype Sororin) on mitotic chromosome architecture. **d**, Metaphase spread analysis showing that cohesin (extrusive- and cohesive-cohesin) induces longer mitotic chromosomes. **e**, Bar graphs showing the quantification of (**d**). *P* values were calculated suing student’s *t*-test.

**Extended Data Figure 13.**
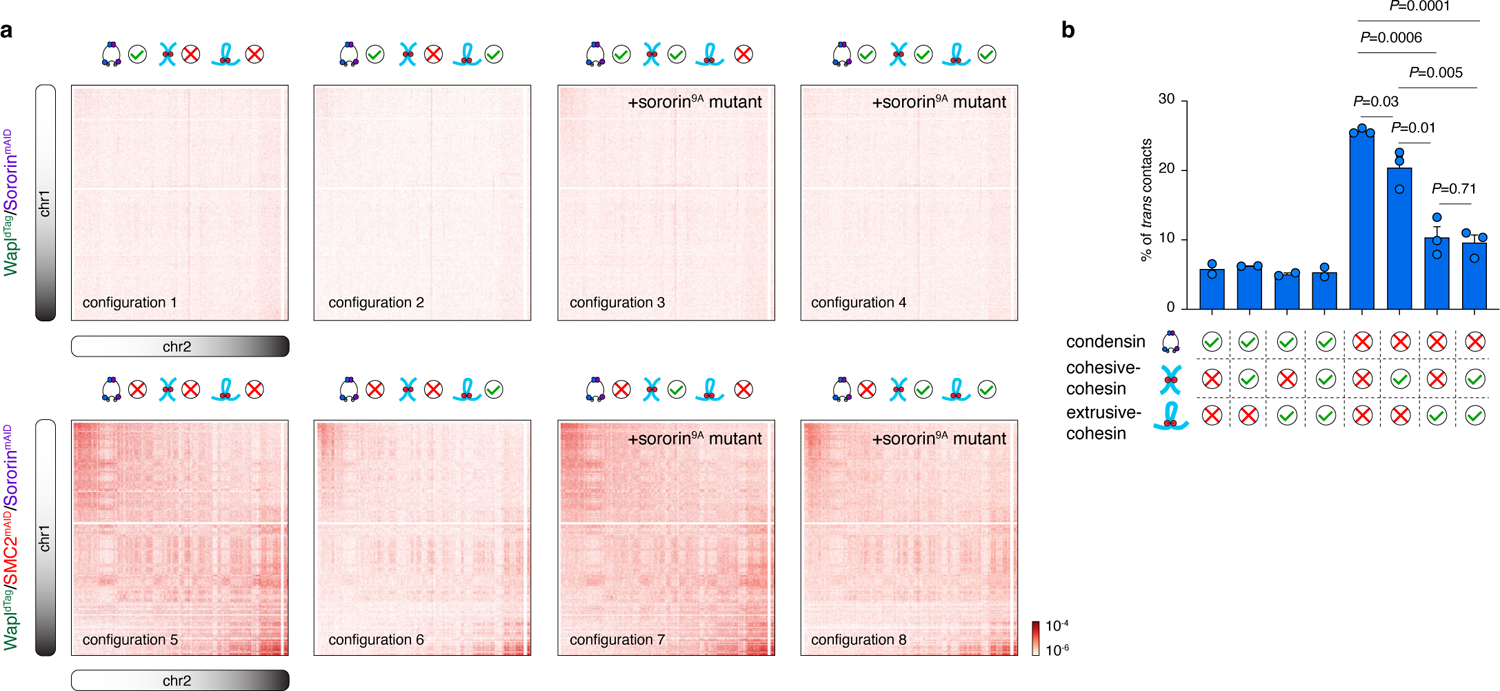
Extrusive-cohesin facilitates chromatin individualization. **a**, KR-balanced Hi-C contact matrices showing the *trans*-chromosomal interactions between chr1 and chr2 in mitotic cells with eight distinct SMC protein complex configurations. **b**, Bar graph showing the percentage of *trans* contacts in mitotic cells with eight distinct SMC protein complex configurations. *P* values were calculated using two-sided student’s *t*-test.

**Extended Data Figure 14.**
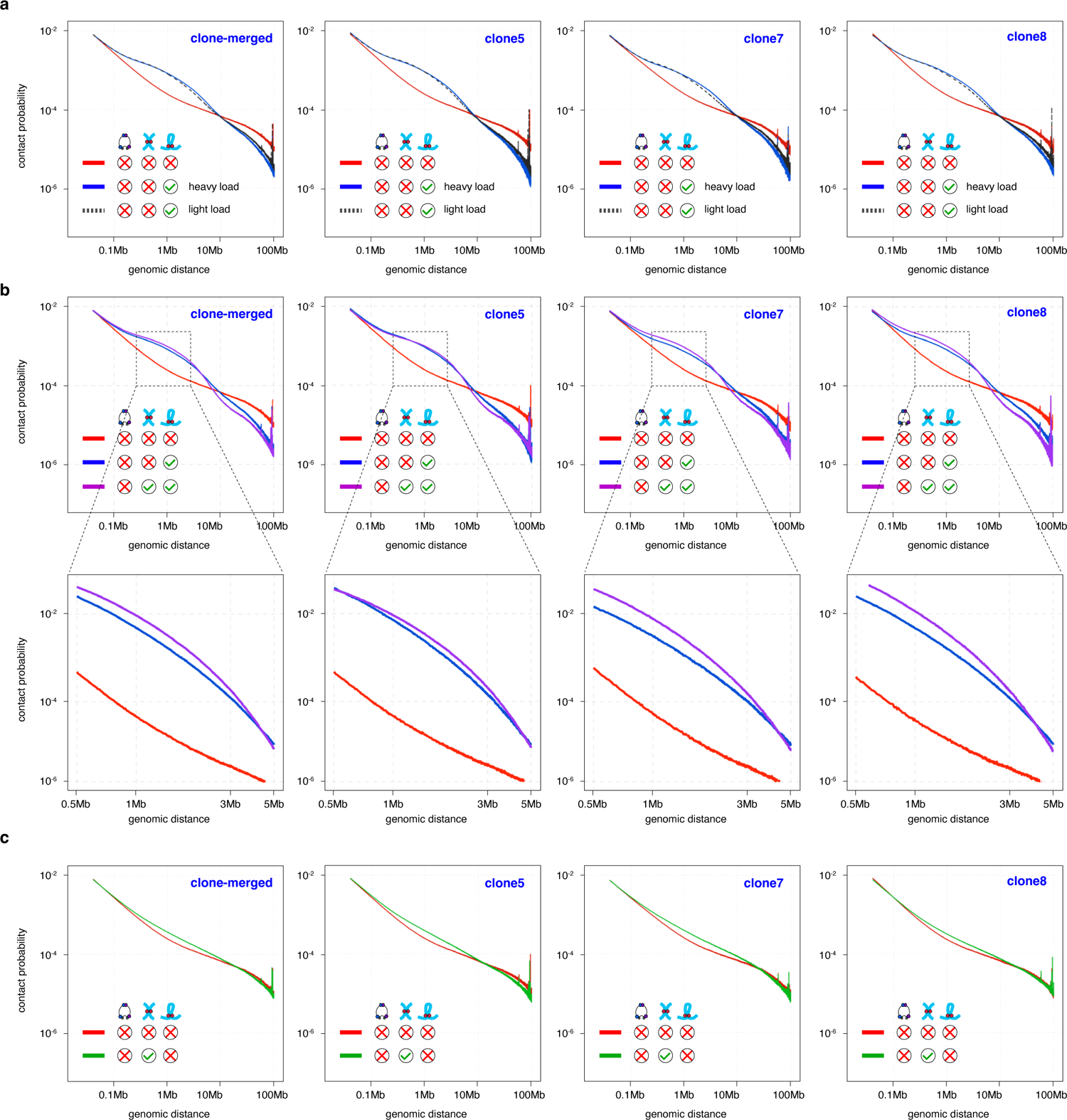
Extrusive-cohesin re-configures condensin-deficient mitotic chromosomes. **a**, *P(s)* curve showing gain of short-range contact frequencies in the condensin-deficient mitotic chromosomes by extrusive-cohesin. Note that light-loaded extrusive-cohesin displayed similar but milder effects. Curves for independent clone were shown. **b**, Upper panel: *P(s)* showing a further increase of short-range contact frequencies when cohesive-cohesin was co-loaded with extrusive-cohesin. Lower panel: zoom-in region of the upper panel. **c**, *P(s)* curve showing the influence of cohesive-cohesin alone on the condensin-deficient mitotic chromosomes.

**Extended Data Figure 15.**
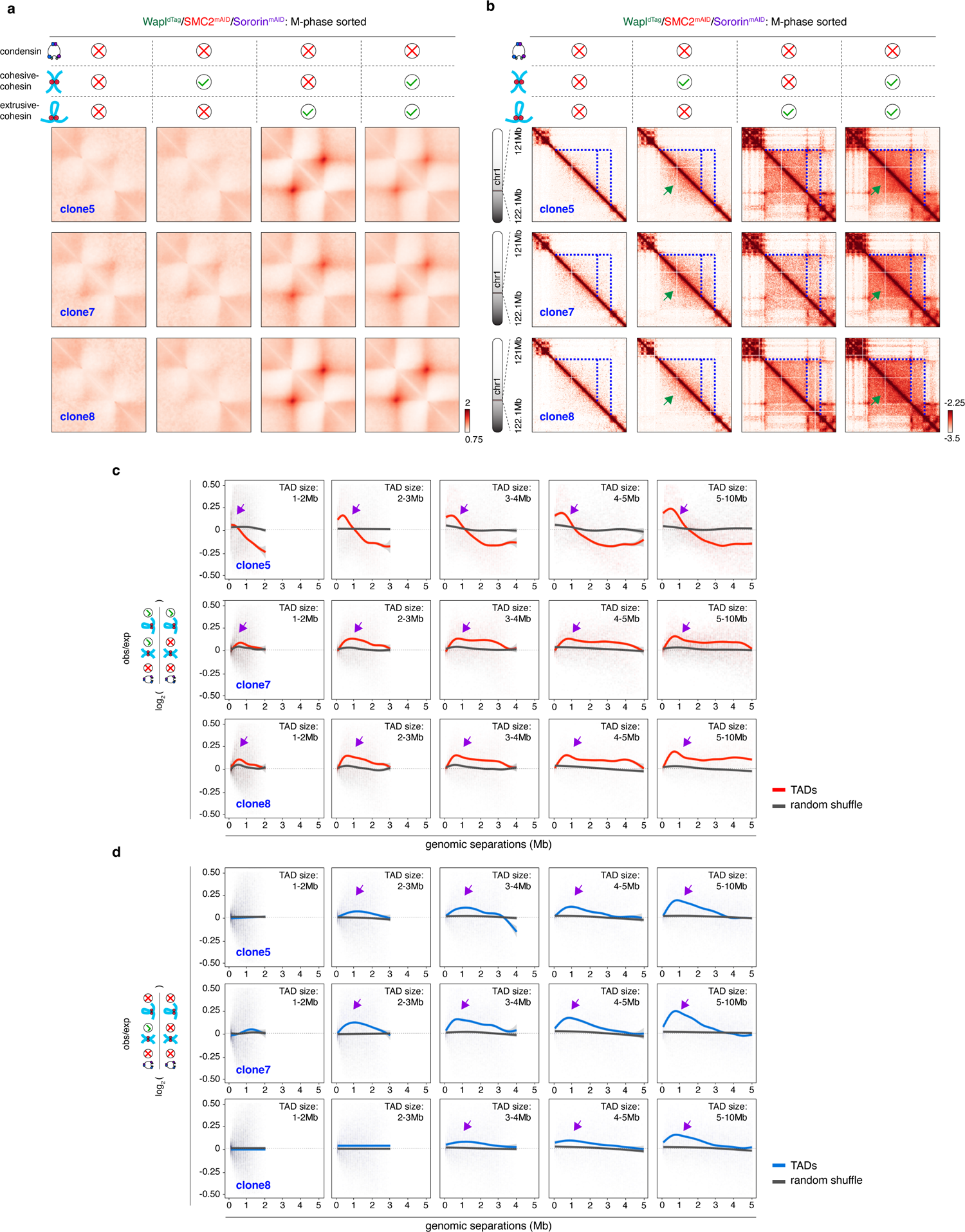
Additional data showing that cohesive-cohesin intensifies TADs. **a**, Composite domain plots for independent biological replicates showing that extrusive-cohesin mediates TADs with corner dot signals in the condensin-deficient mitotic chromosomes. **b**, KR-balanced Hi-C contact matrices showing the intensification (indicated by green arrows) of TADs (dotted blue lines) by cohesive-cohesin. Maps for independent biological replicates were shown. Bin size: 25kb. **c**, Line plots showing the gain of short-range contacts within TADs (indicated by purple arrows) in response to cohesive-cohesin in the condensin-depleted mitotic chromosomes. TADs were grouped based on their sizes. Plots for independent clones were shown. This observation was not seen in random position controls. **d**, Line plots showing the gain of short-range interactions within pre-defined TAD regions (purple arrows) upon cohesive-cohesin loading even without extrusive-cohesin. Plots for independent clones were shown. This observation was not seen in random position controls.

**Extended Data Figure 16.**
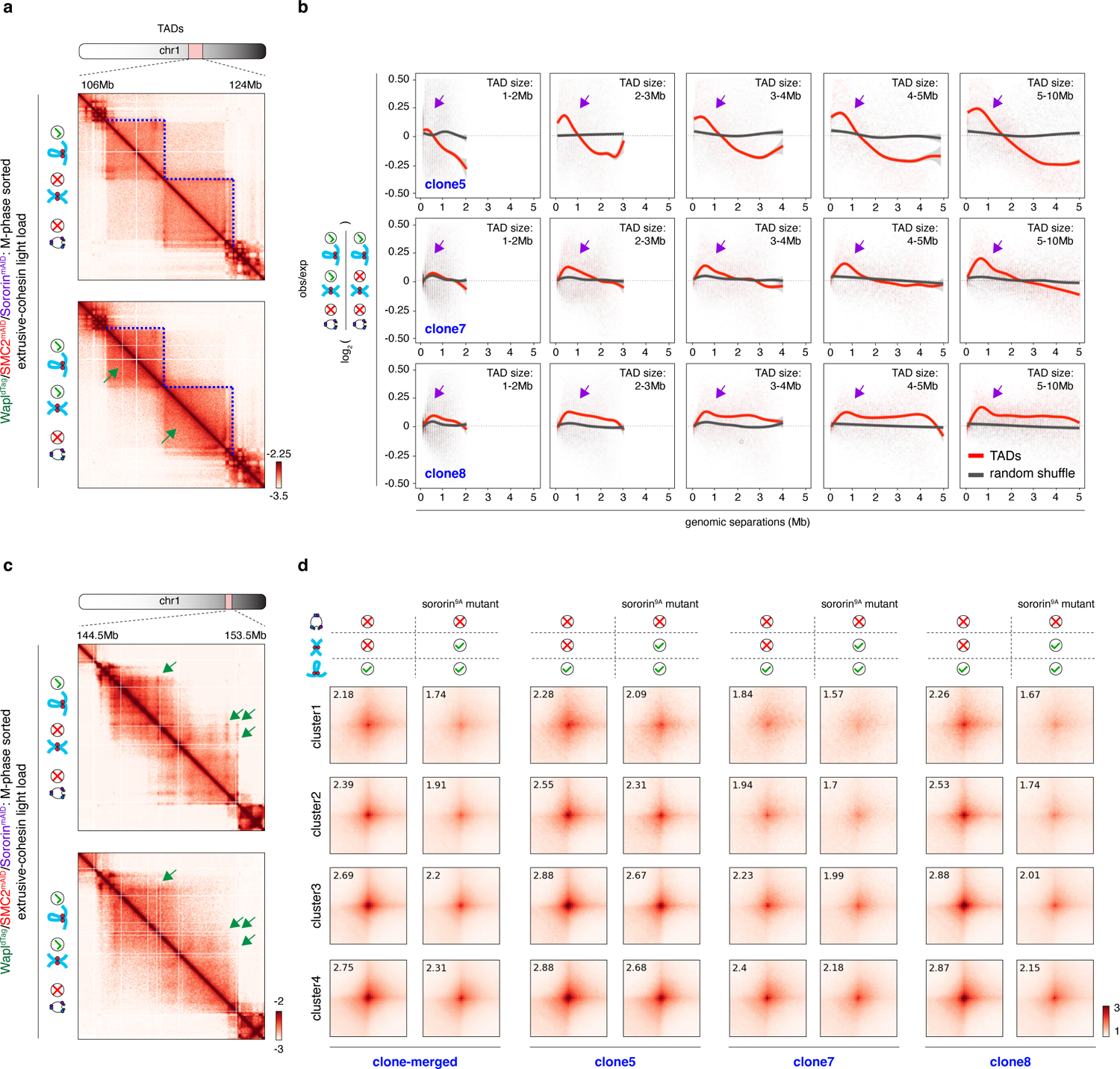
Cohesive-cohesin influences extrusive-cohesin (“light load”) mediated TADs and structural loops. **a**, KR-balanced Hi-C contact matrices showing the intensification (indicated by green arrows) of extrusive-cohesin (light load) mediated TADs (indicated by blue dot lines) upon the introduction of cohesive-cohesin. Bin size: 25kb. **b**, Line plots showing the gain of short-range contacts within extrusive-cohesin (“light load”) mediated TADs (indicated by purple arrows) in response to cohesive-cohesin in the condensin-depleted mitotic chromosomes. TADs were grouped based on their sizes. Plots for independent clones were shown. This observation was not seen in random position controls. **c**, KR-balanced Hi-C contact matrices showing the weakening (indicated by green arrows) of extrusive-cohesin (“light load”) mediated structural loops by cohesive-cohesin. Bin size: 25kb. **d**, APA plots showing the cohesive-cohesin mediated reduction of peak signals for structural loops from all four clusters. Plots for independent clones were shown.

**Extended Data Figure 17.**
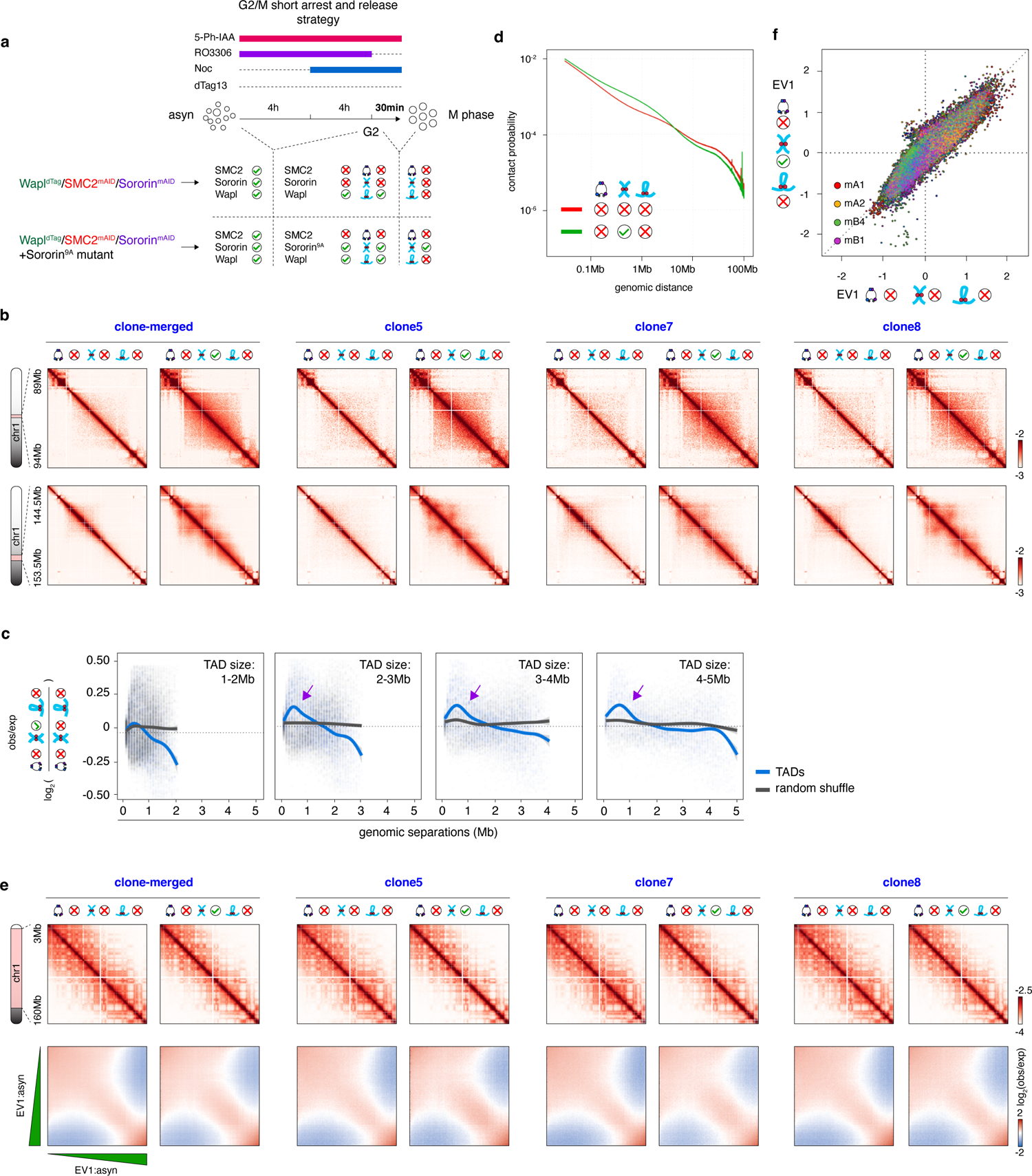
Influence of cohesive-cohesin alone (short-arrest and release protocol) on the condensin-deficient mitotic chromosomes. **a**, Schematic showing the G2/M short-arrest and release protocol to obtain condensin-deficient mitotic cells loaded with cohesive-cohesin alone. Condensin-deficient mitotic cells without cohesive-cohesin were obtained using the same protocol as control. **b**, KR-balanced Hi-C contact matrices showing two representative examples of cohesive-cohesin mediated intensification of pre-defined TAD regions in the absence of extrusive-cohesin. Plots for independent clones were shown. Bin size: 25kb. **c**, Line plots showing the gain of short-range interactions within pre-defined TAD regions (purple arrows) upon cohesive-cohesin loading (short-arrest and release protocol) without extrusive-cohesin. This observation was not seen in random position controls. **d**, *P(s)* curves showing cohesive-cohesin (short-arrest and release protocol) mediated slight gain of short-range interactions in the condensin-deficient mitotic chromosomes without extrusive-cohesin. **e**, Upper panel: KR-balanced Hi-C contact matrices showing the cohesive-cohesin (short-arrest and release protocol) alone does not modulate mitotic compartmentalization patterns. Maps for independent clones were shown. Bin size: 100kb. Lower panel: Saddle plots showing that cohesive-cohesin (short-arrest and release protocol) alone does not influence mitotic compartmentalization strength. **f**, Scatter plots showing the high correlation of EV1 values between the condensin-deficient mitotic cells without cohesive-cohesin (*x*-axis) and with cohesive-cohesin alone (short-arrest and release protocol) (*y*-axis).

**Extended Data Figure 18.**
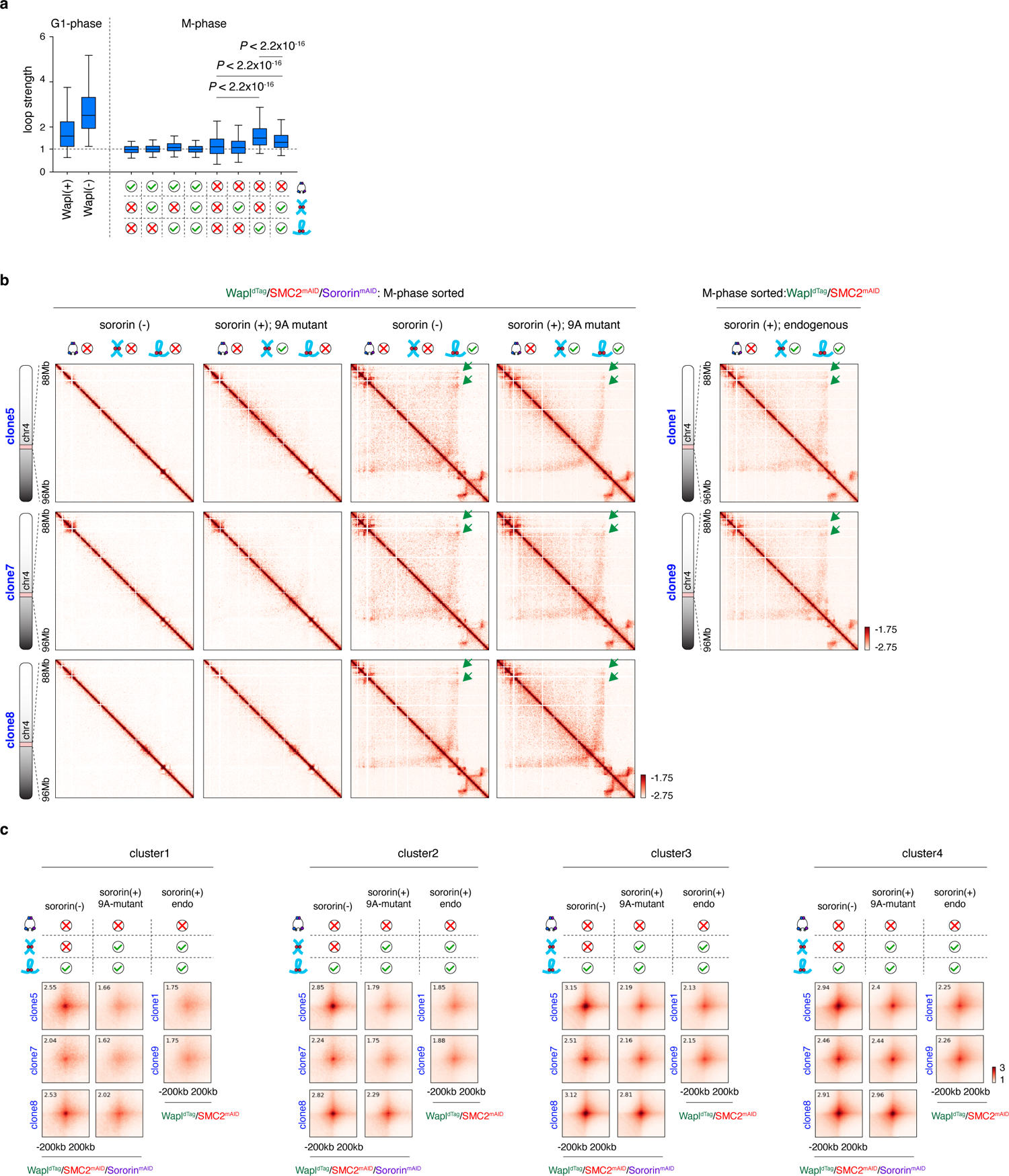
Additional data showing that cohesive-cohesin attenuates extrusive-cohesin mediated structural loops. **a**, Box plots showing the structural loop strength in G1-phase and indicated mitotic samples. Note that structural loops were stronger in condensin-deficient mitotic samples loaded with extrusive-cohesin alone compared to any other mitotic samples. For all box plots, central lines denote medians; box limits denote 25th–75th percentile; whiskers denote 5th–95th percentile. *P* values were calculated using a two-sided Wilcoxon signed-rank test. **b**, Left panel: KR-balanced Hi-C contact matrices showing reduced structural loops (green arrows) upon cohesive-cohesin (mediated by Sororin^9A^) loading. Right panel: KR-balanced Hi-C contact matrices of the same genomic locus as left panel, showing reduced structural loops signal intensity (green arrows) upon cohesive-cohesin (mediated by endogenous Sororin) loading. Plots of independent clones were shown. Bin size: 25kb. **c**, APA plots showing reduction of mitotic structural loop signals upon the loading of cohesive-cohesin (mediated by either Sororin^9A^ or endogenous wildtype Sororin). Structural loops from all four different clusters were compiled separately.

**Extended Data Figure 19.**
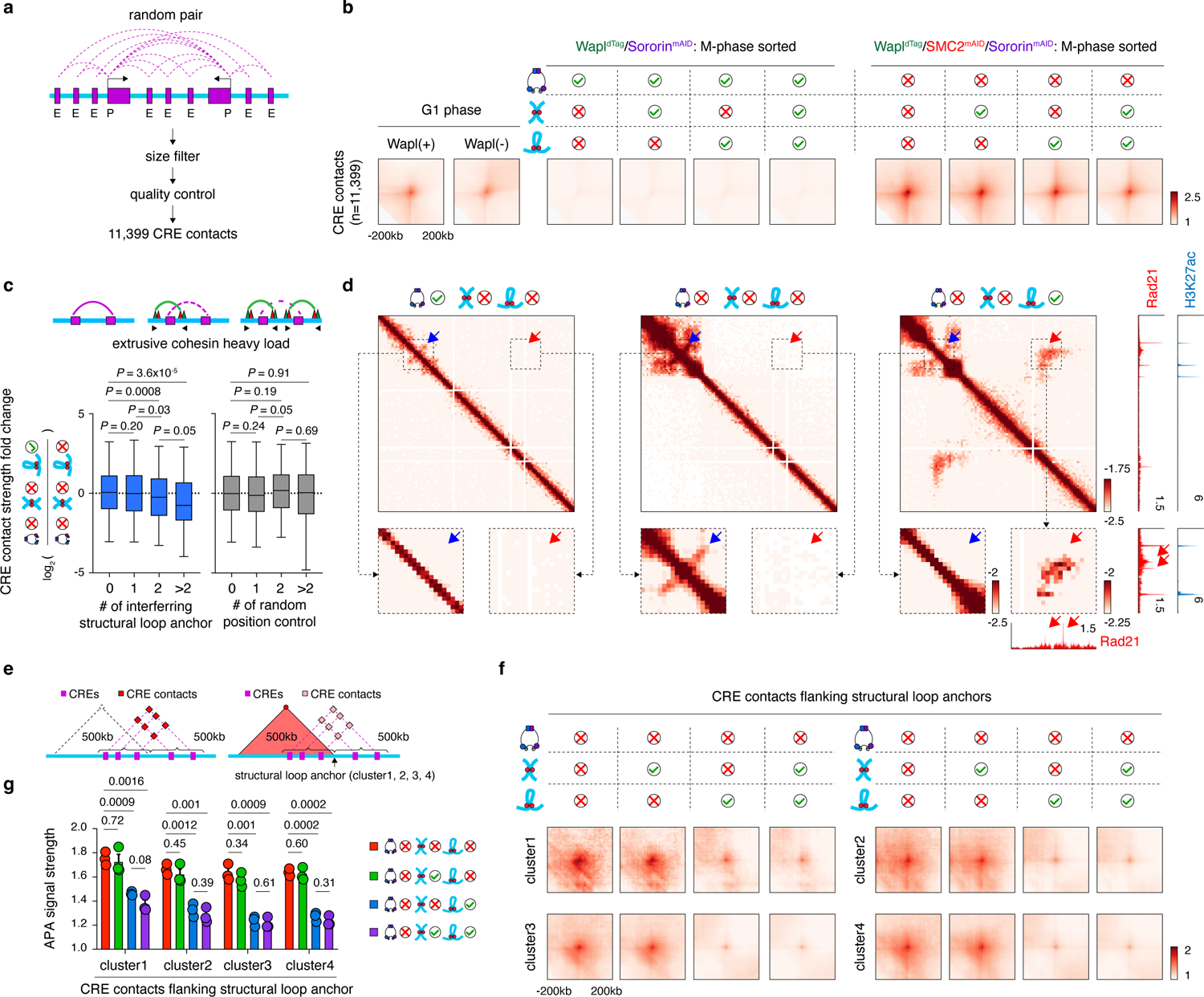
Cohesive-cohesin does not influence the insulation function of structural loops. **a**, Schematic showing stratification of the 11,399 CRE (purple bars) contacts. **b**, APA plot showing the composite signals of CRE contacts in interphase and mitotic samples. **c**, Box plots showing the log_2_ fold change of CRE contacts encompassing 0, 1, 2 or >2 structural loop anchors. For all box plots, central lines denote medians; box limits denote 25th–75th percentile; whiskers denote 5th–95th percentile. *P* values were calculated using a two-sided Wilcoxon signed-rank test. Randomized position controls were shown in parallel. **d**, KR-balanced Hi-C contact matrices showing reduced CRE contact strength (blue arrow) when interrupted by a nearby structural loop (red arrow). Bin size: 25kb. Condensin-replete mitotic cells were shown as control. Browser tracks of corresponding mitotic extrusive-cohesin peaks as well as interphase H3K27ac peaks were shown. Extrusive-cohesin peaks that mediate structural loop formation were also marked by red arrows. **e**, Schematic showing the pair strategy of CREs within a 500kb-range flanking a structural loop anchor. Anchors belonging to cluster1, 2, 3 and 4 structural loops were processed independently. **f**, APA plots showing the composite signals of CRE contact grouped by (**e**) in all four condensin-deficient mitotic samples with different cohesin configurations. Note that addition of cohesive-cohesin did not restore extrusive-cohesin insulated CRE contacts. **g**, Bar graphs showing the quantification of (**f**). n=3 independent biological replicates. *P* values were calculated using two-sided student’s *t*-test.

**Extended Data Figure 20.**
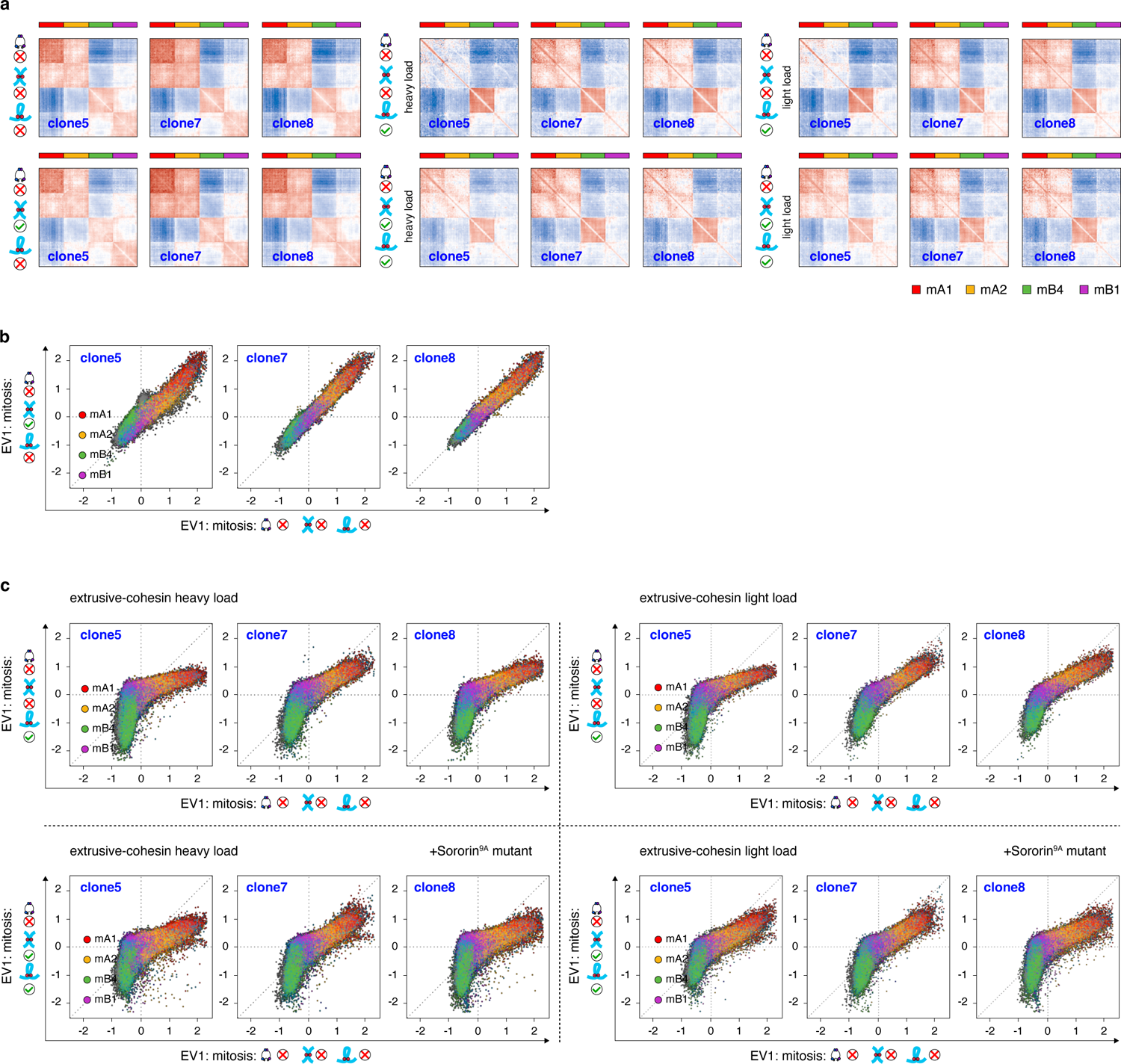
Additional data showing that extrusive-cohesin but not cohesive-cohesin modulates mitotic compartmentalization. **a**, Aggregation-repulsion plots showing the impacts of extrusive-cohesin (“heavy-” or “light-load”) or/and cohesive-cohesin on mitotic chromosome compartmentalization patterns. Plots for independent clones were shown. **b**, Scatter plots showing high correlation of EV1 values between the condensin-deficient mitotic samples without any cohesin (*x*-axis) and those with cohesive-cohesin alone (*y*-axis). Bins were color coded based on their compartment assignment. Plots for independent clones were shown. **c**, Upper panel: Scatter plots showing the reduction of EV1 values for mB4 and mA1 compartments in the condensin-deficient mitotic chromosomes loaded with extrusive-cohesin (“heavy-” or “light-load”) alone. Lower panel: Scatter plots showing the reduction of EV1 values for mB4 and mA1 compartments in the condensin-deficient mitotic chromosomes loaded with extrusive-cohesin (“heavy-” or “light-load”) and cohesive-cohesin. Plots for independent clones were shown. Bins were color coded based on their compartment assignment.

**Table S1** Hi-C statistics

**Table S2** Domain list identified by Arrowhead and insulation score profile

**Table S3** Loop list identified by HICCUPS

**Table S4** CRE contact list

**Table S5** Structural loop anchor list

**Table S6** Oligos for gene editing and genotyping

## Notes

### Competing Interest Statement

The authors have declared no competing interest.

